# Compositional Heterogeneity Structures Microbial Microhabitats across Distinct Mineral Substrates

**DOI:** 10.64898/2026.07.31.742068

**Authors:** Federica Calabrese, Peter Schroedl, Nathan Hadland, Jason Yu, Arthur McClelland, Nicholas Colella, Ryan S. Jakubek, Eric Ellison, Lisa Mayhew, Solange Duhamel, Douglas E. LaRowe, Heather V. Graham, Aaron B. Regberg, Jeffrey J. Marlow

**Affiliations:** Department of Biology, Boston University, Boston, MA, USA; Department of Organismic and Evolutionary Biology, Harvard University, Cambridge, MA, USA; Arizona Astrobiology Center, University of Arizona, Tucson, AZ, USA; J. A. Paulson School of Engineering and Applied Science, Harvard University, Cambridge, MA, USA; Center for Nanoscale Systems, Harvard University, Cambridge, MA, USA; Amentum, NASA Johnson Space Center, Houston, TX, USA; Department of Earth Science, University of Colorado Boulder, Boulder, CO, USA; Molecular and Cellular Biology, University of Arizona, Tucson, AZ, USA; Lunar and Planetary Laboratory, University of Arizona, Tucson, AZ, USA; Department of Earth Sciences, Southern Methodist University, Dallas TX, USA; NASA Goddard Space Flight Center, Greenbelt, MD, USA; NASA Johnson Space Center, Houston, TX, USA

## Abstract

Microbial communities living on and in rocks operate at the microscale, where interactions with minerals fundamentally shape community structure and function. Yet the relationship between micron scale mineralogical configurations and microbial distributions remains poorly understood. We tested the hypothesis that microbial biomass spatially correlates with areas of heightened mineralogical heterogeneity by applying Raman microspectroscopy to rock samples from three geologically distinct substrates: authigenic carbonates from a marine methane seep, volcanic basalt from Iceland, and polymetallic nodules from the abyssal seafloor. Using spectral decomposition and multiple complementary metrics of compositional heterogeneity, we evaluated intra-pixel and inter-pixel heterogeneity patterns in relation to biomass distribution. Our analyses reveal three patterns across all sample types. 1) When spectra are deconstructed into their constituent components, biomass zones are disproportionately dominated by the biomass spectral component compared with primary mineral components in zones of different minerals. 2) Biomass spectra have more homogeneous compositional profiles than mineral spectra. 3) Biomass is surrounded by more heterogeneous microhabitats than mineral pixels. These findings demonstrate that biomass exerts a distinctive and consistent influence on Raman spectral signatures, both within and between pixels, in ways that mineral components do not. Our results establish generalizable principles linking microscale mineralogical properties to microbial biogeography; these properties could be used as a potential biosignature and may provide a standardized workflow applicable to diverse rock systems and astrobiological exploration strategies.

## 1. Introduction

Understanding why organisms inhabit specific environments is one of the most foundational pursuits of ecology. Spatial relationships dictate interactions both among organisms and between organisms and their environment in ways that drive trophic dynamics and evolutionary trajectories. The associated field of biogeography is well-established for plants and animals but is poorly developed in a microbial context^1^. Microbes represent the most abundant and diverse manifestation of life on Earth, and understanding their biogeographic patterns illuminates key factors that drive their ecophysiology, such as environmental filtering, dispersal, and temporal and spatial constraints^2^.

Microbial biogeography is driven by microscale habitat dynamics: variations in mineralogy and physico- chemical properties such as surface roughness, redox state, and pH, for example, modulate microbial colonization and growth^3^. Such microscale considerations are critical because the microbial “sphere of influence” is often restricted to just several tens of micrometers, as measured by a range of modeling, culture-based, and environmental studies^4–6^. Given this range and typical cell densities of soil environments, each microbe interacts with just ∼120 neighboring cells^7^, rendering bulk reconstructions of biomolecular datasets inadequate for developing a realistic understanding of community dynamics.

Mineral assemblages can also vary on the micron scale, where multiple mineral phases can reflect a palimpsest of past episodes of precipitation, dissolution, recrystallization, and diagenesis. Patches of heterogeneity can develop as distinct minerals are incorporated into sediments and rocks, and as particular phases preferentially form or transform over time. Even within a single grain, zonation in composition, chemical reactivity, or redox gradients can affect microbial distribution and weathering processes^8^.

Because microscale spatial context constrains microbe-microbe and microbe-mineral interactions, interrogating rock-hosted microbial communities at the micron scale is critical for understanding ecological and functional relationships.

The interplay between microbes and minerals takes many forms. Most directly, minerals can provide resources, including electron donors or acceptors, as well as nutrients like phosphorus and metals^9^. Minerals can also support microbial populations by providing adsorbed organic matter. For instance, amino acids adsorb onto sulfide, oxide, and silicate minerals through electrostatic attraction and hydrogen and covalent bonding^10^. Trace metals to support enzyme function can also be sourced from minerals: nitrogen-fixing bacteria may be able to extract molybdenum (an essential cofactor for many nitrogenases) from silicates^11^, and copper availability in minerals modulates methane monooxygenase enzyme abundance in aerobic methanotrophs^12^. Independent of their chemical characteristics, minerals provide physical substrates for microbial attachment, and microtopographical details such as lattice dislocations and the orientation of crystallographic axes can influence cell recruitment^13^.

While many specific microbe-mineral associations have been detected, they lack a common framework that spans distinct microbial community compositions and mineralogical identities. Microbial colonization can be driven by mineral-specific configurations, resulting in structured diversity patterns across heterogeneous substrates^14^. Previous research revealed elevated cell concentrations and enhanced metabolic activity of iron oxidizing bacteria at rindlet-saprolite interfaces^15^, and preferential microbial colonization at the interface of igneous rocks and clay-filled fractures in South Africa^16^. However, it remains unclear how generalizable such links between biomass and mineral boundaries are.

We seek to develop such a common framework and advance the study of microbial biogeography by testing the hypothesis that microbial biomass spatially correlates positively with areas of heightened heterogeneity - that is, areas where a higher number of compositionally distinct phases are present compared with other zones. The notion that increased environmental heterogeneity offers more available niches where more organisms co-exist was developed in the context of plants and animals^17^. Further studies have shown that heterogeneity offers opportunities for specialization, diversification, community stability, and even protection across taxa and ecosystems^18^. Extending this concept to microhabitats, we propose that mineralogical heterogeneity may be a key factor in microbial habitation because different mineral phases create distinct micro-niches, and localized gradients in chemical properties could promote energy conservation and nutrient uptake.

In this study, we address these fundamental questions of microbial biogeography and microhabitat heterogeneity in the context of rock-hosted microbiomes using Raman microspectroscopy, which is sensitive to nuances in compositional heterogeneity and offers spatially resolved data through a non- destructive workflow^19,20^. Raman microspectroscopy was used to generate microscale maps of biomass and mineral classes of three geologically distinct substrates: authigenic carbonates from a marine methane seep, volcanic basalt from Iceland, and polymetallic nodules from the abyssal seafloor. We present a standardized, high-throughput workflow for spectral deconvolution that is largely agnostic to the specific mineral identities of the samples, as well as novel approaches to the quantification of both intra- and inter-pixel heterogeneity. We report that biomass contributes disproportionately to spectral composition compared with minerals (independent of the mineralogical context) and has more heterogeneous surroundings than minerals. Our findings indicate that biomass is consistently associated with distinct patterns of substrate heterogeneity across spatial scales and geologically diverse substrates; these results advance fundamental research into microbial biogeography and offer a compelling path forward in the search for life beyond Earth.

## 2. Results

Our findings reveal that biomass exerts a unique and consistent influence on Raman spectral signatures, both within and between pixels, compared with mineral components. These findings extend across geologically varied samples, which included methane seep carbonates (SC), volcanic basalts (VB), and polymetallic nodules (PN) (Fig.1).

**Fig. 1:**
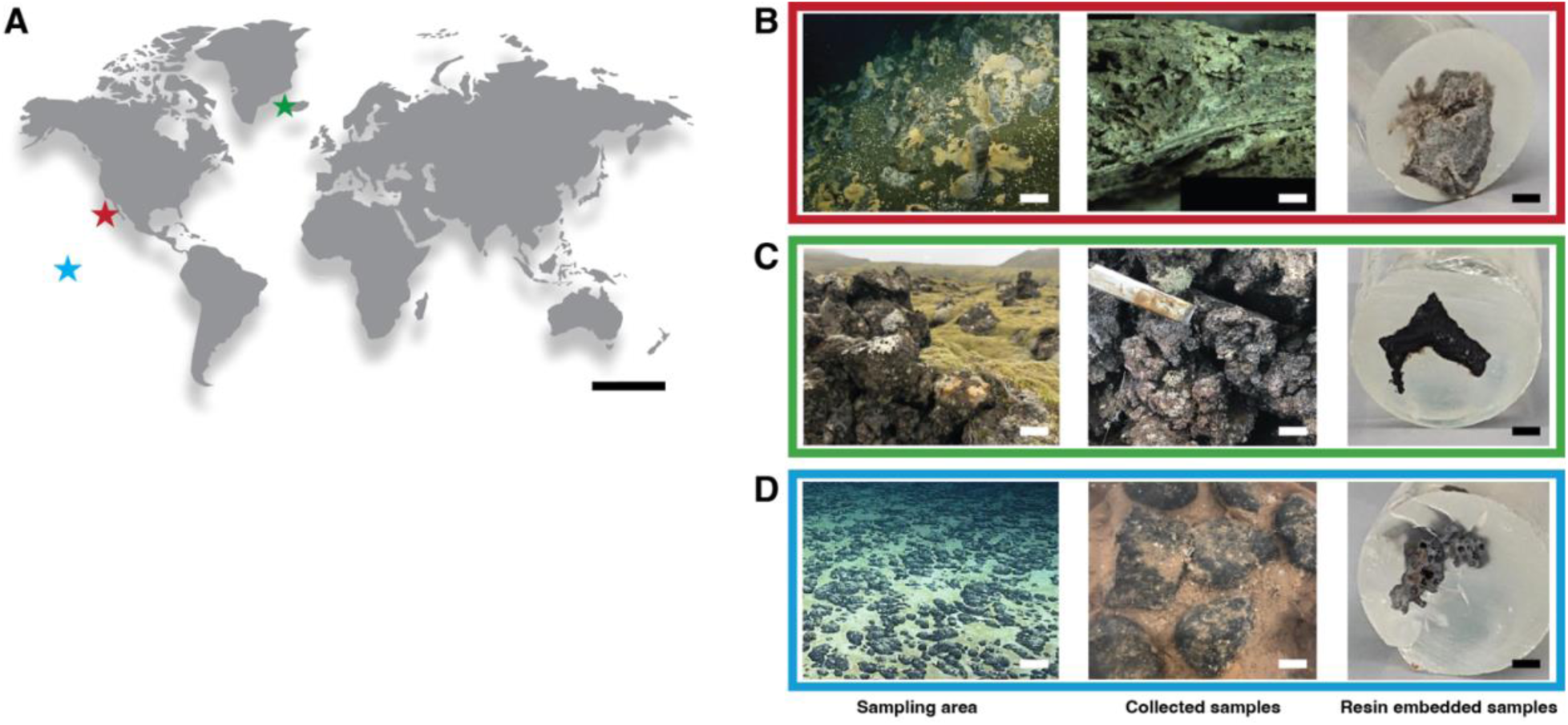
A) A global map indicating where the three samples were collected. Colored stars correspond with the boxes in panels B-D, which show samples at the landscape scale, hand sample scale, and section scale (embedded in resin). B) The Point Dume methane seep, California. C) The Fagradalsfjall volcanic complex, Iceland. D) Polymetallic nodules on the abyssal seafloor in the Clarion Clipperton Zone, Pacific Ocean. Scale bars correspond to approximately 4,000 km for the global context map; 10 cm for sampling area; 1 cm for collected samples, and 2 mm for resin embedded samples.

### 2.1. Different sample types exhibit distinct heterogeneity scores

Spectral differences can provide insights into compositional heterogeneity and inform how samples differ from one another. Overall patterns of inter-pixel heterogeneity measured with Spectral Heterogeneity Scores (SHS) were distinct among the three sample types: SC exhibited the highest median SHS value and PN the lowest (Fig. 2; Extended Fig. 1). Across all FOVs, SHS values were not distributed uniformly, but rather revealed hotspots of compositional diversity and/or patterns that track surfaces and putative precipitation fronts (Fig. 2; Extended Fig. 1). Given the agnostic nature of SHS analysis, this approach could not determine whether the observed differences arose predominantly from mineral or biomass components. Spectral decomposition was therefore applied to resolve their respective contributions.

**Fig. 2:**
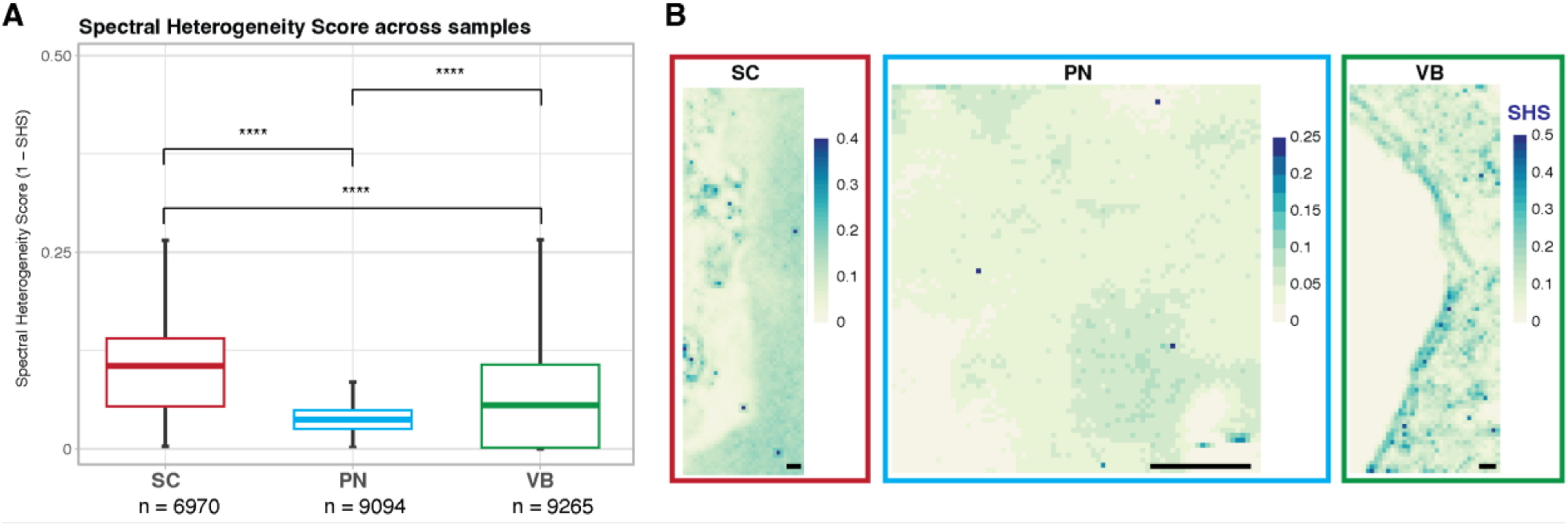
Spectral Heterogeneity Scores for each sample type. A) Boxplots of SHS values for the Seep Carbonate (SC), Polymetallic Nodule (PN), Volcanic Basalt (VB), as measured across all FOVs. B) One representative FOV is shown as a heatmap for each sample (the rest are shown in Extended Fig. 1). Color scale indicates SHS values, ranging from low (white) to high (blue). P values from the Wilcoxon test are reported as * = p < 0.05; ** = p < 0.01; ***= p < 0.001; **** = p <0.0001. On the x axis, *n* indicates the total number of pixels analyzed for each sample. Black scale bars correspond to 20 µm.

### 2.2. Mineral and biomass components were identified from spectral decomposition

Decomposition of Raman spectra using Non-negative Matrix Factorization (NMF) successfully resolved spectral components corresponding to biomass and minerals in the analyzed samples. The SC, VB, and PN samples had seven, nine, and nine components each that we assigned to reference spectra using a cosine similarity values threshold of 0.7, Table S1, Data Set 1). For the SC sample, two additional components, a London Resin White (LR White) resin spectrum and a biomass spectrum, were also included in the analysis despite lower cosine similarity scores since they provided the best matches to known constituents of the sample. Biomass spectra in particular are typically noisier than pure mineral phases, which effectively decreases their cosine similarity scores.

The list of components for each sample, along with the percentage of pixels for which they are the top match, is shown in Table S1. In most cases, we found that several of the top “hits” for each component, as measured by cosine similarity scores, were of different RRUFF database spectra of the same mineral or mineral class (Data Set 1). This consistency, as well as the observation that most of the identified minerals had previously been identified as a constituent of the host rock based on published analyses, often using X-ray diffraction data, provided confidence in the IDs (Supplementary Information). In cases where the specific mineral identified as a component is not traditionally associated with the analyzed rock, the more generalized “class” of mineral, as determined by the Dana classification system, is^21^. For this reason, we map and interpret these broader classes of minerals (e.g., carbonates, sulfides, various types of silicates) throughout the study. This degree of mineralogical resolution is comparable to ecological analyses of microbial community composition, where broader taxonomic groupings (e.g., at the family or phylum level) are often preferred over species- or strain-level analyses in order to derive more general ecophysiological trends. We also manually compared each high-similarity component’s spectrum with the corresponding experimental spectrum to ensure that relevant molecular spectral features were present (Supplementary Information; Fig. S1-S3), bolstering the class-level identifications.

#### 2.2.1. Biomass Verification

Biomass was observed in each Field of View (FOV) of all three rock samples using fluorescence microscopy and general DNA stains (Fig. S4). While our confocal fluorescence and Raman microspectroscopy instruments had different optics and resolutions, making direct, pixel-scale correlations challenging, the confirmation of biomass offered confidence that our downstream detection of biomass-associated Raman spectral components accurately captured biological materials. In this context, we used the fluorescence signal primarily to identify areas where microbial biomass was concentrated, thereby facilitating Raman-based assessments of spatial configurations and spectral properties.

#### 2.2.2. Biomass and Resin Components

In each of the three sample types, three non-mineral components were included in the analysis: biomass, kerogen, and resin. “Fresh” biomass from living organisms typically shows distinct but low-intensity peaks at ∼1230-1290 cm^-1^ (amide III from proteins), ∼1450 cm^-1^ (CH_2_ stretch usually found in lipids and proteins), ∼1123-1129 cm^-1^ (C-N and C-C stretches) and ∼1650-1680 (amide I and unsaturated lipids)^22^. Other features characteristic of biomass are peaks at ∼780 cm^-1^ (nucleotides U, C, T), ∼1008 cm^-1^ (Phenylalanine), and ∼1660 cm^-1^ (amide I from proteins)^23^. Interpretation of biomass spectra is complicated by the fact that the position and size of Raman peaks are sensitive to changes in laser wavelength, fluorescence background, the degree of analyte hydration, and the incidence of the laser on the sample. The identification of biomass with Raman thus relies on the presence of multiple peaks corresponding to proteins, lipids, or nucleic acids.

London Resin White (LR White) is an acrylic resin used for embedding biological and mineralogical specimens and was well-suited to our applications because of its low viscosity, high hydrophilicity, and beam stability^24^. Its primary Raman features are found at ∼1725–1735 cm^-1^ (carbonyl group C=O), 1640 cm^-1^ (C=C stretches), ∼1250–1300 cm^-1^ (C–O stretching mode), and sharp peak at ∼2950–3000 cm^-1^ (C-H_2_ and C-H_3_ stretching)^25^.

#### 2.2.3. Components of the Rock Samples

The most abundant mineralogical component in sample SC was the carbonate mineral aragonite, followed by vesuvianite, a sorosilicate, and kutnohorite, another carbonate (> 9% of the composition, Table S1).

The sulfide mineral class was the top component in nearly 5% of the pixels, while oxide/hydroxide (3.7%) and tectosilicate (2.1%) represented a small percentage of the minerals. Biomass and kerogen components together represented ∼40% of the total pixels, consistent with the dense chemosynthetic communities that inhabit seep carbonates.

In the VB sample, the most abundant mineralogical component was the inosilicate ferrosilite (orthopyroxene), which accounted for 8% of the composition. Other prominent mineral classes included oxide/hydroxide (6.7%), sulfide (4.8%), and other types of silicates (sorosilicates, phyllosilicates, and tectosilicates, all < 4.5 %), including plagioclase feldspar mineral. Resin accounted for nearly 40% of the pixels; since the resin replaces air during embedding, this observation likely reflects the porous nature of volcanic rock.

The four most abundant mineralogical components in our PN sample were oxide/hydroxide minerals (constituting a combined ∼55% of the analyzed area), followed by tectosilicates (1.3%) and phyllosilicates (0.9%) in small proportions. Of the three samples, PN had the lowest biomass abundance.

#### 2.2.4. The Kerogen Component

Kerogen characteristic Raman peaks, usually referred to as D (∼1350 cm^-1^) and G (∼1580 cm^-1^) bands, are often used to assess the thermal maturity of carbonaceous microfossils^26^. In the early stage of maturation, as kerogen undergoes increasing thermal alteration, these bands are relatively broad^27^, reflecting a highly disordered molecular composition. As kerogen thermal maturity increases, structural order increases, bands become narrower, and the peaks separate. Raman is thus able to capture the transition of “fresh” biomass into kerogen. During Raman analysis, fresh organics are susceptible to carbonization due to laser-induced photodegradation. The spectra identified as kerogen in all the three of our samples showed broad D and G bands as well as minimal peak separation (Supplementary Information; Fig. S1-S3), suggesting an immature stage of alteration^27^. In this study, we combined matches of “fresh biomass” and kerogen into the “biomass” component because of previous evidence that incident laser light can alter even intact cell material into organic assemblages that yield best spectral matches to kerogen^28^, as well as the broad peaks of the kerogen components we detected from microbial cells (Supplementary Information; Fig. S1-S3).

### 2.3. Pixel decomposition allowed to reveal microbe-mineral associations

Focusing on the group of pixels identified as biomass, we determined which mineral components preferentially co-localized with biomass for each of the three sample types, informing potential microbe-mineral interactions (Fig. 3 A, C and E). Each pixel is made up of a primary component - the component with the highest weight after NMF - and additional components with smaller compositional contributions. We grouped pixels by their primary component, and quantified secondary components to reveal the most prevalent co-occurrences (Fig. 3).

**Fig. 3.**
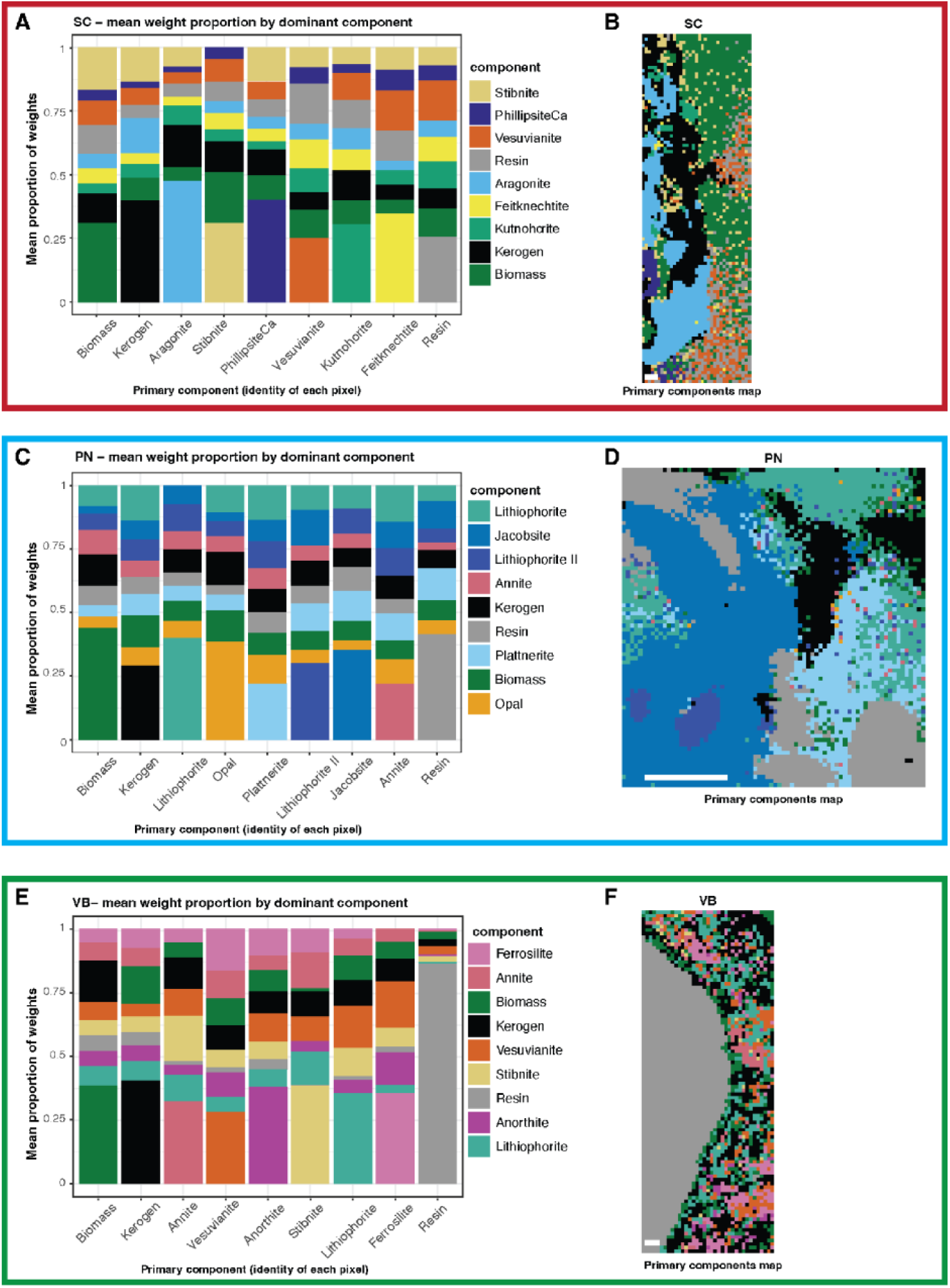
Pixel identity and decomposition as determined by Non-negative Matrix Factorization (NMF) analysis. A), C), E) Stacked bar plots showing the average component composition of pixels assigned to each identity class in the three samples: Seep Carbonate (SC), Polymetallic Nodule (PN), Volcanic Basalt (VB), respectively. Pixels are grouped according to their dominant component identity. In each stacked bar, the bottom segment represents the dominant (primary) component used to assign pixel identity, whereas the upper segments represent the average contributions of the remaining (minor) components. The y-axis indicates the mean component weight across all pixels within each identity class, and the x-axis lists the identified components for each sample. B), D), F) Representative FOVs for samples SC, PN and VB, respectively. Each pixel is colored according to its assigned identity, defined as the component with the highest NMF weight, using the same color scheme as the corresponding stacked bar plots.

For the SC sample, biomass was most associated with the sulfide mineral class, followed by silicates. Kerogen was most associated with carbonates (aragonite), followed by sulfides (Fig. 3A). In the PN sample, biomass and kerogen were primarily associated with Mn oxide/hydroxide minerals such as lithiophorite and jacobsite (Fig. 3C). Both biomass and kerogen pixels in VB showed a relatively even distribution of subsidiary components, most of which were silicate minerals (anorthite, ferrosilite, and vesuvianite) (Fig. 3E).

### 2.4. Intra-pixel heterogeneity measures show consistent differences between biomass and minerals

Intra-pixel compositional heterogeneity was computed using the Gini and Shannon indices. We found that pixels identified as biomass were less compositionally heterogeneous compared with mineral pixels, a trend that was consistent across the three samples and showed a small yet significant effect size (Fig. S5, Data Set 2).

Gini index values were plotted for each sample type and grouped by pixels identified as biomass and those identified as minerals (Fig. 4). Higher values correspond to a pixel where all components are more equally distributed and thus have a more heterogeneous composition. The resin component was omitted from these analyses in order to only capture naturally occurring features. To determine how this pattern was shaped by the presence of biomass itself, we re-computed the Gini index by removing the dominant components from all pixels (Fig. 4B). Following this adjustment, pixels identified as biomass exhibited a significant increase in compositional heterogeneity while mineral pixels did not. Notably, since the removal of a mixture’s most prevalent component inherently raises its “1-Gini” value (by decreasing compositional inequality), the shift in the index values when biomass was removed was substantially more pronounced than when the top component was removed from mineral pixels (Fig. 4, “no dominant” boxes, Data Set 2). Taken together, these analyses indicate that, across all sample types, biomass is a more dominant component of “biomass” pixels than primary mineral components are in their respective “mineral” pixels.

**Fig. 4:**
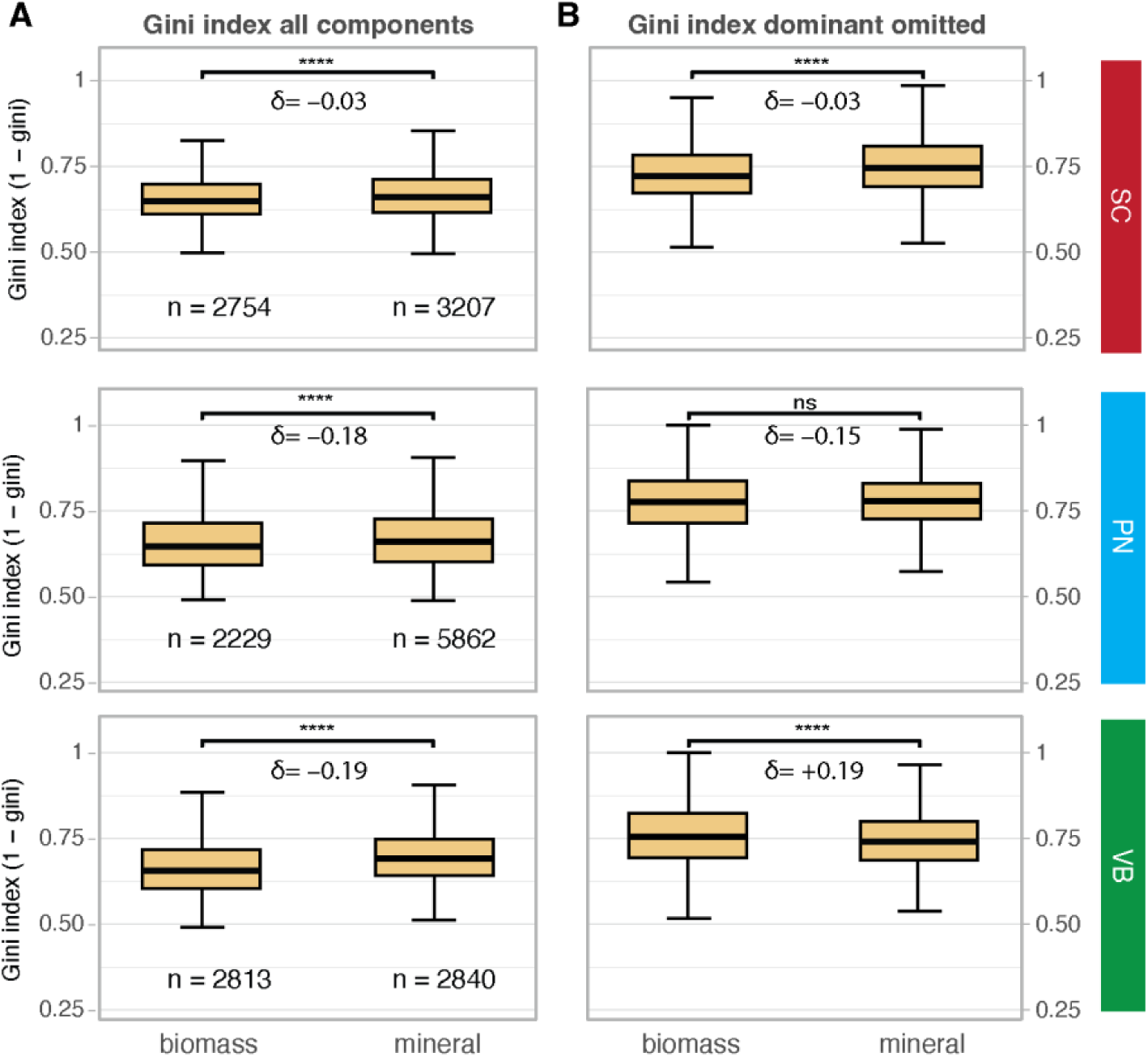
Gini index plots for biomass and mineral pixels in the Seep Carbonate (SC), Polymetallic Nodule (PN), and Volcanic Basalt (VB) samples showing inequality of component distributions across samples and pixels type. A) Data is plotted when all components are considered, and B) when the dominant (primary) component of each pixel is omitted. Cliff’s delta values (δ) indicate the value of the effect size for each comparison biomass vs. mineral. Negative signs indicate that biomass scores are lower. The *n* values indicate the number of pixels for each category. Note that resin pixels have been omitted from the calculations.

In all three samples, Shannon index values were higher for mineral pixels than biomass pixels, indicating that mineral pixels have more heterogeneous compositions (Fig. 5). Median Shannon values were 1.586 and 1.529 for minerals and biomass in SC (Cliff’s delta = −0.05), 1.653 and 1.51 for minerals and biomass in PN (Cliff’s delta = −0.17), and 1.546 and 1.43 for minerals and biomass in VB (Cliff’s delta = −0.22) (Fig. 5A). Removing the primary (top-weighted) component from each pixel did not change this trend: median Shannon values were 1.647 and 1.562 for minerals and biomass in SC (Cliff’s delta = −0.04), 1.717 and 1.579 for minerals and biomass in PN (Cliff’s delta = −0.16), and 1.422 and 1.393 for minerals and biomass in VB (Cliff’s delta = −0.04) (Fig. 5B). These results show that the compositional heterogeneity is not driven by dominant components in the mineral pixels but is rather a more ingrained property reflected by the full range of components in biomass and mineral mixtures.

**Fig. 5:**
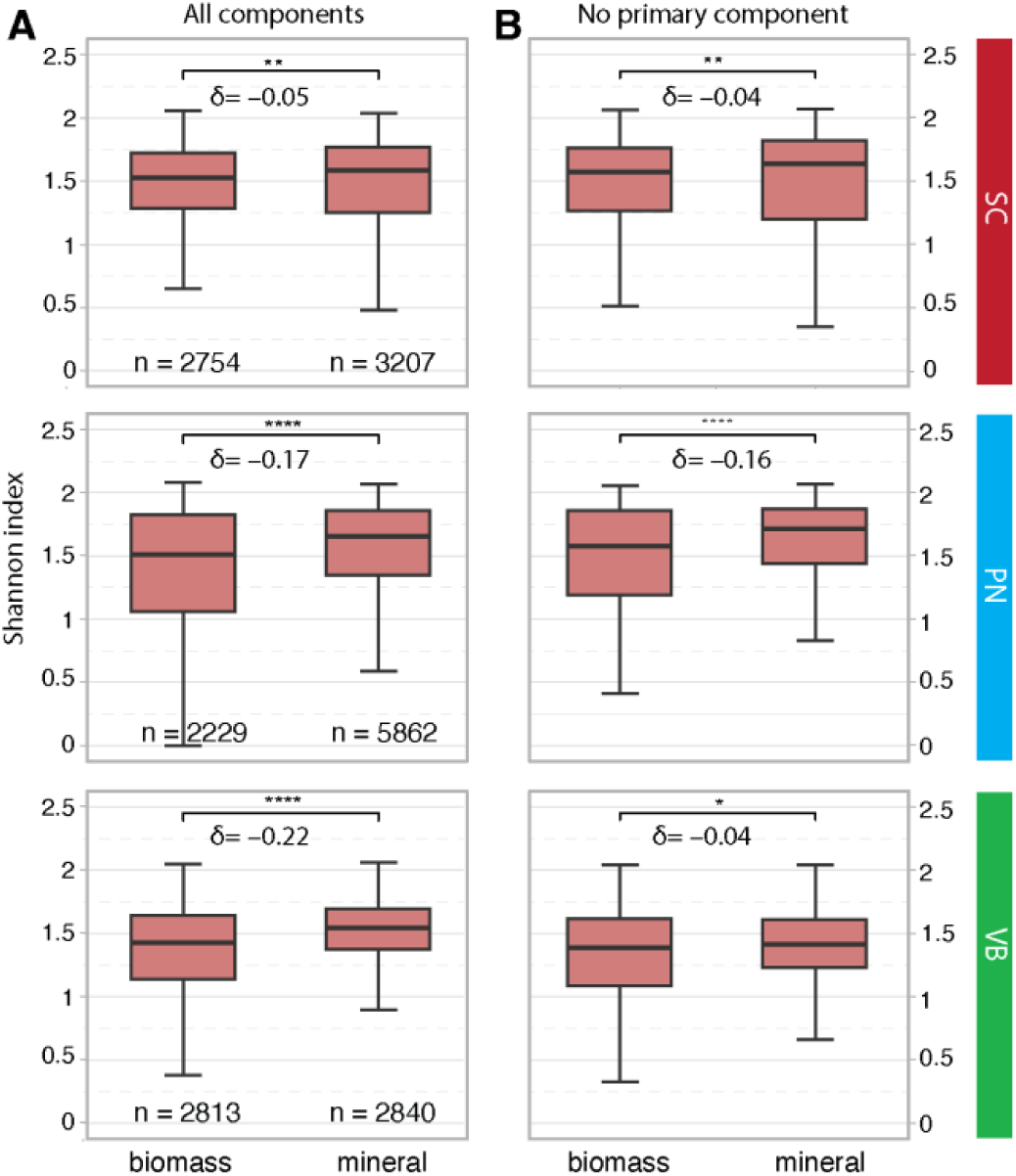
Shannon index plots for biomass and mineral pixels in the Seep Carbonate (SC), Polymetallic Nodule (PN), and Volcanic Basalt (VB) samples. A) Boxplots of pixel values when all components are considered, and B) when each pixel’s dominant component is omitted. Cliff’s delta values (δ) indicate the value of the effect size for each comparison of biomass vs. mineral, where negative signs indicate that biomass scores are lower. The n values indicate the number of pixels for each category. Note that resin pixels have been omitted from the calculation and biomass include both biomass and kerogen pixels. P values from the Wilcoxon test are reported as * = p < 0.05; ** = p < 0.01; ***= p < 0.001; **** = p <0.0001.

### 2.5. Inter-pixel heterogeneity measures indicate that biomass is found in heterogeneous microhabitats

To evaluate potential links between biomass and its surroundings, we used two independent measures of inter-pixel heterogeneity: the compositionally agnostic SHS and the spectral components-based Rao’s Q index. SHS analyses showed that the spectral surroundings of biomass and mineral pixels were statistically distinct. For SC rock, biomass “neighborhoods” were more homogenous than those of mineral pixels, while for PN and VB samples, this relationship was inverted and biomass pixels had more heterogeneous neighboring pixels than minerals (Fig. 6). These patterns were also confirmed by Cliff’s delta values.

**Fig. 6:**
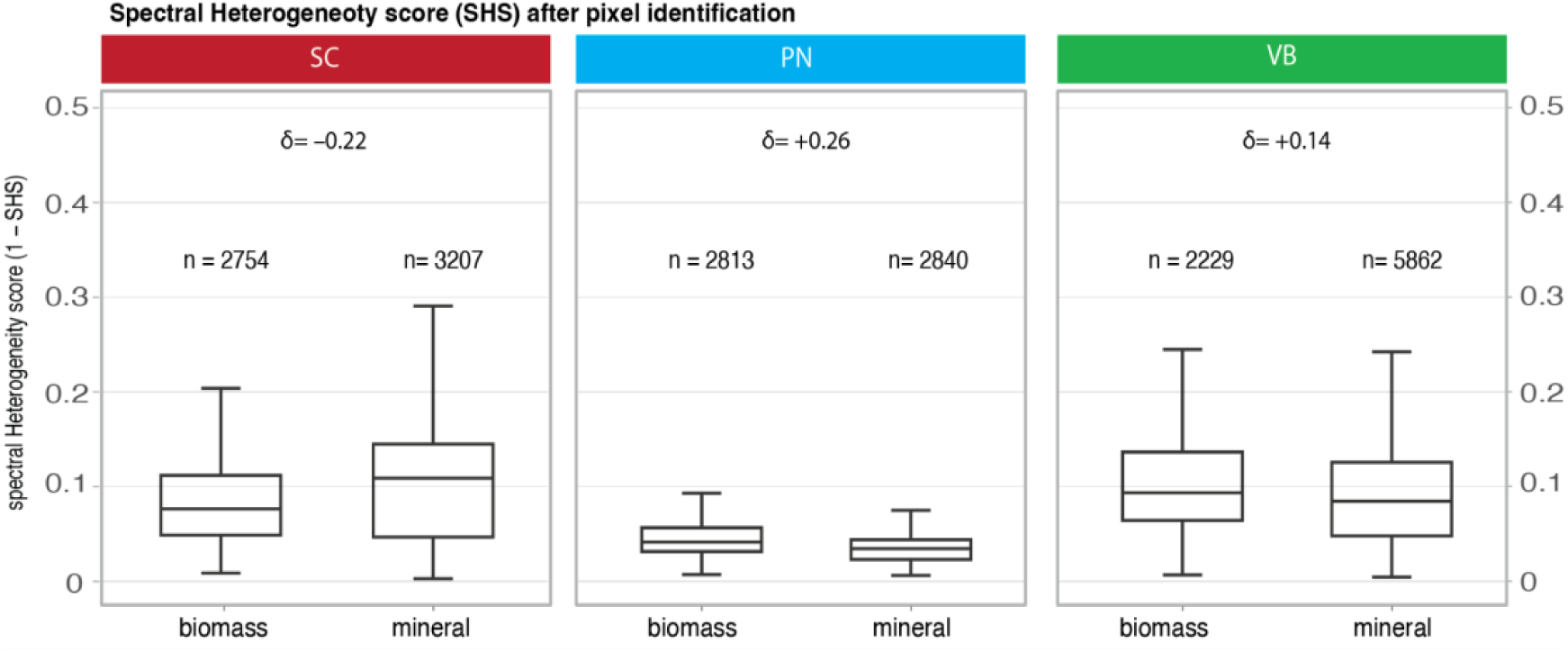
Spectral Heterogeneity Score (SHS) plots for biomass and mineral pixels in the Seep Carbonate (SC), Polymetallic Nodule (PN), and Volcanic Basalt (VB) samples. Cliff’s delta values (δ) are reported for each comparison of biomass vs. mineral pixels. Negative signs indicate that biomass scores are lower. The n values indicate the number of pixels for each category.

Rao’s Q calculations indicated that, when compared with their surroundings, biomass pixels have a higher compositional heterogeneity than mineral pixels when all the components are included in the calculation. This trend was consistent across the three sample types (Fig. 7A). When the dominant component of each pixel was removed (Fig. 7B), biomass pixels still displayed consistently higher Rao’s Q values.

**Fig. 7:**
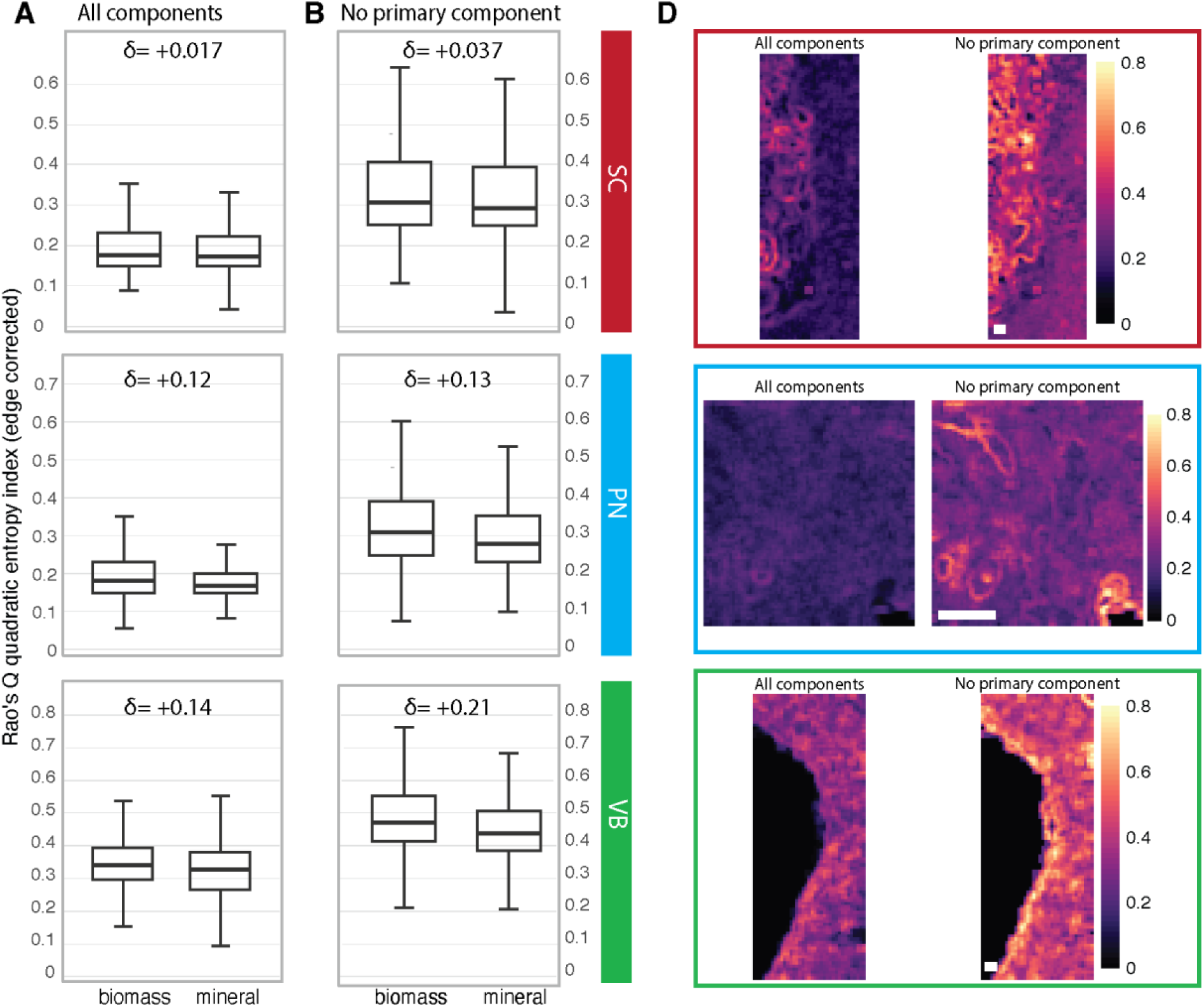
Rao’s Q values for biomass and mineral pixels in the SC, PN, and VB samples A) when all components are considered, and B) when the dominant/primary component of all pixels was removed. Cliff’s delta values (δ) are shown for each comparison. C) For each sample, Rao’s Q maps for a representative FOV are shown enclosed in boxes with the same heat map scale representing the magnitude of the Rao’s Q values. Rao’s Q maps for the remaining FOVs can be found in Extended Fig. 2. Scale bar is 20 µm.

Because Rao’s Q and SHS are computed from each pixel’s neighbors, adjacent pixels might be correlated by default, violating the independence assumption of the Wilcoxon rank-sum test. For these two metrics we thus report the effect sizes instead of the p value (Data Set 2).

To account for spatial autocorrelation, repeated thinning of the data was performed (Table S2). The analysis revealed that the observed effects on intra-pixel contrasts (Gini and Shannon) and inter-pixel metric (SHS and Rao’s Q) are consistent and are not artifacts of spatial proximity. Taken together, our data show that biomass pixels retain greater intrinsic compositional diversity in relation to their surroundings - even when looking past the primary component for each pixel - compared with mineral pixels.

### 2.6. Distinct microbial communities inhabit seep carbonate, polymetallic nodule, and volcanic basalt samples

The microbial communities associated with the different sample types were compositionally distinct, and the dominant microbial families were consistent with previously described metabolic interactions between these taxa and the minerals we detected.

In the SC sample, families associated with sulfate reduction accounted for three of the six most abundant families: *Desulfobacteraceae*, *Desulfobulbaceae*, and *Desulfuromonadaceae* made up 13%, 10.6%, and 3.6% of the community, respectively. (*Desulfobulbaceae* also includes sulfide-oxidizing cable bacteria^83^, though no known cable bacteria genera were detected, Data Set 3.) The aerobic methanotrophic bacteria *Methylomonaceae* accounted for 11.9% of the community. Two bacterial families of primarily (facultatively) anaerobic heterotrophs were also abundant: *Marinifilaceae* (4.9%) and Clostridiales Family XII (4.1%). ANME were relatively scarce, accounting for just 2.5% of the sequences. Of these, more than 90% were ANME-1, which are the most phylogenetically divergent of the ANME clades^84^. The remaining ANME sequences belonged to the ANME-2a-2b subgroup, which are the most common lineage at marine cold seeps^85^.

In the PN microbial community, archaeal *Nitrosopumilaceae* accounted for 17.7% of the sequences (Data Set 3), and were three times more abundant than the next most prevalent family, the *Woeseiaceae* (5.7%). Other taxa potentially involved in biomineralization and metal cycling were present in the sequenced community, such as *Magnetospiraceae* (2.6%), *Nitrosomonadaceae* (0.8%), and *Nitrosococcaceae* (0.8% each)^86–89^. *Shewanella* and *Colwellia* were found at low relative abundances (0.1% each).

The Fagradalsfjall basalt-hosted community was characterized primarily by heterotrophic taxa (Data Set 3). *Chthoniobacteraceae*, a family within the *Verrucomicrobia*, accounted for 17.3% of the sequences, *Solibacteraceae* represented 13.1% of the community, and *Acetobacteraceae* 12.2%.

## 3. Discussion

Understanding where organisms live at scales relevant to the physiological interactions they have with their environment is a foundational objective of ecology. For microbial communities inhabiting rocks — which may constitute the largest microbiome on Earth — this process is particularly challenging given the small spatial scales involved and the difficulty of distinguishing microbes from minerals with traditional microscopic techniques. Here, we leveraged the spectral “fingerprinting” capabilities of Raman microspectroscopy to evaluate links between rock compositional heterogeneity and the presence of biomass. We used multiple metrics of intra- and inter-pixel heterogeneity to derive common properties across a range of geological / mineralogical contexts and microbial community compositions.

In all but one of the relationships detailed above, measures of intra- and inter-pixel heterogeneity showed a consistent trend in differences between pixels identified as biomass and those identified as minerals.

Furthermore, in nearly all cases, the trends were consistent across all three sample types; given that these substrates represent very different geological contexts and microbial communities, the patterns we report may be generalizable properties of rock-hosted life. These patterns can be summarized as follows. 1) Biomass as a primary component had a more pronounced impact on the compositional heterogeneity of biomass pixels than mineral primary components did on mineral pixels. 2) Biomass pixels had more unequal compositions compared with mineral pixels. 3) Biomass had more heterogeneous microscale surroundings than minerals. These findings indicate that biomass influences the Raman spectral signatures, both within and between pixels, in ways that minerals do not.

By identifying mineral components and assessing microbial community composition, we can interrogate putative microbe-mineral relationships across each sample type. Authigenic carbonates from the Point Dume methane seep exhibit millimeter-scale variations in carbonate polymorphs, elemental abundances, and sulfur minerals, which likely reflect changes in methane flux and mineral precipitation over time^28^. The prevalence of biologically induced carbonate and sulfide minerals at methane seeps is well established^29,30^, and can exhibit spatially heterogeneous patterns based on localized variations in fluid chemistry and microbial activity^31^. Seep carbonates also incorporate detrital sediment grains such as silicates^32^, particularly when seep sites are close to shore^33^. These direct links between microbial processes and a range of minerals may explain our observation that SC had the highest median SHS.

Microbial community analysis can be used in concert with biomass-mineral associations from Raman spectral decomposition to illuminate putative links between metabolism and mineral distributions. In the SC sample, biomass was primarily associated with the sulfide, carbonate, and silicate mineral classes, and the microbial community included prevalent SRB and aerobic methanotrophs. SRB have been shown to induce authigenic carbonate precipitation by increasing alkalinity through their metabolic activity^34^, and by serving as nucleation sites for mineral formation^35^. Sulfide mineral precipitation is also enhanced in the presence of SRB, via the metabolic generation of dissolved sulfide, as well as by mineral templating on cell surfaces and extracellular polymeric substances^36^. The aerobic methanotrophic *Methylomonaceae* family is common at methane seeps^37^ and, as observed at a different region of seepage off the California coast (Coal Oil Point), may be transported from the benthos to the water column by gas bubbles^38^. We also detected sequences from ANME Archaea, which form syntrophic relationships with SRB to perform the anaerobic oxidation of methane^39^. The Point Dume seep carbonates were previously shown to possess abundant and metabolically active ANME-SRB consortia^40^, which induce authigenic carbonate precipitation by producing bicarbonate and increasing localized alkalinity levels^29^. These consortia have been observed in direct physical association with framboidal pyrites (a sulfide mineral)^40^, and can promote localized silica precipitation^41^. Nonetheless, the relatively low proportion of ANME sequences was surprising; it is possible that AOM from earlier stages of the community’s development facilitated the precipitation of authigenic carbonate and silicate mineral phases, and that the continuing dominance of SRB enhanced carbonate and sulfide mineral formation.

Polymetallic nodules form over million-year timescales through the precipitation of metal oxides on or just beneath the seafloor^42^, and they frequently have concentric Fe–Mn (oxy)hydroxide layers that record shifts in hydrogenetic and diagenetic growth processes^43^. Hydrogenetic layers, which accumulate elements from water column seawater, are relatively uniform and broadly enriched in Co, Si, Cl, and S. Diagenetic layers, which source metals from sediment porewater, are more variable and retain higher concentrations of Mn, Ni and Cu^43,44^. However, since nodule laminae are mm-scale features, each of our FOVs was restricted to a single layer, which may account in part for our observation that PN analyses had the lowest SHS median value.

Biomass was most commonly associated with Mn oxide / hydroxide minerals, which is consistent with scanning electron microscopy analysis that showed endolithic microorganisms primarily associated with Mn-rich layers^45^. *Nitrosopumilaceae*, which was the most abundant family in our sequence data, is often found within nodules and their associated sediments across the CCZ^46^, and is typically associated with nitrogen cycling and ammonium oxidation^47^. Members of this family may also facilitate iron and manganese-based metabolisms that aid in the formation of the Mn oxides comprising the nodules themselves^48^. The magnetotactic bacteria *Magnetospiraceae* (2.6%) may mobilize iron from the prevalent metal oxide minerals we detected with Raman, contributing to the estimated 18-77% of nodule volume composed of microbially derived minerals^49^. Blöthe et al. have proposed an internal cycle of manganese oxidation and reduction^50^, which could position microbes at mineral interfaces where heterogeneity may be elevated, consistent with our results. This putative cycle was anchored by *Shewanella* and *Colwellia*, which we find at low relative abundances in our sample, though uncharacterized taxa could play similar roles.

Petrological and geochemical studies of Fagradalsfjall tholeiitic basalt demonstrate that significant mineralogical heterogeneity can develop within a single volcanic system based on magma evolution and localized crystallization processes both before and after eruption^51^. Nonetheless, lava flows from individual eruptive events on the Reykjanes Peninsula exhibited remarkable chemical consistency, indicating that mixing likely occurred prior to extensive crystallization within the crust, leading to compositionally homogeneous basalt^52^. This process may be reflected in the relatively low overall compositional heterogeneity we found in the VB sample.

Biomass in the VB sample was primarily associated with silicates. Experimental and field-based mineral weathering studies have shown that microbes interact with silicate minerals through the production of organic ligands, enhancing mineral dissolution and facilitating the release of biologically relevant elements (e.g., Ca, Na, K, Al)^53^. The VB microbial community was characterized largely by heterotrophs with metabolic adaptations that may facilitate their survival within the basaltic rocks.

*Chthoniobacteraceae* have the capacity for fermentation and nitrate reduction; they encode xylanase enzymes that accelerate the degradation of complex organics^54^, and may have experienced streamlining selection to enhance fitness in nutrient-limited environments^55^. *Solibacteraceae* have a proclivity for phosphorus acquisition from otherwise nutrient-limited soils^56^, and while this phylum (*Acidobacteria*) has primarily been characterized as heterotrophs, some members encode CO dehydrogenase, which could allow for autotrophy in environments with limited organic carbon^57^. *Acetobacteraceae* are common in arid environments including airborne dust^58^, and encode an array of iron binding and copper chelating molecules^59^ that allow some taxa to grow chemolithotrophically via iron oxidation^60^.

*Solirubrobacteraceae* have been observed in silicate-weathering communities inhabiting Icelandic volcanic basalt^61^, and were positively correlated with Ca^62^, which is a prominent cation in silicate minerals we detected with Raman. Despite these putative associations, direct links between microbial metabolism and mineral constituents – particularly via biomineralization – are not as well established in basalts as in authigenic carbonates at seeps or in polymetallic nodules.

The detection of kerogen in all of our analyzed samples – including exponential phase pure cell cultures – suggests that caution is necessary when interpreting Raman spectra designated as biomass or kerogen, particularly within mineralogical matrices. In this context, susceptibility to laser damage changes substantially from sample to sample and spot to spot due to variations in overcharging or heating. As a result, cell or organic molecule damage can cause otherwise “modern”, intact biomass to be detected as kerogen-like peaks.

Looking beyond specific minerals and microbial taxa, we evaluated potential patterns in compositional heterogeneity and assessed both intra-pixel heterogeneity, which examined the components constituting each individual pixel, and inter-pixel heterogeneity, which considered diversity across pixels. Gini index data showed that biomass pixels were less heterogeneous than mineral pixels. We also found that the biomass component exerted a stronger influence on pixels identified as biomass than top mineral components did on their respective pixels. The Shannon index, which incorporates measures of a distribution’s richness and evenness, showed that mineral pixels were more heterogeneous than biomass pixels across all three sample types. Even when each pixel’s dominant component was removed, minerals maintained higher median Shannon index values, indicating that the observed diversity differences were resilient, to some extent, to increased evenness, and were not solely attributable to the most abundant component.

The consistently higher Gini and Shannon index values observed in the mineral pixels indicate that they maintain both greater component diversity and a more even distribution of component weights than the biomass pixels. Together, these patterns suggest that minerals exhibit a more structurally heterogeneous assemblage, with relative compositional weights distributed across a broader range of components compared with biomass zones. However, after excluding the dominant component, the biomass pixels exhibited higher Gini index values compared with mineral pixels despite maintaining lower Shannon entropy. This result indicates that the lower evenness observed in the biomass pixels was primarily driven by the dominant species, whereas the remaining components were comparatively more evenly distributed. In contrast, the mineral pixels retained higher component diversity (via Shannon) after dominant components were removed, suggesting that their greater heterogeneity is attributable to a more broadly varied composition rather than being shaped by the influence of one highly abundant component.

To explore inter-pixel relationships, we used Rao’s Quadratic Entropy (Rao’s Q), which incorporates both abundance and pairwise differences among components between neighboring pixels. Rao’s Q is frequently applied to measure structural and spatial heterogeneity, where a higher Rao’s Q indicates a more varied, complex, and heterogeneous landscape^63^. Biomass pixels showed higher Rao’s Q values than mineral pixels, indicating higher heterogeneity between a given biomass pixel and its surroundings. The persistence of higher Rao’s Q values in biomass pixels after removal of the dominant component and the increase in the magnitude of this difference (Fig. 7), shows that the higher heterogeneity of biomass pixels is not driven by the biomass alone. Rather, it reflects an underlying mineralogical complexity based on components that are intrinsically more distinct from one another, with the contribution of these differences becoming more pronounced once the influence of the dominant component is removed (Fig. 7C). In short, biomass pixels are more distinct from their surrounding pixels than mineral pixels are, largely based on non-biomass components. This finding suggests that cells either preferentially colonize or create microhabitats of heightened mineralogical heterogeneity.

SHS calculations also revealed a higher degree of heterogeneity between biomass pixels and their surroundings than between mineral pixels and their surroundings (Fig. 6). (Note that SHS compares full spectra, but we were able to assign pixel “identities” from NMF and spectral decomposition.) The only exception to this trend was for sample SC, where biomass pixels’ SHS median value was lower than that of mineral pixels. This result, also confirmed by the effect size analysis, may reflect the biomass-induced precipitation of carbonate, which could generate a relatively homogeneous mineral covering immediately around cells^64^. Because SHS values are mineralogically agnostic and depend solely on pairwise comparisons between the spectra themselves, the metric may be particularly useful for detecting compositional diversity that may not manifest as distinct mineral phases.

Given the “snapshot” nature of our samples, which capture a single moment in time, it is unclear how long the biomass detected in our survey had inhabited each rock at the time of collection, and different scenarios lead to different interpretations. If we captured a relatively recent community, or one that did not appreciably affect its mineralogical surroundings, then our results may indicate that microbes preferentially settle on sites of lower compositional heterogeneity such as specific minerals that adsorb nutrients or provide electron donor / acceptor opportunities. Cases of heightened inter-pixel heterogeneity metrics associated with biomass (VB and PN) further suggest that biomass occurs as single cells or small colonies / communities, and that extensive biofilms covering many pixels (tens of microns) are rare, as such situations would result in low inter-pixel SHS and Rao’s Q values. On the other hand, if we sampled a more established community that had shaped the substrate on which it settled, e.g., through biomineralization, dissolution, and/or redox transformations, then our observations may reflect the time-integrated effects of biological activity. For example, by converting methane carbon into calcium carbonate minerals, ANME-SRB in the SC sample may be homogenizing the mineralogical composition of both their immediate footprint and their surroundings. The best way to distinguish between these scenarios would be to track the progression of biomass distribution and mineralogical composition of rock-hosted microbial communities over time, from a sterile surface to a mature geomicrobiological habitat. Observing changes over time will clarify the degree to which microbial presence and activity transforms mineralogy and, more broadly, constructs or degrades zones of mineralogical heterogeneity.

We anticipate future implementations of this technique may incorporate molecular tools and incubation experiments to derive more detailed ecophysiological insights linking metabolic activity, microbial identity, and spatial complexity. For example, (dual) BONCAT^65^, Raman-stable isotope probing^66^, and Fluorescence *in situ* Hybridization^67^ could be used to precisely locate and identify anabolically active cells. Such information will be critical for revealing specific microbe-microbe and microbe-mineral interactions that are favored by particular mineralogical configurations and identities.

Our finding that higher spatial heterogeneity is associated with a greater abundance and diversity of organisms aligns with previous work in the context of macroorganisms^18^. Spatial environmental heterogeneity may thus be a general parameter that extends across different levels of biological organization. The fundamental principles of microbes’ spatial relationships with their microscale surroundings can also inform astrobiology exploration strategies. For example, Raman systems aboard the *Perseverance* Mars rover could be used to identify microscale zones of decreased compositional heterogeneity (as measured by the Shannon index) whose dominant component contributes disproportionately to Gini coefficient values. Such samples might then be prioritized for more detailed astrobiological study, including through sample return, to possibly detect signatures of extinct life.

By establishing that microbial biomass exhibits consistent spatial heterogeneity patterns across geologically distinct substrates, we provide a generalizable framework for pinpointing microbial life in complex natural environments. These findings offer a new window into microbial biogeography and provide powerful tools for searching for life on Earth and beyond.

## 4. Methods

### 4.1. Sample Context & Collection

In order to illuminate general principles linking mineralogical configurations with microbial abundance in endolithic habitats, we used rocks from three distinct geological contexts: seafloor methane seeps, a terrestrial volcanic field, and the deep-sea abyssal plain (Fig. 1).

Methane seep carbonate rocks (“SC” hereafter, for “seep carbonates”) were collected from the Point Dume seeps (720-730 m water depth, 5.4 °C) off the coast of southern California, during R/V *Falkor* leg FK181005 in October 2018. The site is positioned at the base of the Dume submarine canyon located within the northern end of the inner Continental Borderland, a marine basin bounded by active fault systems. Previous work demonstrated that the Point Dume carbonate chimneys are composed primarily of biogenic calcium carbonate precipitated over the last ∼20 kyr^32^, and that they support active microbial communities anaerobically oxidizing methane at the highest potential rates measured to date^40^. Volcanic basalt (“VB”) was collected from Fagradalsfjall complex, a shield volcano on the Reykjanes Peninsula in southwestern Iceland. The rocks used in this study are from a lava flow that erupted ∼2 kya, consisting of basaltic rocks made mostly of olivine, plagioclase, and pyroxene, formed from mantle-derived melts^52^.

Airborne microbes rapidly colonize newly erupted Icelandic lava rocks, and the community is subsequently shaped through a combination of stochastic and deterministic effects^68^. Polymetallic nodules (“PN”) were recovered from the Clarion-Clipperton Zone (CCZ). Nodules can precipitate on and beneath the seafloor at rates as low as a few millimeters per million years, accumulating elevated concentrations of elements such as Ni, Cu, Zn, and Co, among others^69^. Previous microbiological studies have reported a diverse microbial community of Bacteria and Archaea within the nodules, including taxa implicated in manganese and iron cycling^46,50^.

### 4.2. Sample Processing

At each sampling site, rock samples were processed for both DNA sequencing and correlative microscopy analyses. Immediately after collection, one subsample was taken using a sterile hammer and chisel, placed in a sterile Whirlpak bag, and deposited in a -80 ℃ freezer for downstream DNA extraction. A second subsample was chemically fixed in paraformaldehyde (PFA) to preserve cell structures and biological macromolecules. To initiate the fixation, each sample was submerged in 4% PFA in Phosphate Buffer Saline (PBS) overnight (∼16 h) at 4 ℃. Following the fixation, samples were rinsed twice with PBS to remove fixative and stored in 50:50 PBS: Ethanol (EtOH) at -20 ℃ before further processing.

Prior to resin embedding, which was necessary to maintain the precise spatial relationships of microbes and minerals throughout downstream processing and analysis, samples were dehydrated by soaking in increasing concentrations (50, 70, 80, 90, 100%) of anhydrous EtOH in distilled water for 30 minutes each. LR White resin (LRW; Electron Microscopy Science) was selected because of its minimal fluorescence background, low viscosity, and chemically inert nature, as well as its prior characterization via Raman spectroscopy^25^. Infiltration of LRW was performed by soaking each sample in increasing concentrations of resin in EtOH, i.e. LRW: EtOH (50:50), LRW: EtOH (70:30), LRW: EtOH (90:10), and two iterations of LRW 100%. Each step of infiltration was performed in the dark with no agitation for 4 hours, with the exception of the second 100% step, which was performed overnight. After the last soak in pure LRW resin, samples were allowed to cure in the dark at 65°C for 18 hours. Fully hardened, resin embedded samples were sectioned with a macrotome saw (Tech Cut 4, Allied High-Tech Products) using a diamond-tipped blade (4″ outer diameter, 0.5″ arbor, 0.012″ over-diamond thickness) precleaned and lubricated with deionized water. This sectioning exposed a fresh interior surface of each rock fragment for downstream analysis.

### 4.3. Fluorescence and Raman Microscopy Workflow

Sectioned rock fragments were analyzed by confocal fluorescence and then Raman microscopy. Sample surfaces were first stained with SYBR Green (Thermo Fisher Scientific, Waltham, MA), which binds to double-stranded DNA, to visualize putative microbial cells. These stained rock surfaces were imaged using a Leica Stellaris 5 confocal laser-scanning microscope (Leica, Wetzlar, Germany) with a 10x objective lens. We used excitation wavelengths of 405 nm (to assess autofluorescence of minerals such as carbonates), 498 nm (to excite SYBR Green), and 705 nm (to assess autofluorescence of other minerals, such as silicates and oxides), with corresponding emission windows of 420 - 480 nm, 520 - 580 nm, and 720 - 780 nm, respectively. Maps of fluorescence signals were generated by stitching 327 (SC), 264 (VB), 450 (PN) Fields of Views (FOV) together, with an overlap of 10%, using LasX software from Leica. The resolution for the 3 maps was ∼ 0.8 - 1.5 um per pixel.

Four FOVs from each sample were imaged with Raman microspectroscopy, encompassing a total of 0.142, 0.0346, and 0.2363 mm^2^ for the SC, PN, and VB rocks, respectively (Table S3; Fig. S6-S8). Two Raman systems were used in this study: a hyperspectral darkfield XploRA Raman microscope and a LabRAM Soleil Raman microscope (both manufactured by Horiba, Japan). These microscopes use a quartz halogen aluminum reflector lamp (150 watts) and a high-performance LED light source, respectively, for brightfield illumination. Brightfield sample overviews were acquired with a 5x objective using the Mosaic option from the LabSpec Horiba software. This process enabled us to overlay maps acquired with fluorescence microscopy imaging and then identify specific areas that could be further investigated via Raman microspectroscopy. Raman spectra were acquired with a 20x objective for PN and VB samples with the hyperspectral system equipped with a 405 nm laser, and with a 40x objective for SC with the Soleil system equipped with a 325 nm laser. Maps were acquired across 105–2425 cm^−1^ (325 nm laser) and 170-3250 cm^−1^ (405 nm laser) wavenumbers, with 5-7 s acquisition time and a single accumulation. Pixel sizes were 1-7 µm depending on the Field of View (FoV) (Table S3). Fluorescence correction was performed by using the Horiba FLAT Correction tool, an automated fluorescence removal algorithm, applied during acquisition with the 325 nm laser. After data acquisition, a computational pipeline was developed for this study.

### 4.4. Generation of Raman Reference Library

Raman spectra of minerals were downloaded from the RRUFF database^70^, which comprises standard or published spectra of more than 3,700 distinct mineral species. To ensure the retention of high-quality spectra from the RRUFF library, we filtered only for spectra whose identity had been confirmed by XRD and chemical analysis. We interpolated the spectra across the spectral range (100 to 3000 cm^-1^) using a step size of 3 cm^-1^; this was required because the wavenumbers at which data are collected varied across all samples and reference data due to different spectrometer platforms, acquisition parameters, and analysis settings. Some of the mineral spectra covered a shorter spectral range; in these cases, wavelength ranges not acquired in the original spectra were assigned intensity values of 0.001, to avoid introducing zeros that would complicate downstream analysis steps. After our filtration step, the RRUFF repository had no manganese oxide spectra, so we included publicly available Raman spectra of different isoforms of manganese oxides from the published literature^71^.

To add reference spectra of microbial samples, internal and external references were used. We measured reference spectra of the bacterium *Shewanella oneidensis* using freshly grown and chemically fixed pure cultures. To expand the phylogenetic range of microbial Raman spectra, we downloaded publicly available spectra from Kanno et al., 2021^72^. This dataset included spectra of three bacterial (including Gram positive and negative species), and three archaeal pure cultures: *Escherichia coli, Bacillus subtilis, Thermus thermophilus, Thermococcus kodakarensis, Sulfolobus acidocaldarius,* and *Nitrososphaera viennensis*. Finally, we acquired spectra of LRW resin in both liquid (uncured) and solid (cured at 65 °C overnight) forms to add to the database. All of the raw spectra from internal and external references were processed via baseline correction (lambda=4, p=0.0001, iteration = 500, method=*’als’*) and Savitzky– Golay filtering and smoothing (p=2, n=15) in Rstudio by using package *baseline* and *lsa*, respectively.

We used these same parameters when processing our raw experimental data prior to interpolation and spectra decomposition analysis.

### 4.5. Spectral Decomposition

Raman spectra of natural materials are often highly complex due to the large number of vibrational modes and the frequent occurrence of overlapping bands. Unequivocal assignment of all spectral features is often challenging, and identifications are frequently performed using diagnostic Raman bands, where a single peak can serve as a reliable spectral marker for a specific molecular component. For this reason, we performed spectral decomposition, enabling the identification and quantification of individual Raman bands that may otherwise be obscured within the composite spectrum.

Non-Negative Matrix Factorization (NMF) was applied to decompose processed Raman spectra from each pixel into their constituent spectral components while preserving the non-negativity of all values. Following interpolation, each spectrum was normalized between 0 and 1 before applying NMF, thereby reducing the influence of absolute intensity differences and enhancing sensitivity to spectral shape variations while facilitating comparison across samples.

NMF is a dimensionality-reduction technique that decomposes a nonnegative data matrix X into the product of two lower-rank nonnegative matrices W and H, such that X is roughly the product of W * H. In our case, matrix X represented each sample’s Raman map, composed of pixel spectra and their position on the map. Each row in the matrix corresponded to a different pixel spectrum, while the columns represented the Raman shift values at each wavenumber. From this we obtained W, a matrix with Raman shift as rows and components as columns (such that each cell contained the intensity value at that specific wavenumber for the specific component) and H, a matrix with components as rows and pixels as columns (where each cell contained the weight of that component per each pixel). Because all components are constrained to be nonnegative, the resulting factors provide a more interpretable representation of the original data than other decomposition approaches^73^. This process allowed us to identify the types of minerals and organics in each pixel as well as their relative contributions to the full spectrum.

We performed NMF in R studio (version 2.6.1) using the package “*nmf*” (version 0.24). To achieve an optimal balance between silhouette and variance metrics, parameters were set to n.run = 5 and rank = 15. We combined four FoVs from each sample, interpolated the data as described above, and ran the *nmf* to individually decompose the spectra of each sample type and identify the main spectral components (Data Set 1). For each sample, the output consisted of maps showing each identified component and its relative contribution to the original spectrum (the weights) for each pixel (Extended Fig. 3).

To agnostically identify the components of our experimental Raman spectra, the pairwise cosine similarity between each component’s spectrum and each mineral and microbial spectrum in the reference library was calculated using *lsa* package in R, according to equation 1, where A and B are vectorized spectra being compared.

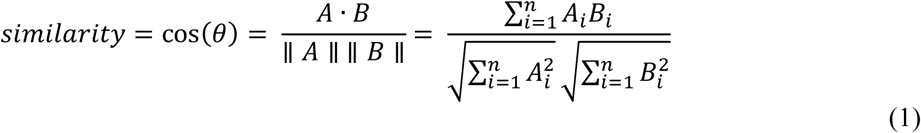

Cosine similarity was used to assess spectral correlation, as it emphasizes similarities in spectral shape while minimizing the influence of absolute intensity differences between spectra. Cosine similarity compares two spectra by treating their intensities as vectors and measuring the cosine of the angle between them, so spectra with similar peak patterns will have values close to 1 and spectra orthogonal to each other will have values close to 0. This approach is widely used for spectral data and library matching^74^. A threshold of 0.7 was chosen to select only strong matches between components and reference spectra^75^. Although no universally accepted cutoff exists for Raman or other spectroscopic techniques, threshold selection represents a trade-off between breadth and specificity, and usually is dataset-dependent given variations in calibration, spectral quality and variability, preprocessing, and signal-to-noise ratios^76,77^. Previous work on surface-enhanced Raman spectroscopy has shown that correlations between unrelated spectra rarely exceed 0.6, and that strong similarities are reliably detected above this threshold^78^. For our analysis, each pixel’s identity was assigned as the component with the highest weight; often, multiple top matches corresponded to different database entries of the same substance (mineral or biomass type) (Data Set 1), offering additional confidence of the identification. We manually compared the top match(es) with each component spectrum to confirm the alignment of spectral features (Fig. S1-S3).

### 4.6. Quantifying Compositional Heterogeneity Within and Across Pixels

The selection of suitable heterogeneity metrics depends upon the specific research question being addressed and the data structure^79^. Four different quantitative measures were used to test our hypothesis that the presence of biomass correlates positively with regions of greater compositional heterogeneity. Two of these measures (Gini index and the Shannon entropy index) assessed the heterogeneity of components within pixels (intra-pixel), and two of them (Spectral Correlation and Rao’s quadratic entropy, or Rao’s Q) assessed heterogeneity between a given pixel and its neighbors (inter-pixel). These metrics are appropriate for compositional data, as they capture complementary properties of composition, evenness, and pairwise dissimilarity within pixels.

For intra-pixel heterogeneity, two metrics were selected because they capture distinct aspects of heterogeneity: the Gini index^80^ emphasizes inequality and dominance of one component within each pixel, while Shannon entropy^81^ measures diversity in components’ distributions, taking both richness and evenness into account. Shannon metrics were calculated after normalization of each pixel and thresholding. In compositional data, low-abundance components are often associated with high relative uncertainty^82^; therefore, a 10% abundance threshold was applied to remove low-abundance components for Shannon calculations, reducing the influence of noise and enhancing meaningful differences between pixels. For interpixel heterogeneity, our metrics focused on two different types of dissimilarity: while spectral correlation emphasizes spectral similarity by examining all adjacent pixels and correlating their entire Raman spectra in an agnostic manner, the modified Rao’s Q measures compositional diversity, weighted by pairwise distance between pixels and across components. In each case, values were determined for every pixel of areas scanned with Raman microspectroscopy, generating heat maps that could be directly compared with other pixel-scale data such as the spectral component weights.

#### 4.6.1. Intra-pixel Heterogeneity Metrics

The *Gini index* was originally applied to understand income distribution patterns within a population, but more recently, it has gained traction in biological applications, as in transcriptomics analyses measuring inequality of gene expression under different contexts^83^, chemical biological studies assessing the binding affinity with specific molecules^84^, and single-cell studies quantifying phenotypic heterogeneity in metabolic activity within a microbial community^85^. We used the Gini index to understand how equally-distributed the components that comprise each pixel are: if the value is high, one or a few components dominate, indicating an unequal distribution. Conversely, a low Gini index indicates a more heterogeneous composition, in which components are more evenly distributed within each pixel. The Lorenz curve (Fig. S9) was calculated for each pixel, plotting the cumulative percentage of component weights against the cumulative percentage of components. Gini values were computed according to

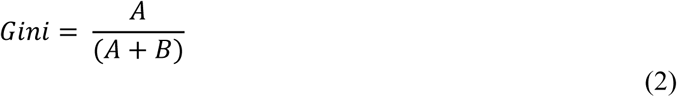

where A is the area between the line of perfect equality and the Lorenz curve, and B is the area between the Lorenz curve and the perpendicular lines bounding it. Calculations were performed using the *gini*() function from the *ineq* package in R. For directional consistency across all heterogeneity metrics, we report “1 - Gini” throughout, such that higher values correspond with higher heterogeneity. Hereafter, we refer to this parameter as the Gini index.

*The Shannon entropy index* (hereafter, “Shannon”) is a quantitative measure of diversity that incorporates a distribution’s richness and evenness according to equation 3

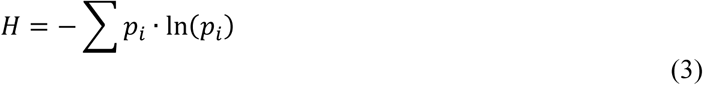

where H is the diversity index and p_i_ is the weight of each spectral component (i.e., the proportion of the sample spectrum accounted for by the component). Compared with other alpha diversity metrics such as the (Inverted) Simpson or Pielou’s indices, Shannon is sensitive to changes across the entire abundance distribution, including rare, low-abundance components. (Its formula uses the natural logarithm of each component’s weight, so even a small value contributes significantly to the overall sum). Shannon was computed in Rstudio by using the *vegan* package, choosing ad diversity index = “shannon”.

#### 4.6.2. Inter-pixel Heterogeneity Metrics

*Spectral correlation* was used as a compositionally agnostic measure of “mineralogical complexity”, quantifying the difference between a given pixel and its neighboring pixels. All pairwise combinations of a central pixel and each of its eight neighboring pixels were established; for each pair, the cosine similarity was calculated between the two baseline-corrected and interpolated spectra. By averaging the eight cosine similarity values, a “spectral heterogeneity score” (SHS) for the central pixel was determined as “1 - mean cosine similarity”, such that higher values corresponded with higher inter-pixel heterogeneity. To avoid edge effects and smearing, for the pixels at the edge or at the corners of each FoV, cosine similarity average for the central pixel was calculated just with the 5 and 3 neighboring pixels, respectively.

*Rao’s quadratic entropy* (Rao’s Q) was introduced as an ecological metric that quantifies landscape diversity by computing the average spectral dissimilarity between pairs of pixels within a moving window^86^. In this context, dissimilarity can be measured as distance in any metric (e.g., phylogenetic, taxonomic, or morphological distance); we used the proportion (weight) of each component and distances between components per each pixel, according to equation 4.

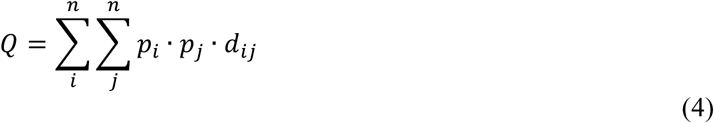

In this equation, n is the number of components, p_i_ and p_j_ are the weights of each component across all pixels, and d_ij_ is the distance between components. Rao’s Q will be low if pixels in the moving window are spectrally similar to each other (i.e., a small d_ij_) and high when they are dissimilar (a large d_ij_). To calculate Rao’s Q, we followed the framework described by Rocchini et al.^63^ and modified the *spectralrao* original function^87^ in R. The two major customizations were computing the mean pairwise Euclidean distance between neighboring pixels and reducing the window size at pixel borders instead of using a fixed-size neighborhood. For each central pixel in the moving window, eight neighbors are expected; however, edge and corner pixels have five and three neighbors, respectively. To correct for the edge effect, we considered only the real neighbors and averaged across them as described above for the SHS.

### 4.7. DNA Extraction & Sequencing

When coupled with 16S rRNA gene high-throughput sequencing, our microspectroscopy workflow can be used to derive site-specific findings of ecophysiological links between biomass abundance, taxonomic identities, mineralogical heterogeneity, and particular mineral components. Interior portions of each rock, defined as volumes at least 5 cm from an exterior surface, were crushed with a sterile mortar and pestle and DNA was extracted with the Qiagen DNeasy PowerSoil Pro genomic extraction kit using the manufacturer’s instructions. Negative controls for each extraction batch, which used double deionized (MilliQ) water rather than rock, were below detection limit when measured by a Qubit 4 fluorometer and were not sequenced. Extracted DNA was sent to Laragen Inc. (Culver City, CA), where it was amplified using the 515F (5′-GTGYCAGCMGCCGCGGTAA)^88^ and 806R (5’- GGACTACNVGGGTWTCTAAT)^89^ primers first developed by Caporaso et al.^90^. 100 ng of PCR amplified DNA was sequenced using a PE 150 kit on an illumina MiSeq. Raw sequencing reads were processed and rarified using an implementation of QIIME2^91^ hosted on Boston University’s Shared Computing Cluster. First, dada2 was used to generate amplicon sequence variants (ASVs) with default settings to filter reads with two or more expected sequencing errors and truncate reads at the first instance of a quality score of 2. On average, 90% of all reads from each sample passed our quality control criteria. Following this filtering step for each read, we required a minimum overlap of 12 bases between forward and reverse reads, resulting in >80% of reads successfully merging after removal of chimeric reads.

Unjoined reads were discarded and not included in our analyses. Following the generation of ASVs with dada2, ASVs were classified with 95% or greater sequence identity to the SILVA_132_99 database^92^ with VSEARCH^93^. ASV counts were then batch corrected using ComBat-seq^94^, a negative binomial regression model, to account for technical variation between sequencing runs.

### 4.8. Statistics & Reproducibility

Differences in compositional heterogeneity between biomass and mineral pixels were assessed within each sample type (SC, VB, and PN), separately for each index (intra-pixel: Gini and Shannon; inter-pixel: SHS and Rao’s Q), and for each condition (all components, dominant component omitted, biomass component omitted). The biomass component combined biomass and kerogen top hits, and resin pixels were excluded. The unit of observation is a single map pixel, and n is the number of biomass and mineral pixels pooled across the four fields of view (FOVs) per each sample type.

Statistical methods were used to test the magnitude of the differences between biomass and mineral pixels, as well as to check their statistical significance. Initially, p-values were calculated for each comparison by means of Wilcoxon test (Data Set 2). Because these maps contain thousands of pixels per group, conventional null-hypothesis significance tests may not fully capture biological importance, as small p-values can incorporate a dependence on the pixel count rather than exclusively depicting the size of the effect^95,96^. To account for this issue, we report Cliff’s delta (δ) for biomass-versus-mineral comparisons. Cliff’s delta is a non-parametric measure of effect size that equals the difference between the probability that a randomly chosen biomass pixel exceeds a randomly chosen mineral pixel and the reverse probability^97^. It is bounded on [−1, 1] (δ = 0 under stochastic equality), is invariant to monotone rescaling, and makes no distributional assumption, which is appropriate for our study because the indices are bounded and skewed. By our sign convention, δ > 0 indicates that biomass pixels tend to exceed mineral pixels in terms of the heterogeneity indices. We additionally report the Hodges–Lehmann median shift (the median of all pairwise biomass − mineral differences) as an effect size in the original index units^98^ (Supplementary Information, Data Set 2).

For the inter-pixel indices (1 − SHS and Rao’s Q), map pixels are spatially autocorrelated by design, so the pooled pixels within a FOV are not independent. As a sensitivity check against spatial non-independence, we repeated the effect-size calculations after intensively spatially thinning each FOV to approximately one pixel per spatial-decorrelation block (block size set from a per-FOV empirical variogram; 1,000 random thinnings). For each thinned dataset, we repeated the primary analysis and compared the direction of the estimated effect size with those obtained from the full dataset, summarizing the proportion of thinned datasets that retained the original effect (Table S2).

## Supporting information

Data Set 1

Data Set 2

Data Set 3

## Acknowledgments

The authors would like to thank collaborators and research partners involved in sample acquisition. For work at the Point Dume methane seep, we thank the science team and crew of R/V *Falkor* leg FK181005 in October 2018, particularly Dr. Peter Girguis and Dr. Daniel Hoer. Collection of the polymetallic nodule sample occurred as a part of The Metals Company’s environmental baseline study aboard the Maersk *Launcher* in 2021. For work at Fagradalsfjall, we thank Dr. Christopher Hamilton, Erin Frates, and Dylan Mankel.

Support for this work came primarily from NASA Exobiology Grant 80NSSC23K0224 to JM, AR, HG, and DEL. DEL also acknowledges financial support from the NASA Habitable Worlds program under grant 80NSSC20K0228, the NASA Exobiology program under grant NNH22ZDA001N, and the Claude C. Albritton, Jr. Endowment at Southern Methodist University. SD was supported by NSF RAPID grant #2128606, and NH was supported by Geological Society of America Graduate Student Grant 13183-21 and the University of Arizona Graduate and Professional Student Council Travel Grant.

## 5. Code availability

All preprocessing, analysis, and visualization scripts, which are detailed above, are publicly available through a GitHub repository (https://github.com/Fcalabrese-11/mineralogical_heterogeneity).

---

**Extended Figure 1.**
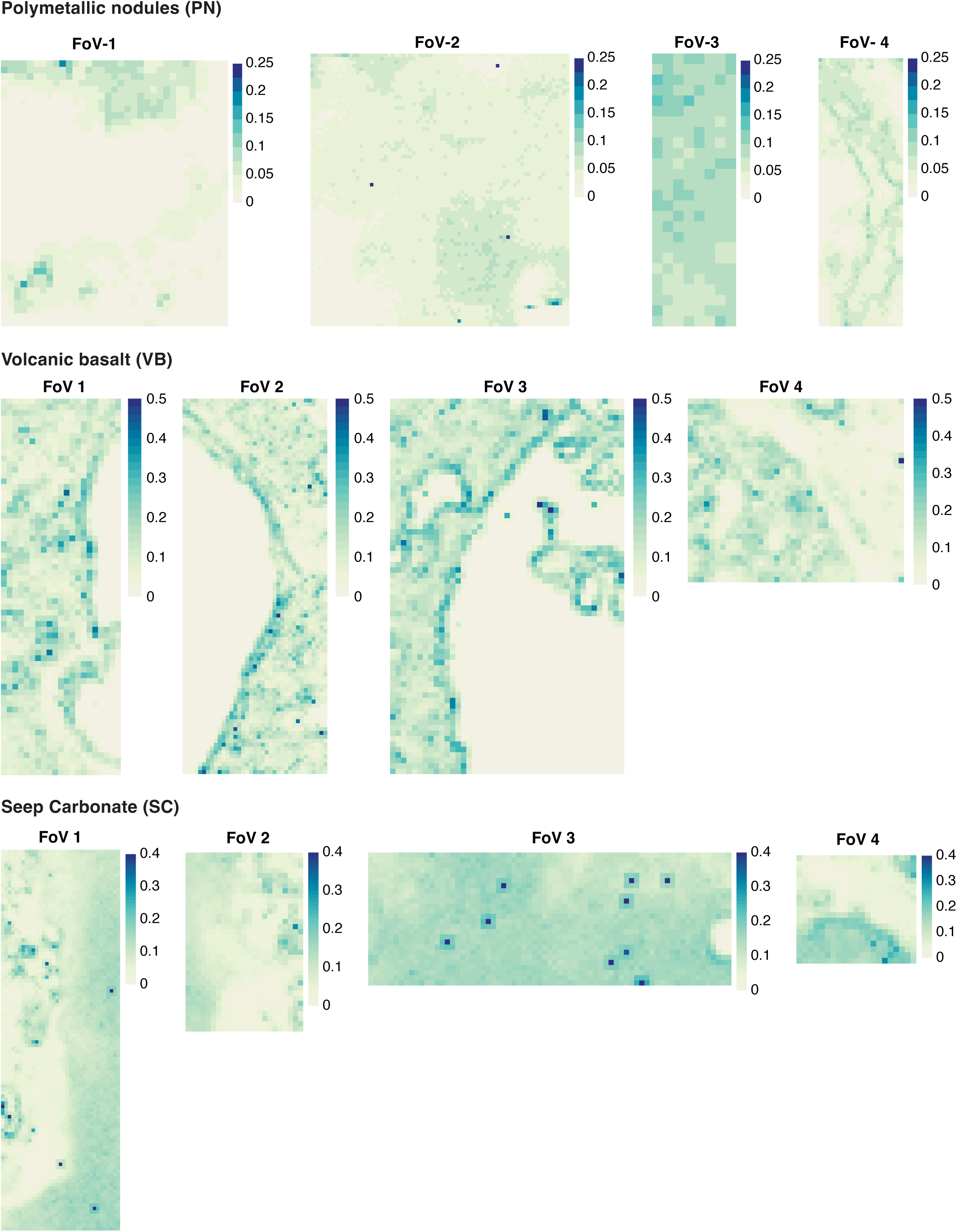
Spectral Heterogeneity Scores (SHS) shown as heatmaps for each field of view (FoV) for each sample. The color scale indicates SHS values, ranging from low (white) to high (blue).

**Extended Figure 2.**
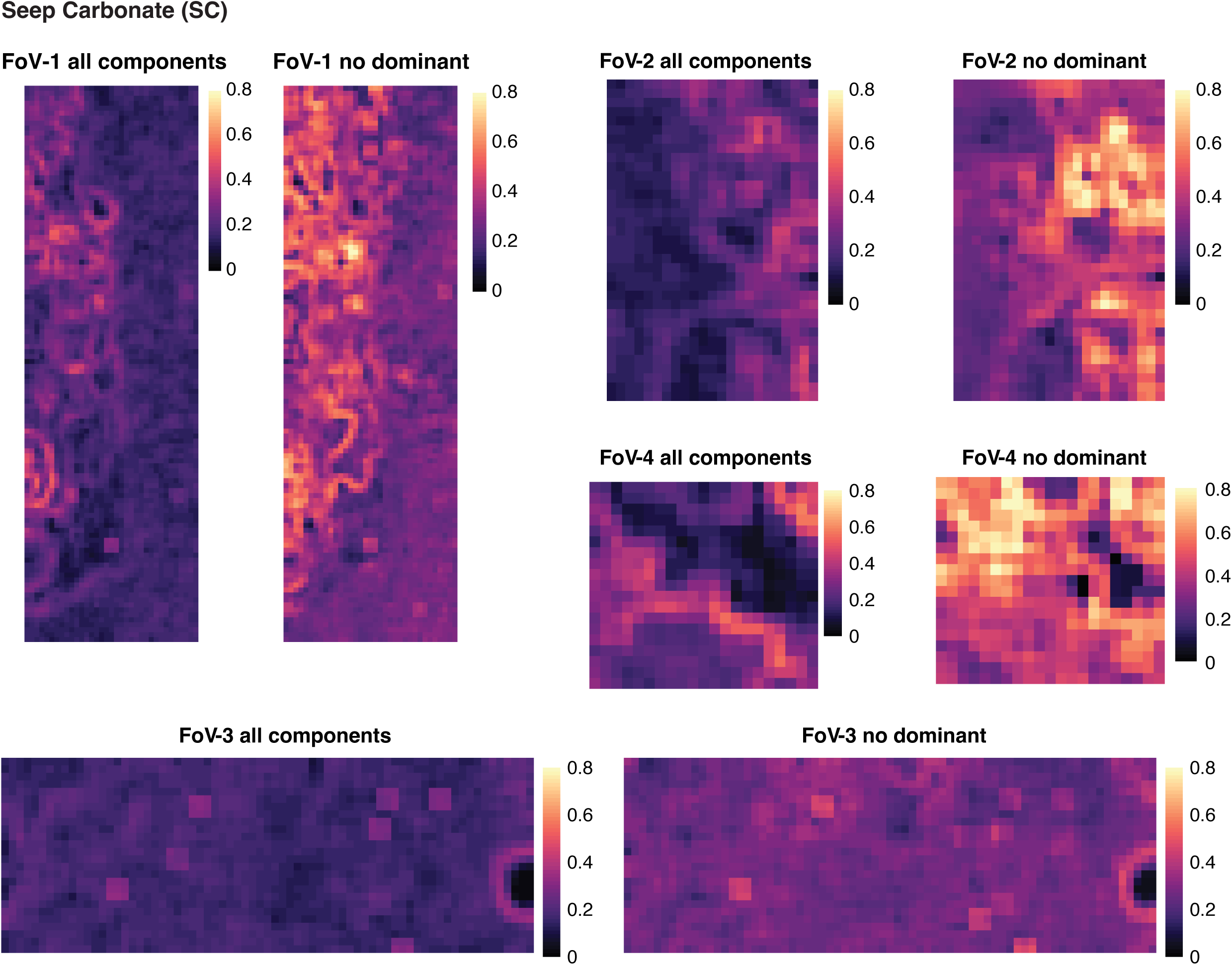
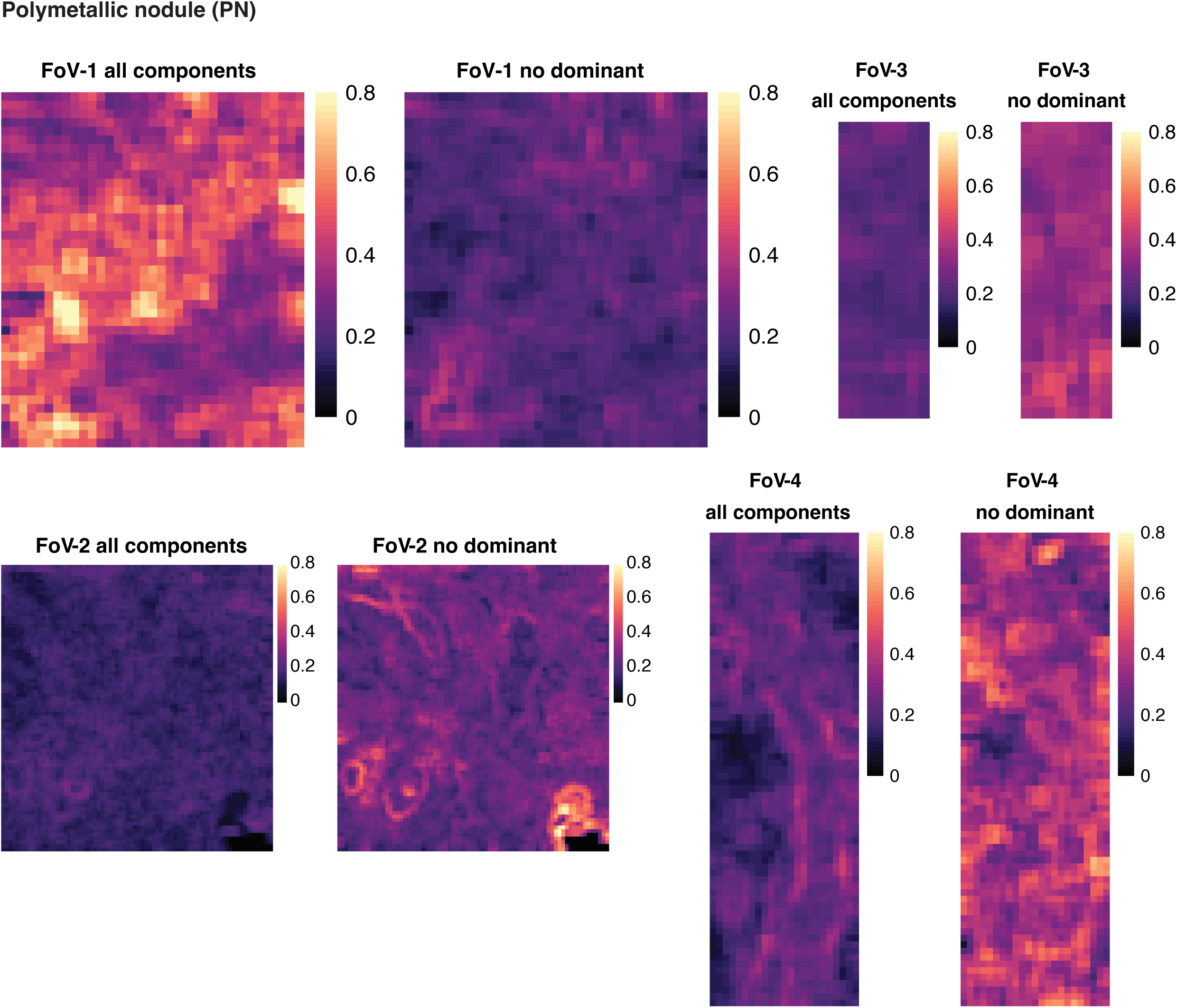
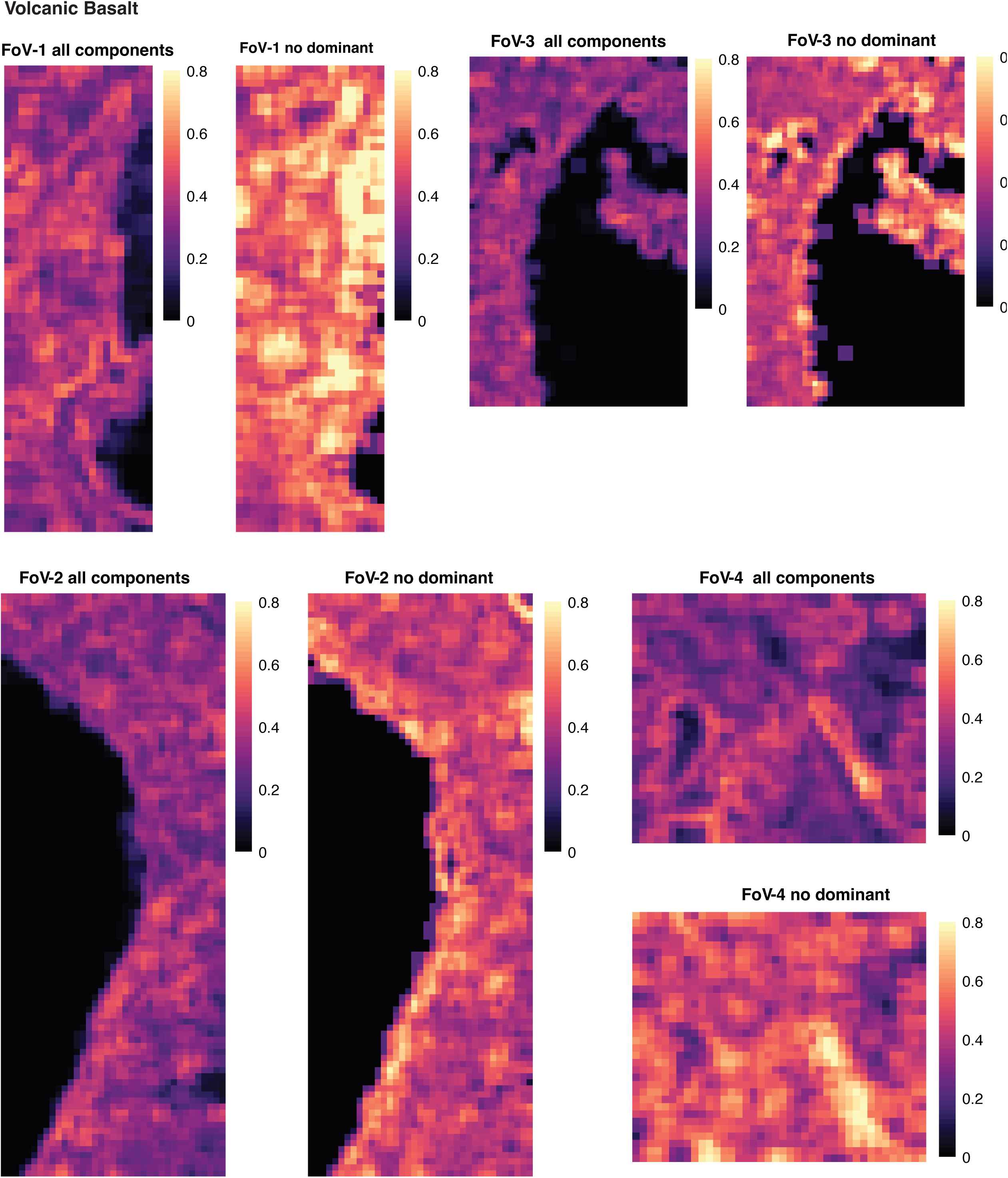
Rao’s quadratic entropy (Rao’s Q) is a mathematical measure of biodiversity which calculates the average difference or dissimilarity between two randomly chosen individuals in a community. We applied this index to pixels to measures the compositional heterogeneity, i.e. how distinct pixel composition is across space and across different type of pixels. For each pixel, we calculate an average score by combining relative component abundance and weight-value differences within a moving window of adjacent pixels. We then compare the difference between pixels identified as biomass and pixels identified as minerals. We also compared difference between the same groups when all components were included in the calculation (except for resin) and when the dominant (primary) component was omitted from the calculation. The maps below represent each Field of View (FoV) in each sample, in the presence and absence of a dominant component. The color scale indicates the values of Rao’s Q index.

**Extended Figure 3.**
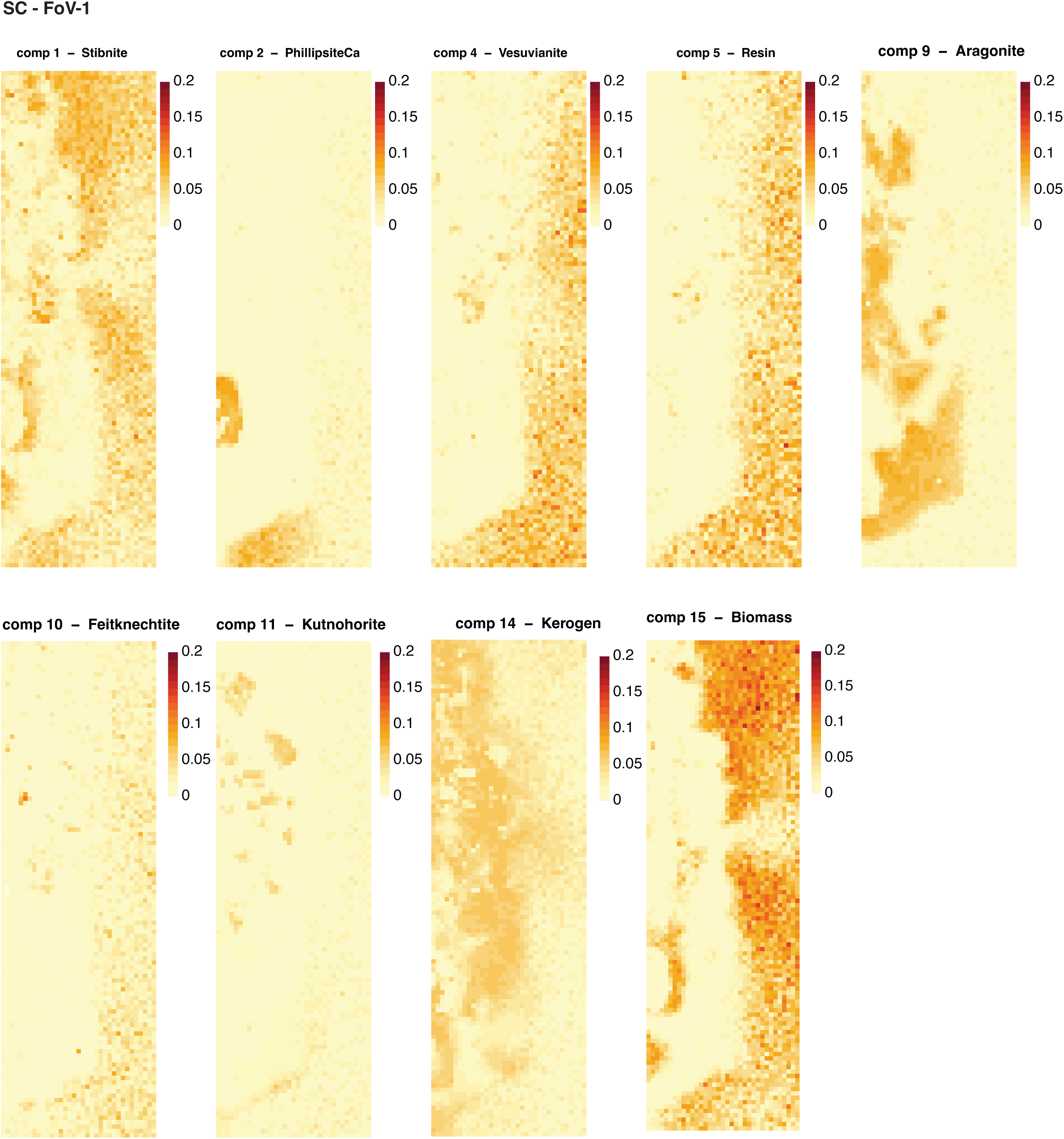
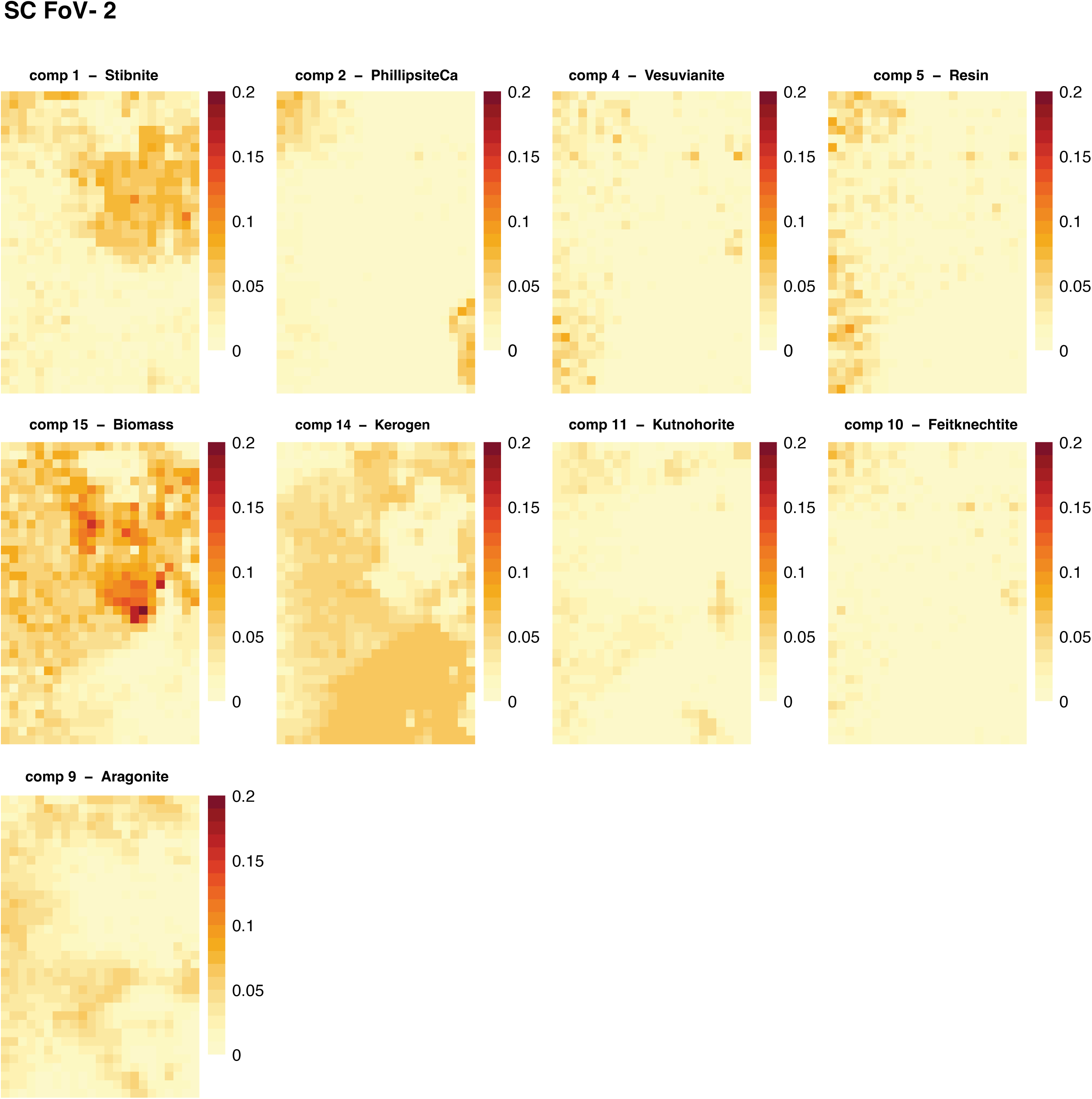
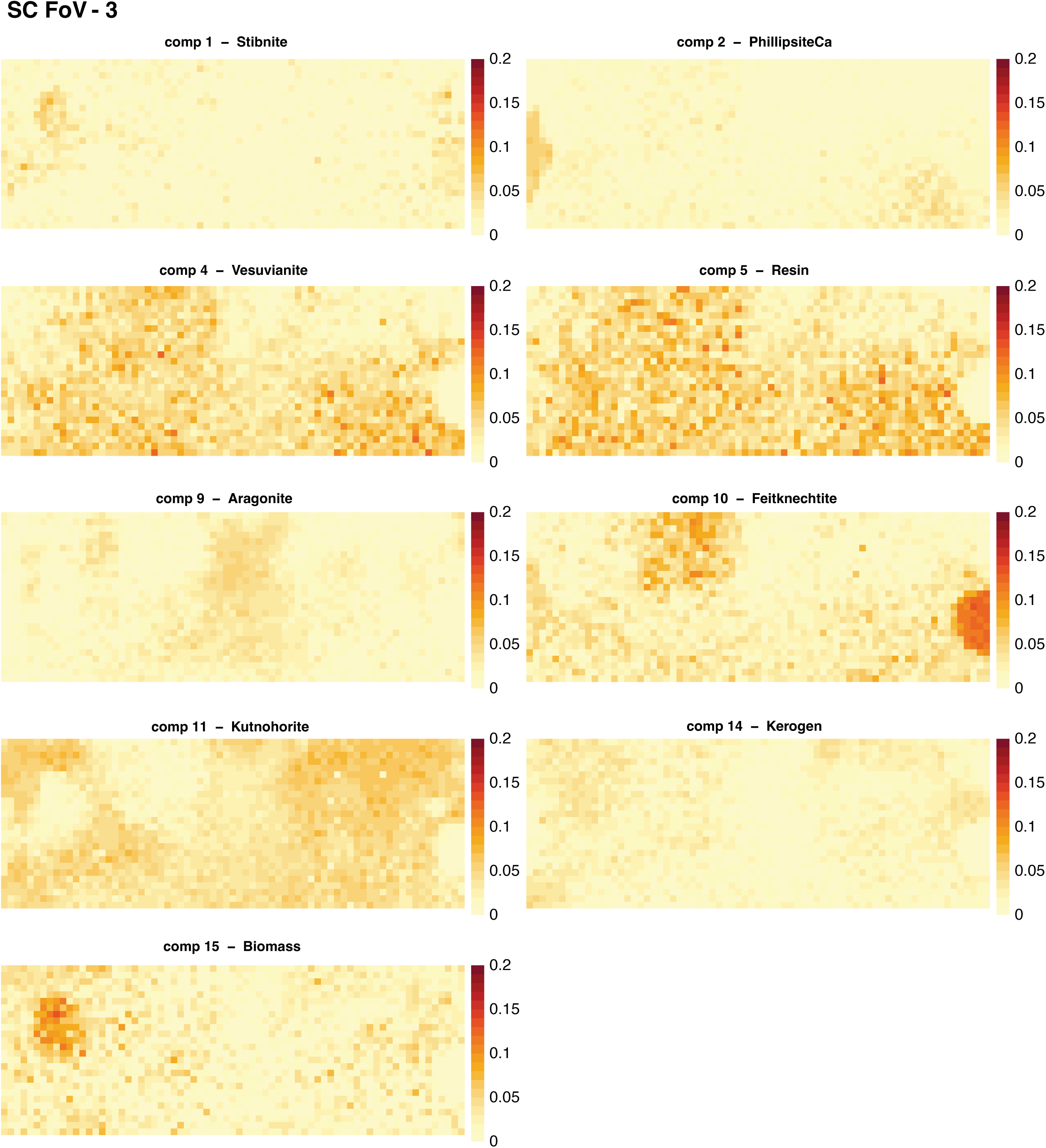
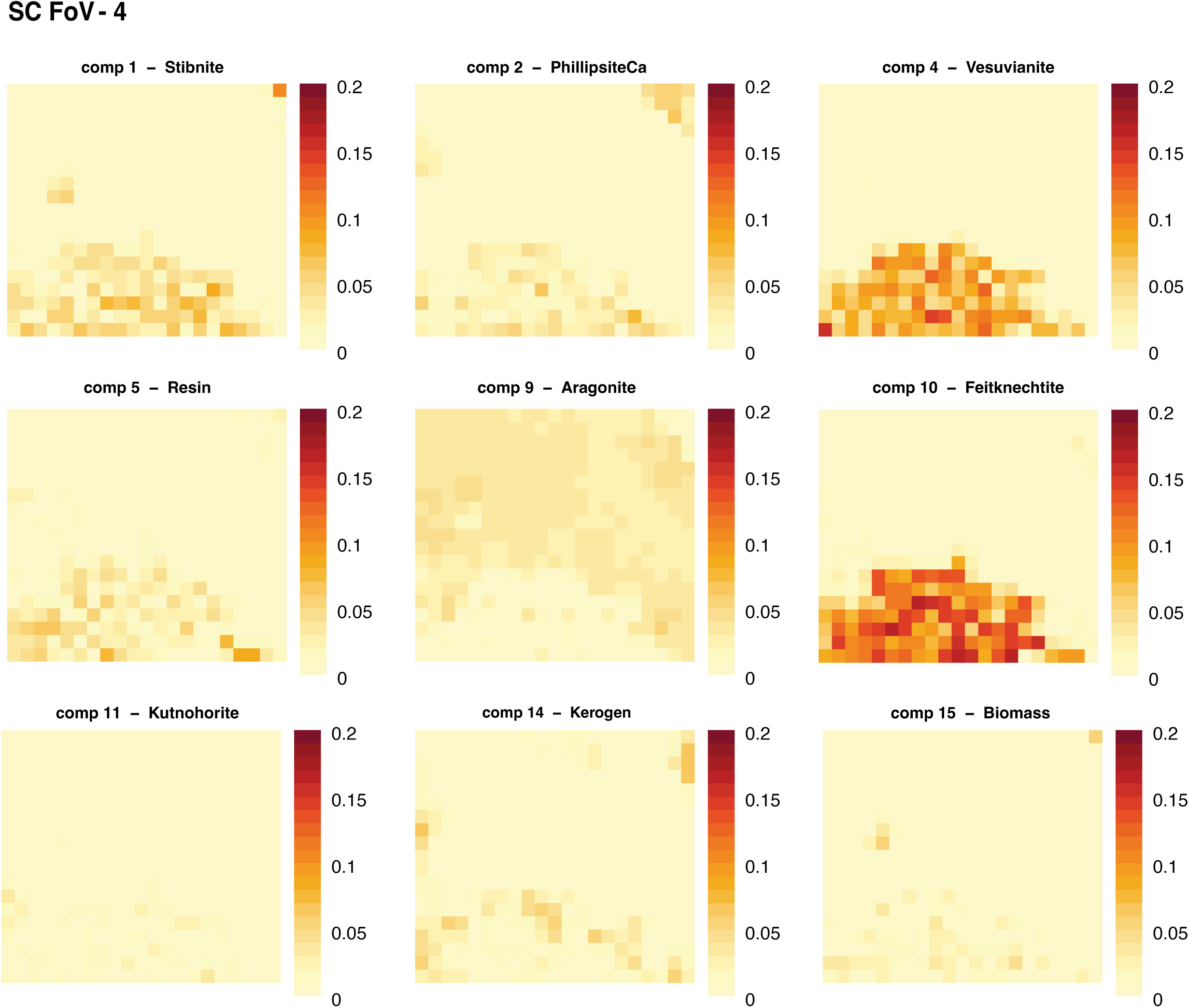
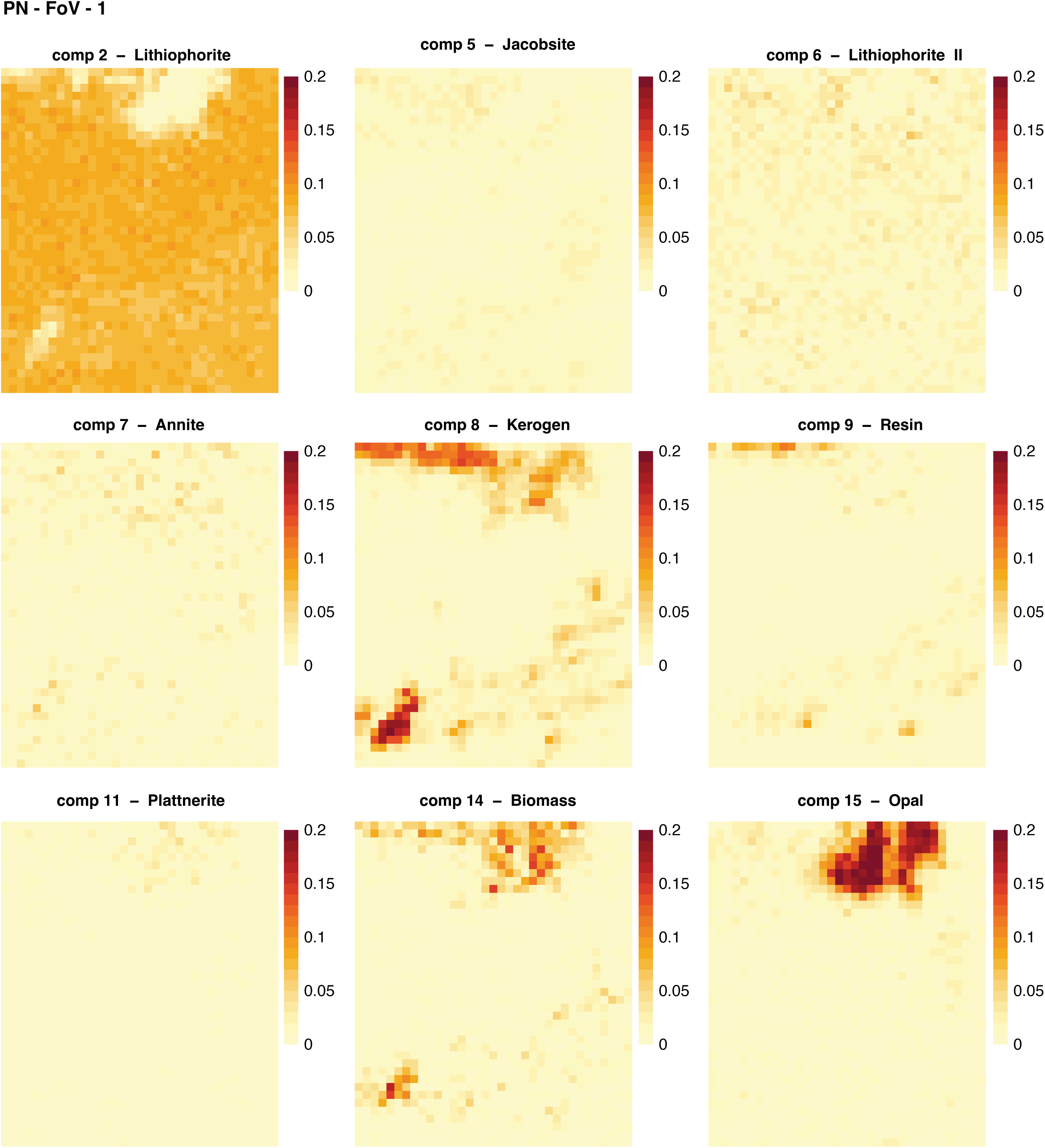
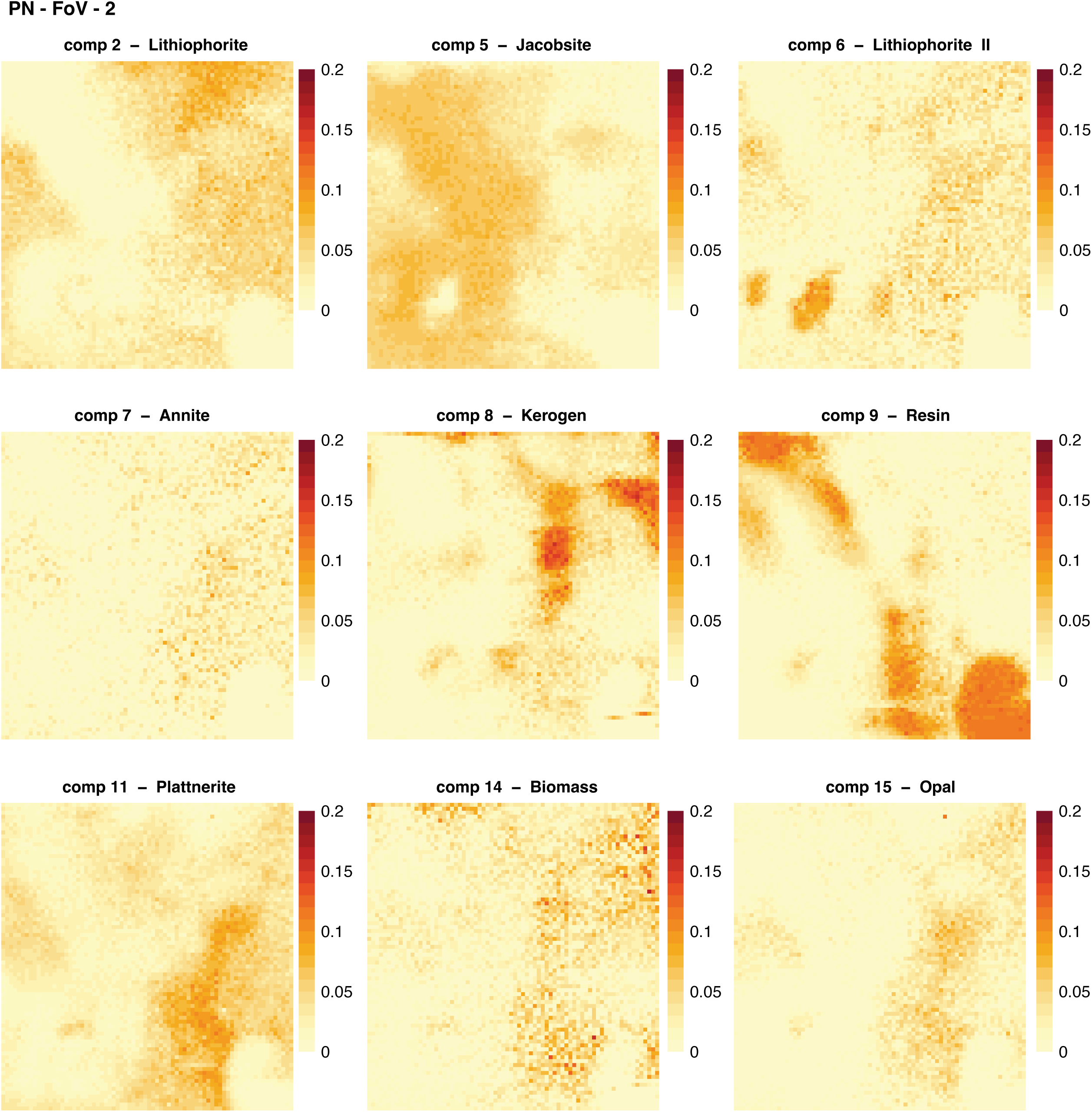
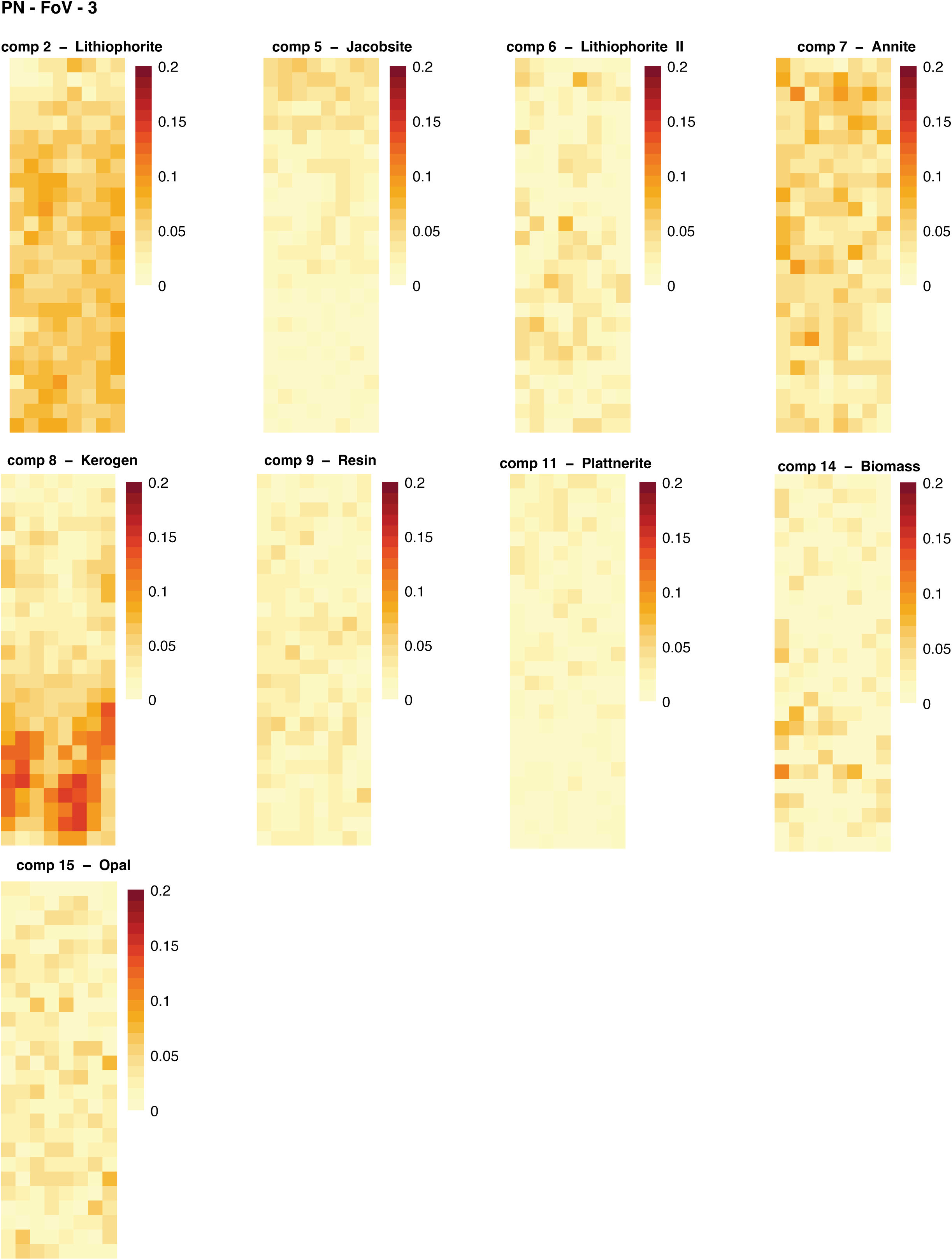
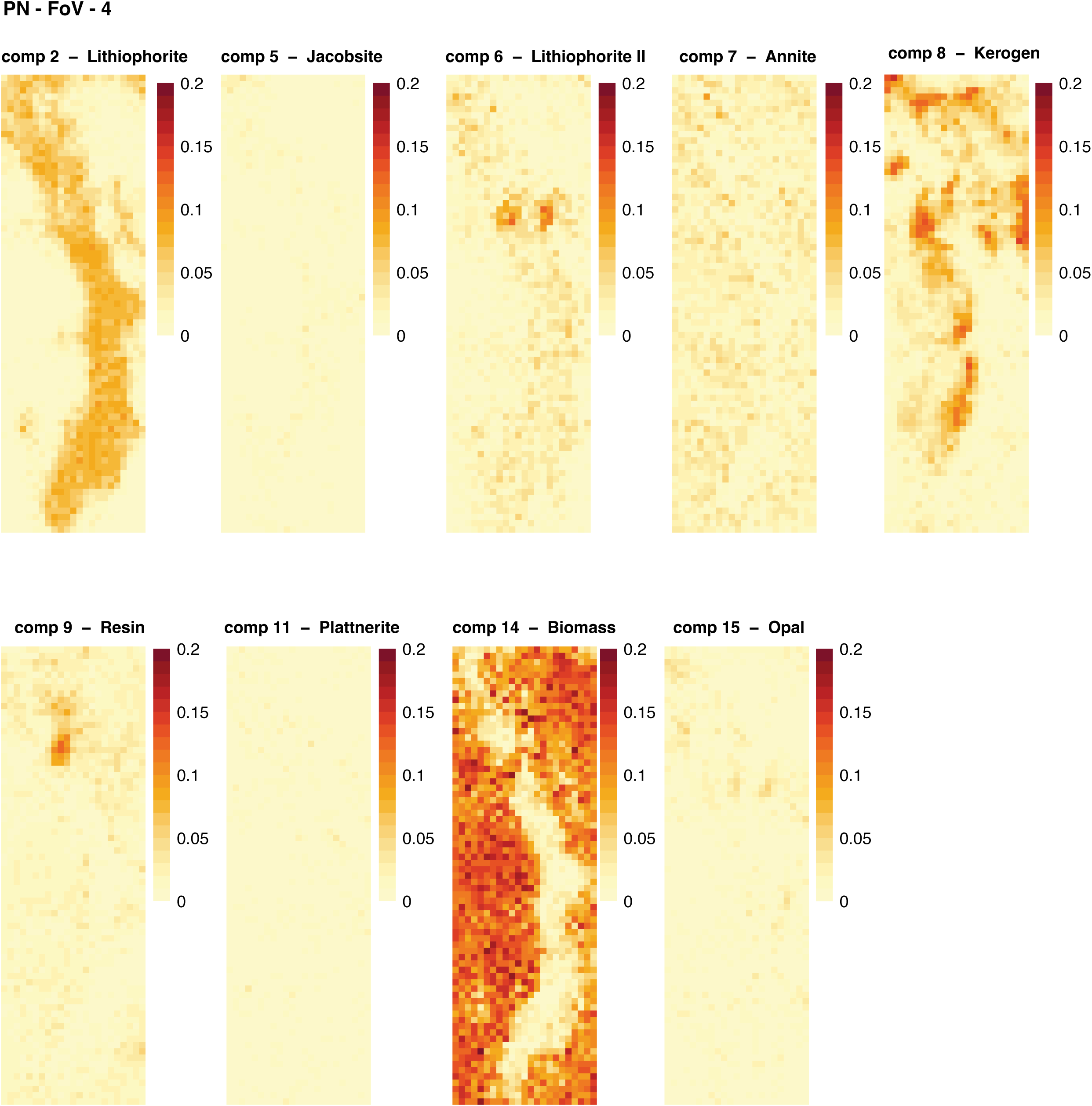
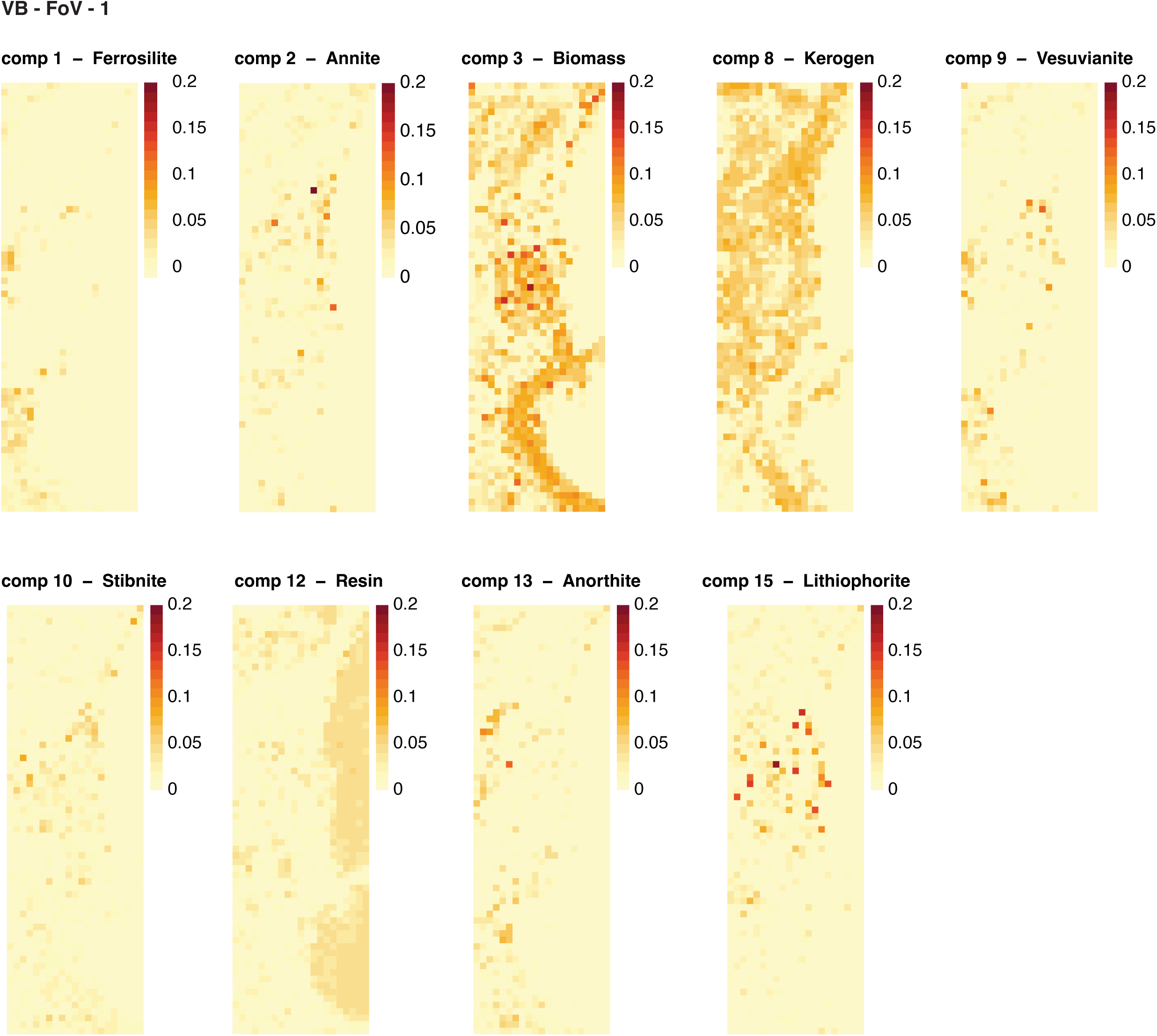
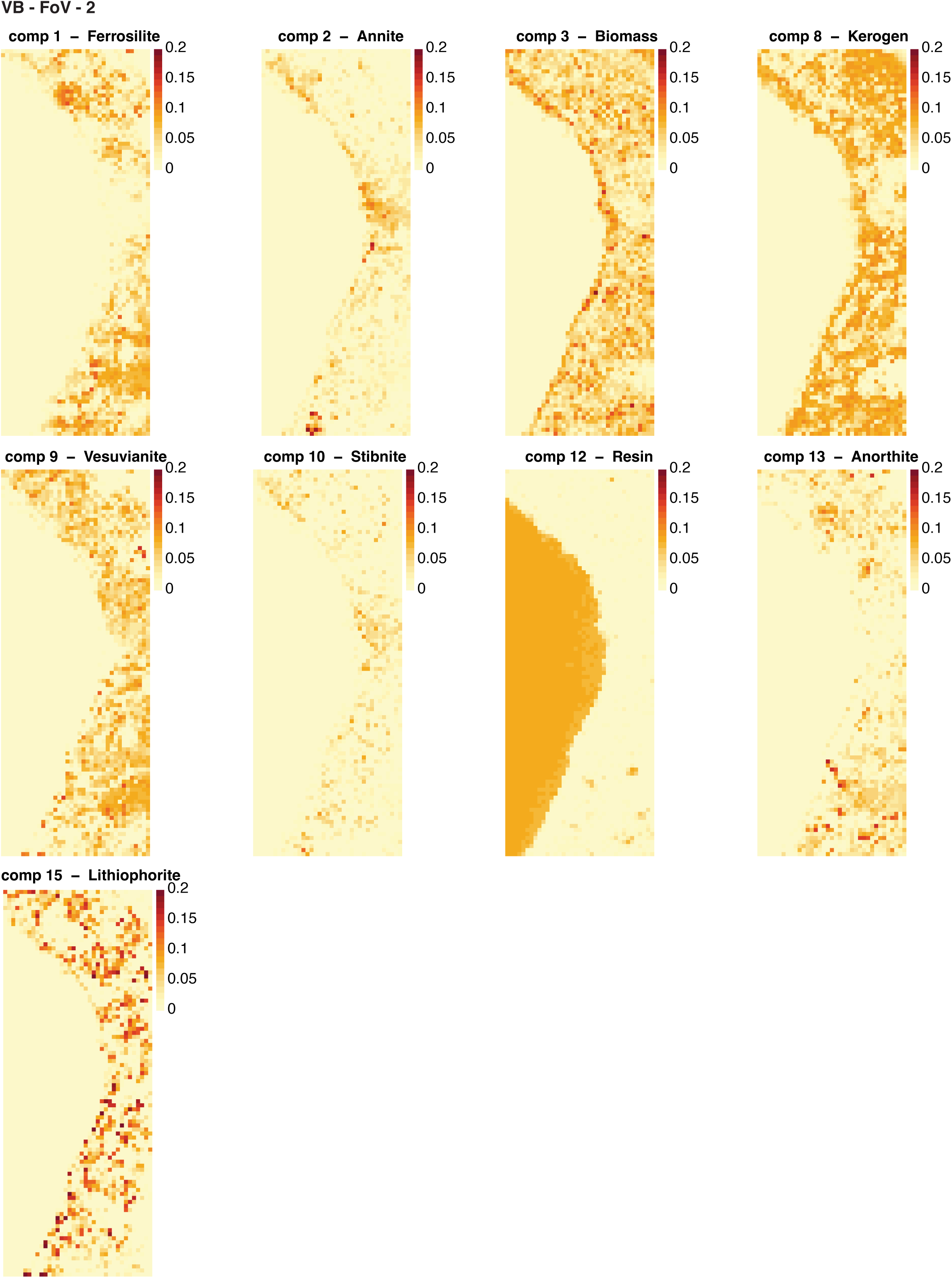
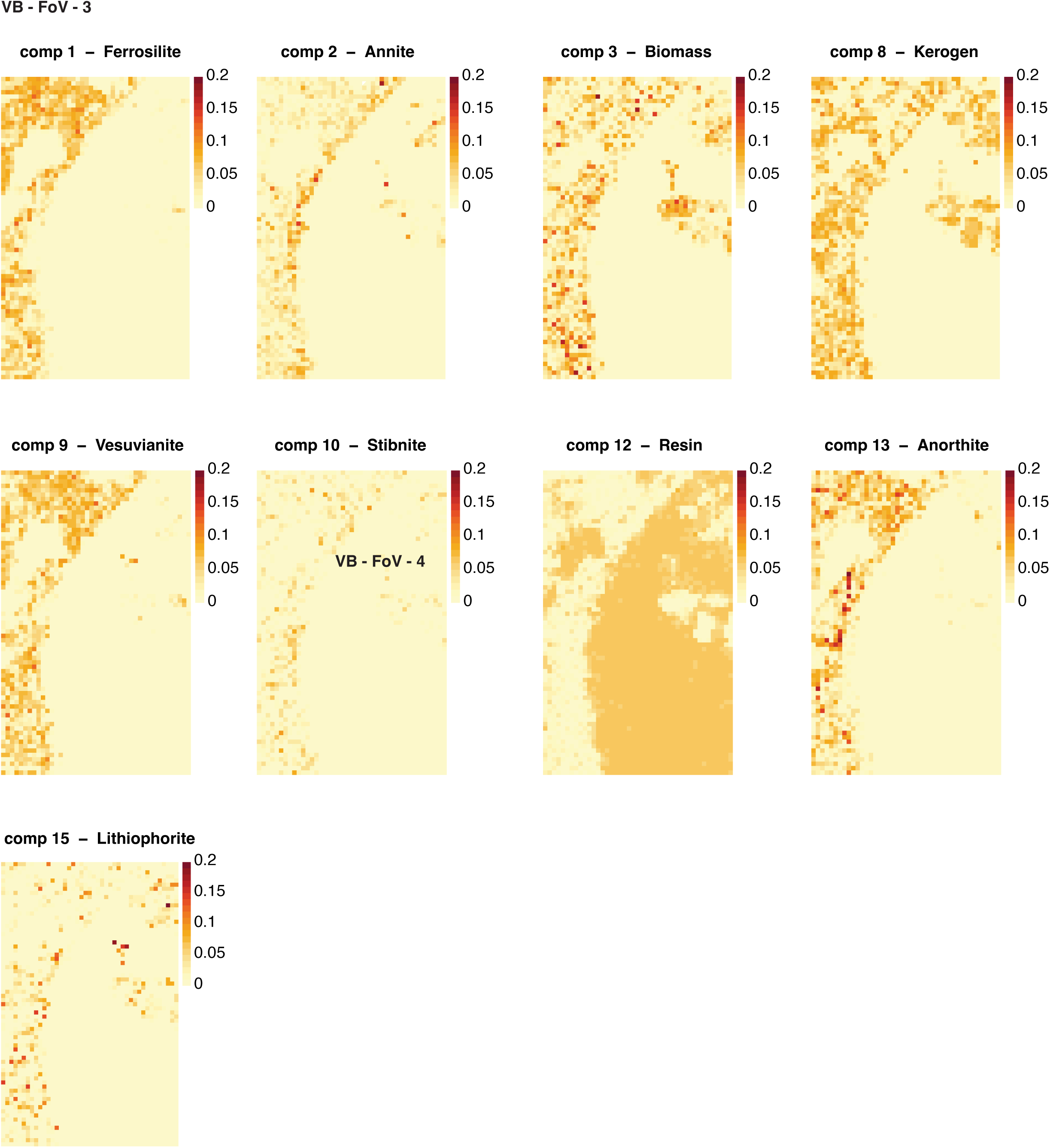
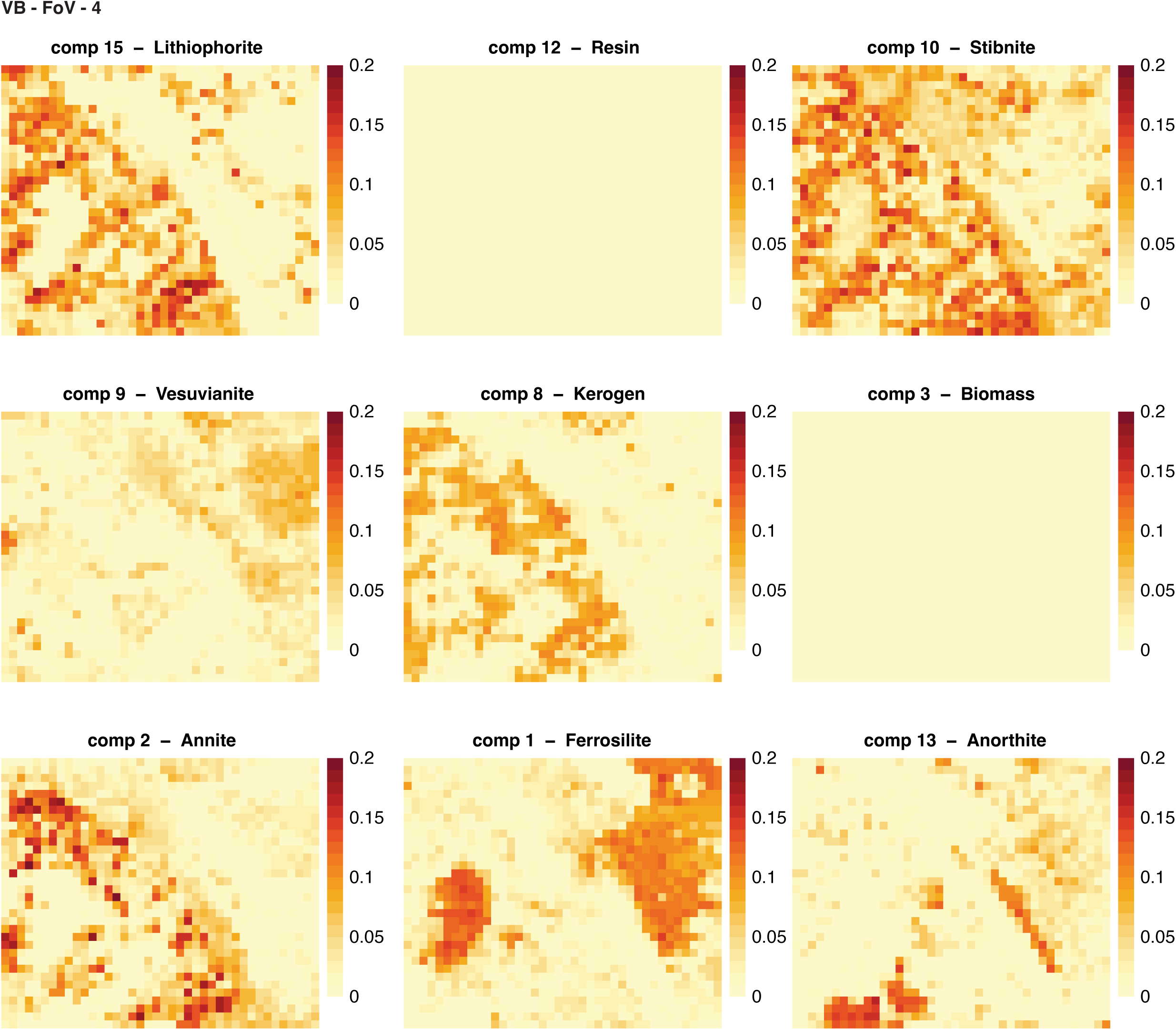
After spectral decomposition with Non-negative Matrix Factorization (NMF), we identified nine components for each type of sample. Below are the maps of each individual component for each Field of View (FoV) in each sample. The color scale represents the weight of each pixel for a specific component. The four FoVs analyzed for each sample in this study are shown below for SC, PN and VB, respectively.

