## Supplementary information for "Compositional Heterogeneity Structures Microbial Microhabitats across Distinct Mineral Substrates"

Note: sections in this document, including their associated figures and tables, appear in the order in which they appear in the Main Text.

### 1. Non-Negative Matrix Factorization (NMF) and identity of the components

Raman spectra decomposition was performed with NMF for the three samples, SC, VB, and PN. To further validate the reliability of this computational matching approach, we performed manual peak matching using component and database spectra. The data are provided in Fig. S1-S3 below. For the majority of the components, the peaks in their spectra (first row in each panel of Fig. S1-S3) align with the fingerprint peaks of the "best match" mineral class ($2^{nd}$ to $4^{th}$ rows), which are typically used in the manual peak matching procedure.

Fig. S1. Manual peak matching for the SC components. The top row for each component is the component's spectrum, while rows 2-4 are the spectra of the top three matches from our expanded reference database.

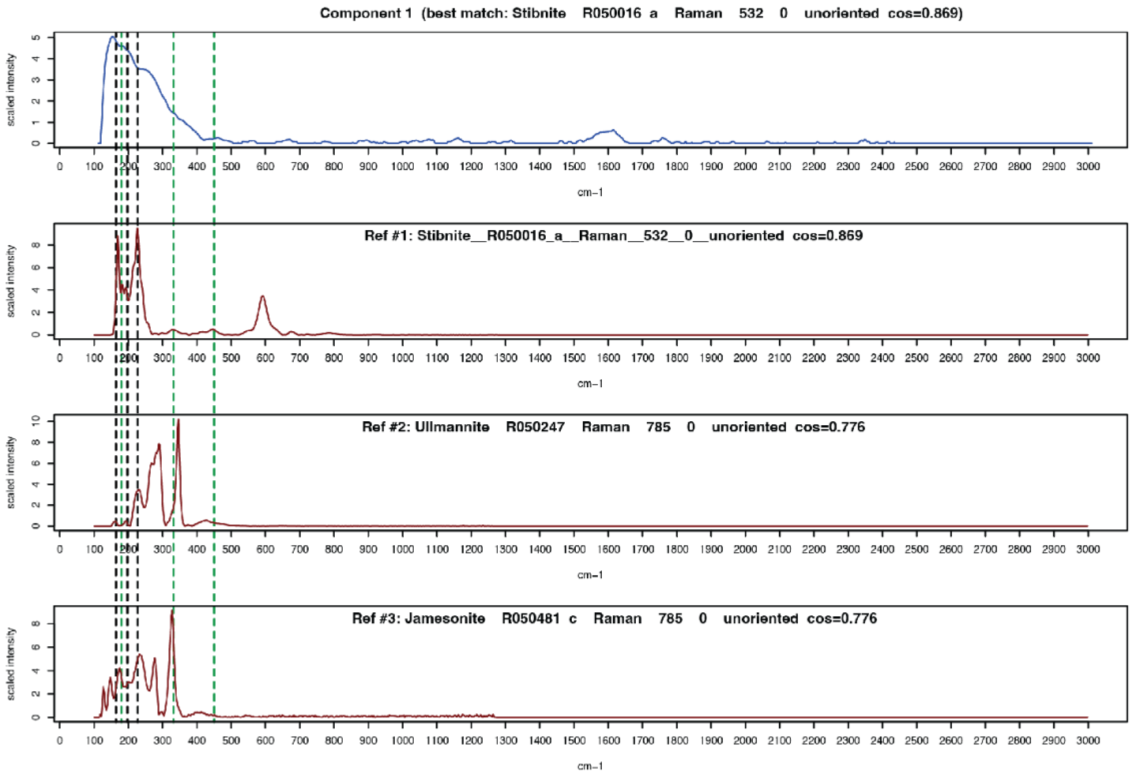
Component 1 (best match: Stibnite R050016 a Raman 532 0 unoriented cos=0.869)
Ref #1: Stibnite__R050016_a__Raman__532__0__unoriented cos=0.869
Ref #2: Ullmannite R050247 Raman 785 0 unoriented cos=0.776
Ref #3: Jamesonite R050481 c Raman 785 0 unoriented cos=0.776
scaled intensity
cm−1

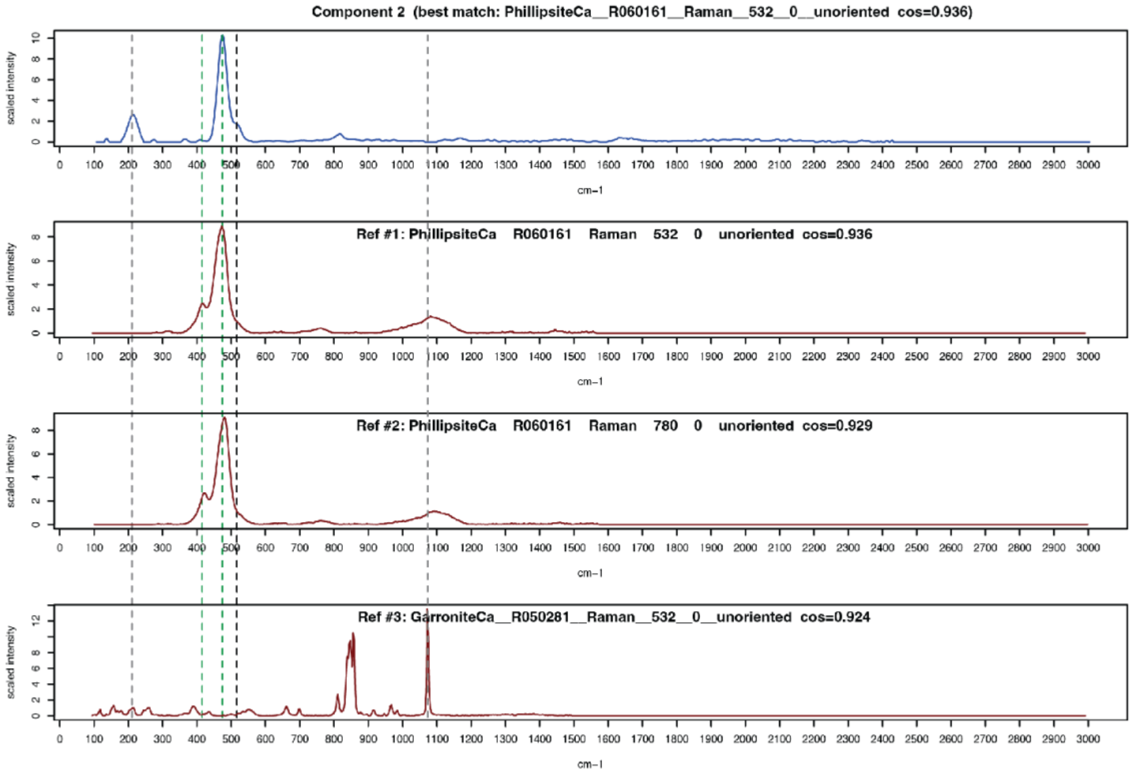
Component 2 (best match: PhillipsiteCa__R060161__Raman__532__0__unoriented cos=0.936)
Ref #1: PhillipsiteCa R060161 Raman 532 0 unoriented cos=0.936
Ref #2: PhillipsiteCa R060161 Raman 780 0 unoriented cos=0.929
Ref #3: GarroniteCa__R050281__Raman__532__0__unoriented cos=0.924
scaled intensity
cm−1

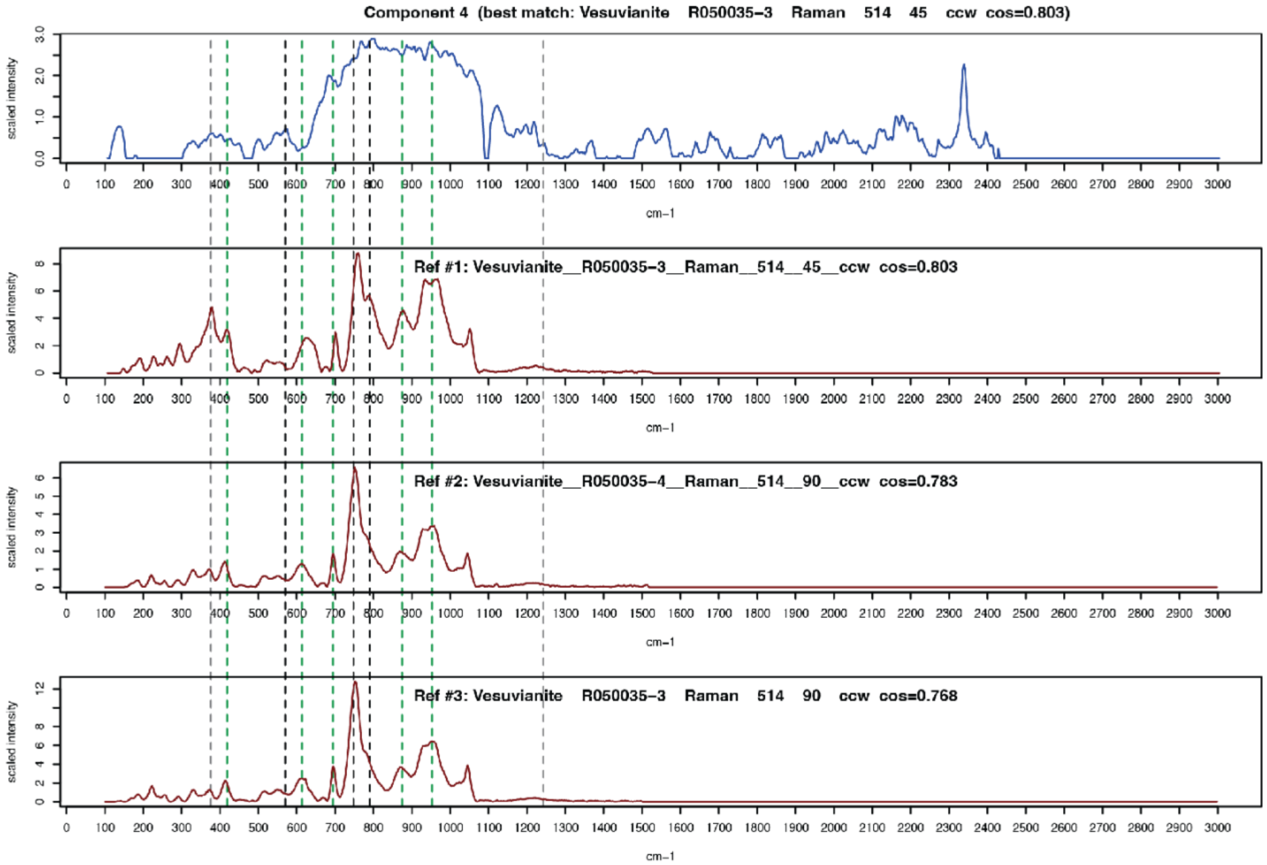
Component 4 (best match: Vesuvianite R050035-3 Raman 514 45 ccw cos=0.803)
Ref #1: Vesuvianite__R050035-3__Raman__514__45__ccw cos=0.803
Ref #2: Vesuvianite__R050035-4__Raman__514__90__ccw cos=0.783
Ref #3: Vesuvianite R050035-3 Raman 514 90 ccw cos=0.768
scaled intensity
cm−1
0 100 200 300 400 500 600 700 800 900 1000 1100 1200 1300 1400 1500 1600 1700 1800 1900 2000 2100 2200 2300 2400 2500 2600 2700 2800 2900 3000

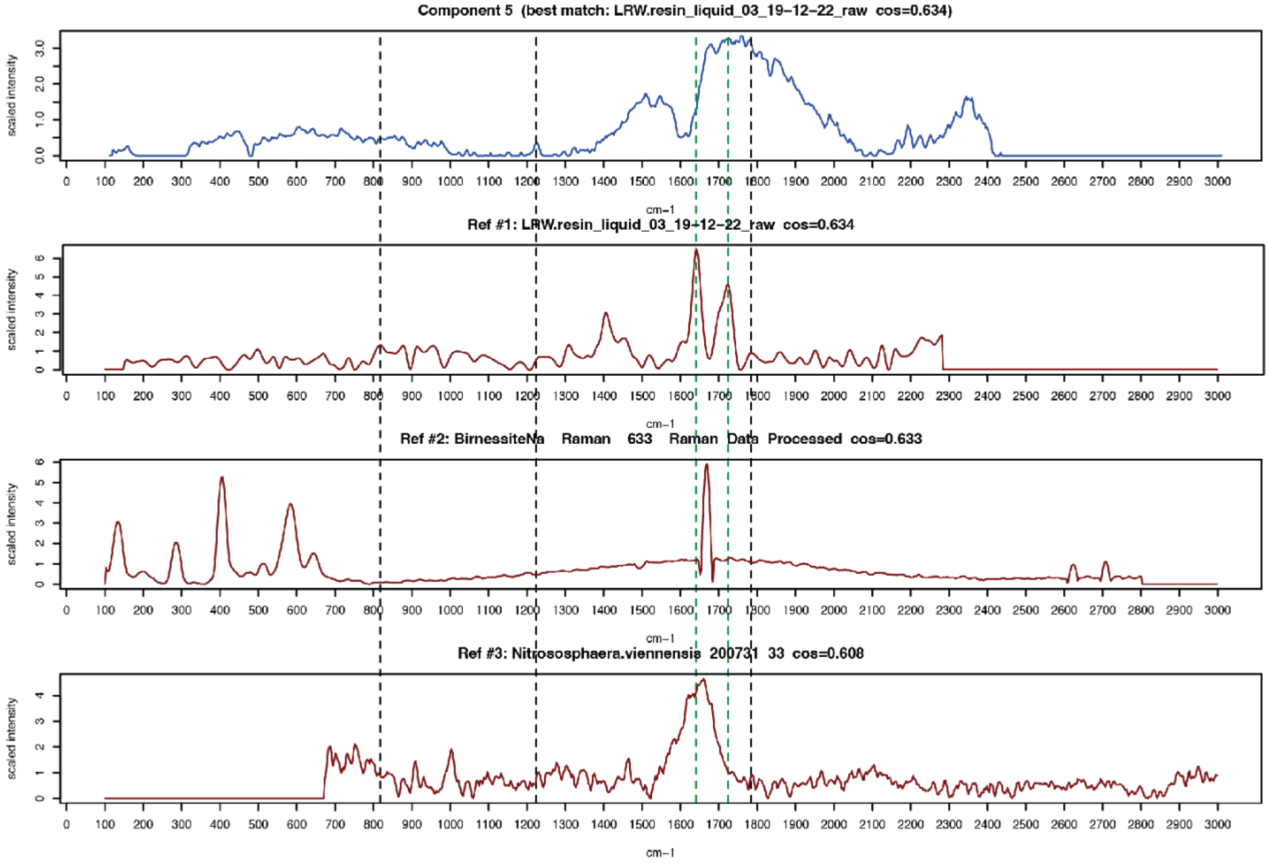
Component 5 (best match: LRW.resin_liquid_03_19-12-22_raw cos=0.634)
Ref #1: LRW.resin_liquid_03_19-12-22_raw cos=0.634
Ref #2: BirnessiteNa Raman 633 Raman Data Processed cos=0.633
Ref #3: Nitrososphaera.viennensis 200731 33 cos=0.608
scaled intensity
cm−1
0 100 200 300 400 500 600 700 800 900 1000 1100 1200 1300 1400 1500 1600 1700 1800 1900 2000 2100 2200 2300 2400 2500 2600 2700 2800 2900 3000

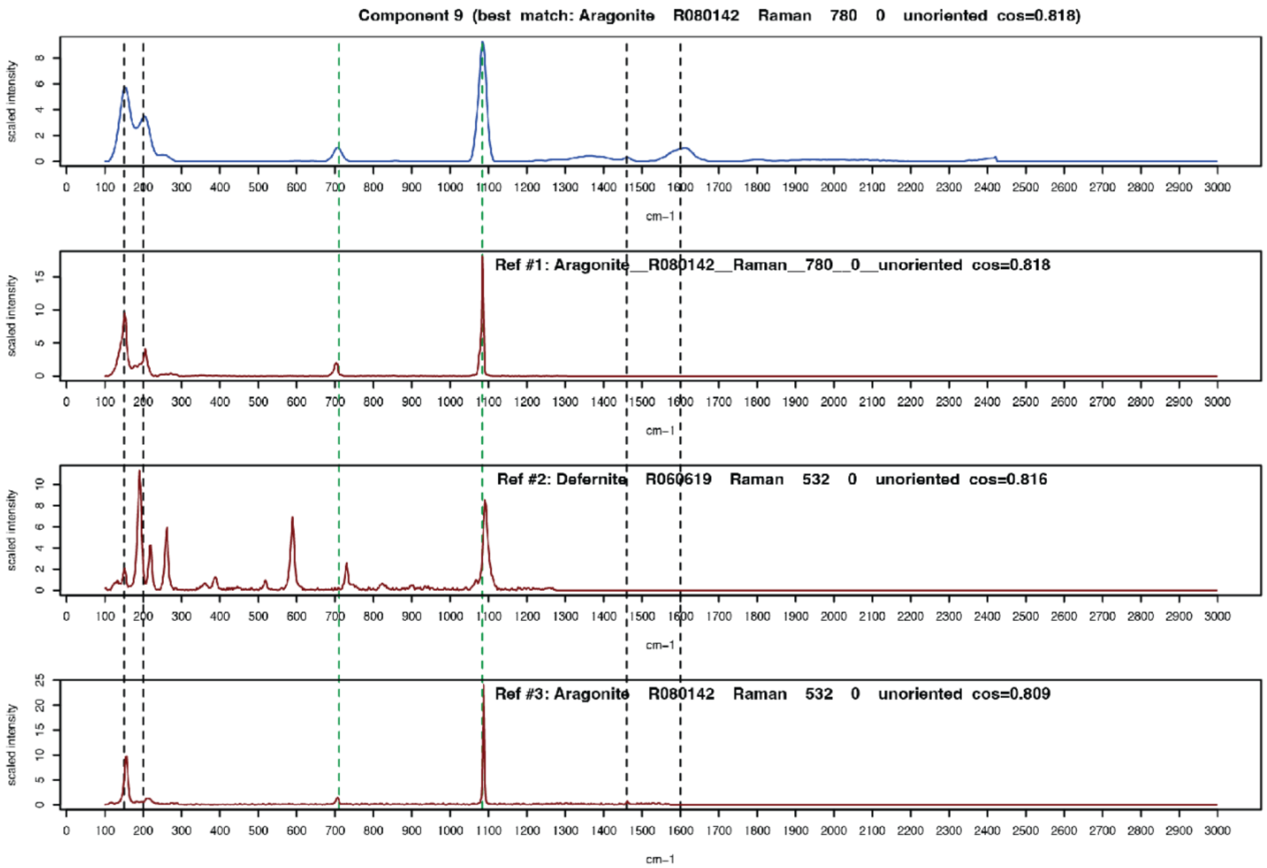
Component 9 (best match: Aragonite R080142 Raman 780 0 unoriented cos=0.818)
Ref #1: Aragonite__R080142__Raman__780__0__unoriented cos=0.818
Ref #2: Defernite R060619 Raman 532 0 unoriented cos=0.816
Ref #3: Aragonite R080142 Raman 532 0 unoriented cos=0.809
scaled intensity
cm−1

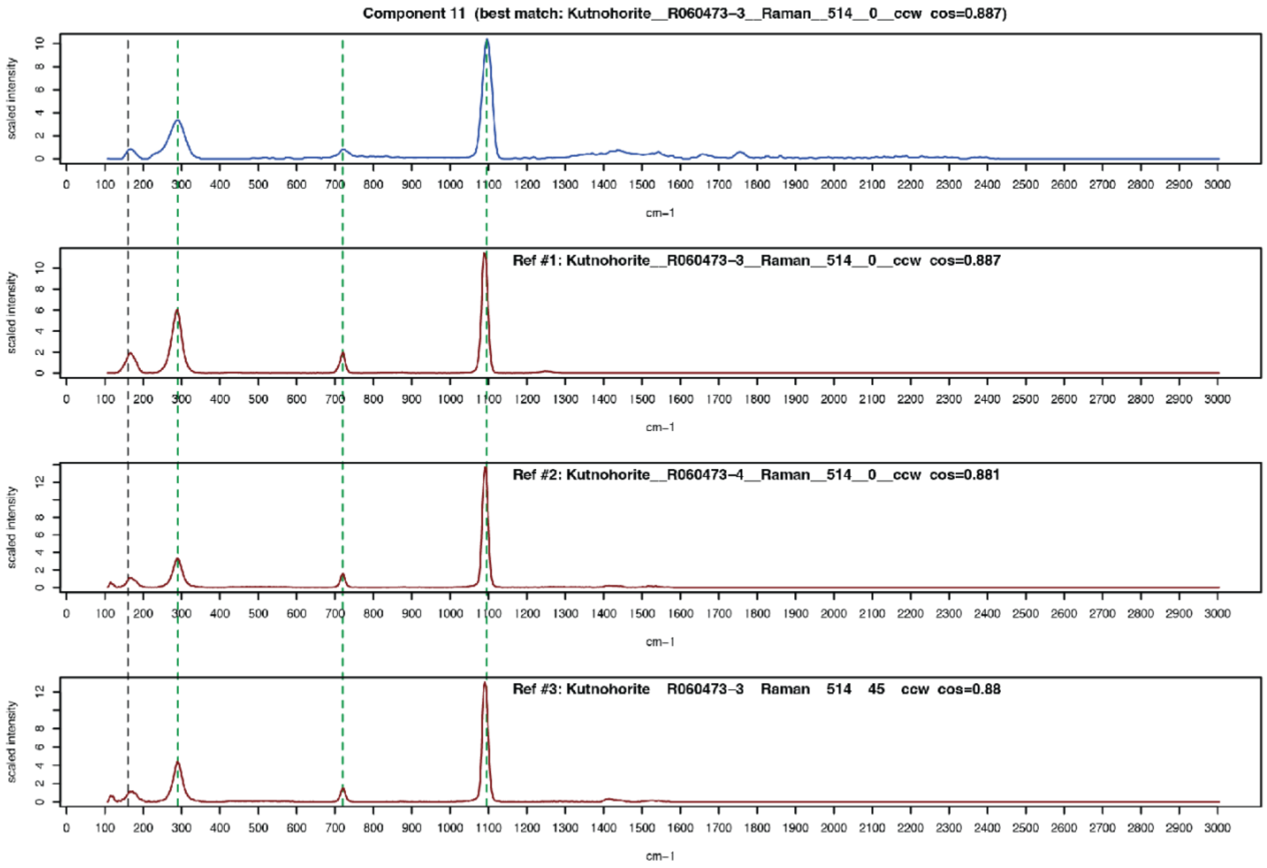
Component 11 (best match: Kutnohorite__R060473−3__Raman__514__0__ccw cos=0.887)
Ref #1: Kutnohorite__R060473−3__Raman__514__0__ccw cos=0.887
Ref #2: Kutnohorite__R060473−4__Raman__514__0__ccw cos=0.881
Ref #3: Kutnohorite R060473−3 Raman 514 45 ccw cos=0.88
scaled intensity
cm−1

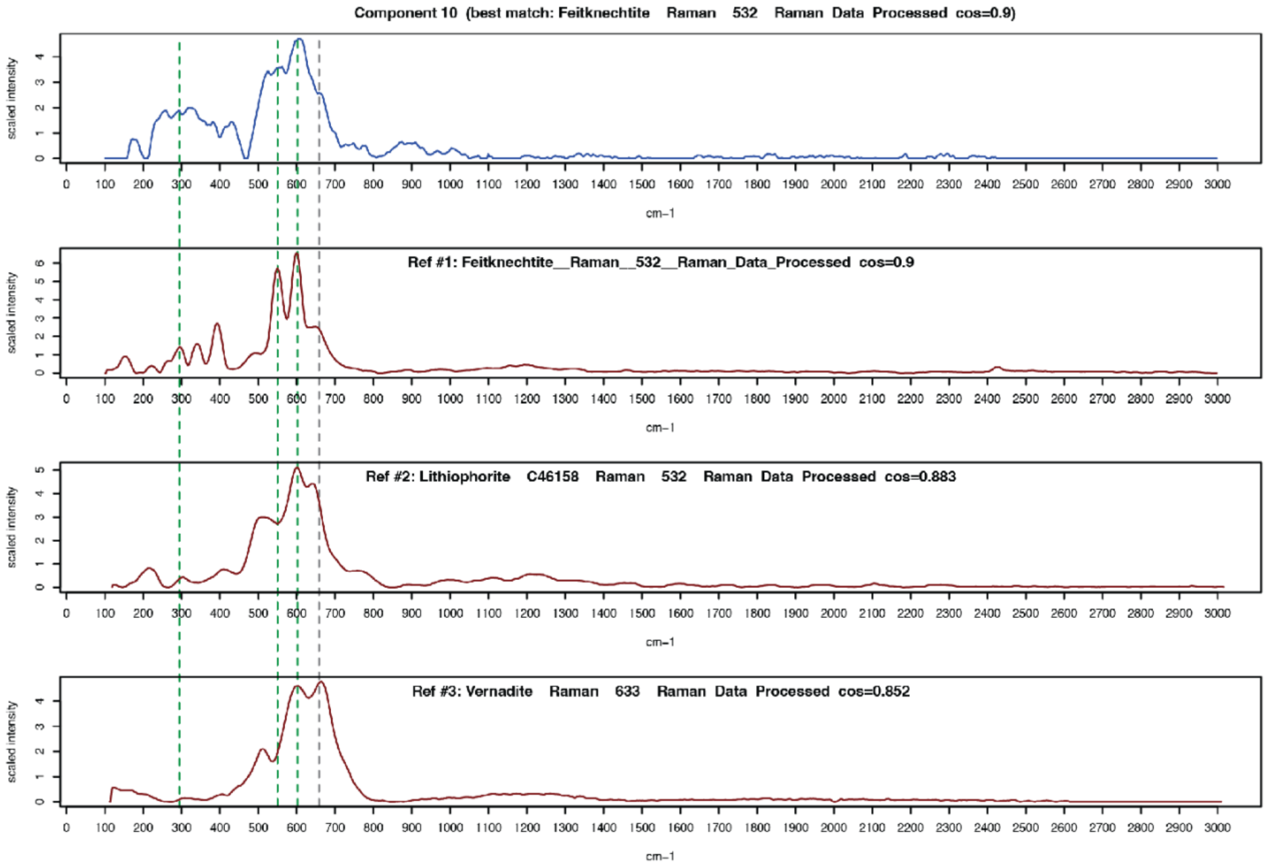
Component 10 (best match: Feitknechtite Raman 532 Raman Data Processed cos=0.9)
Ref #1: Feitknechtite__Raman__532__Raman_Data_Processed cos=0.9
Ref #2: Lithiophorite C46158 Raman 532 Raman Data Processed cos=0.883
Ref #3: Vernadite Raman 633 Raman Data Processed cos=0.852
scaled intensity
cm−1

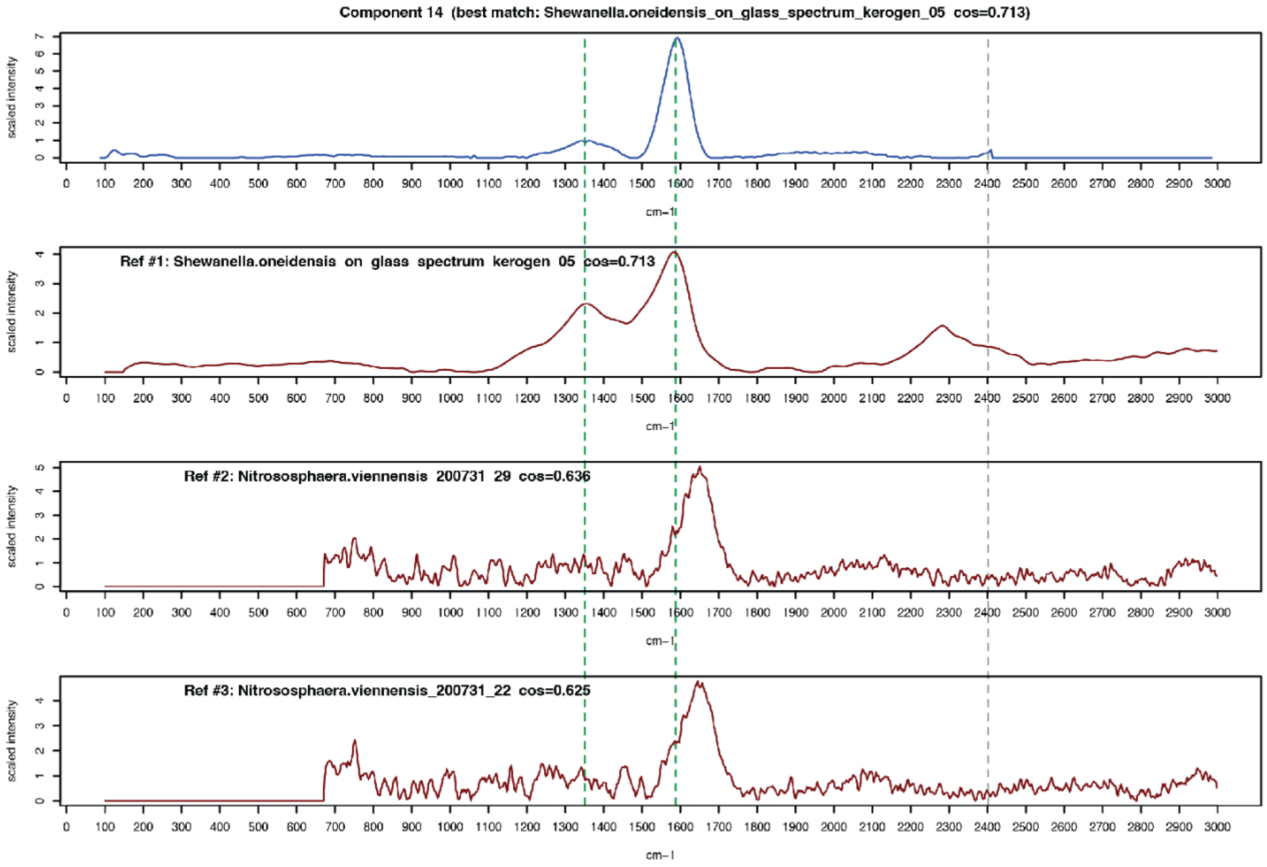
Component 14 (best match: Shewanella.oneidensis_on_glass_spectrum_kerogen_05 cos=0.713)
Ref #1: Shewanella.oneidensis on glass spectrum kerogen 05 cos=0.713
Ref #2: Nitrososphaera.viennensis 200731 29 cos=0.636
Ref #3: Nitrososphaera.viennensis_200731_22 cos=0.625
scaled intensity
cm−1

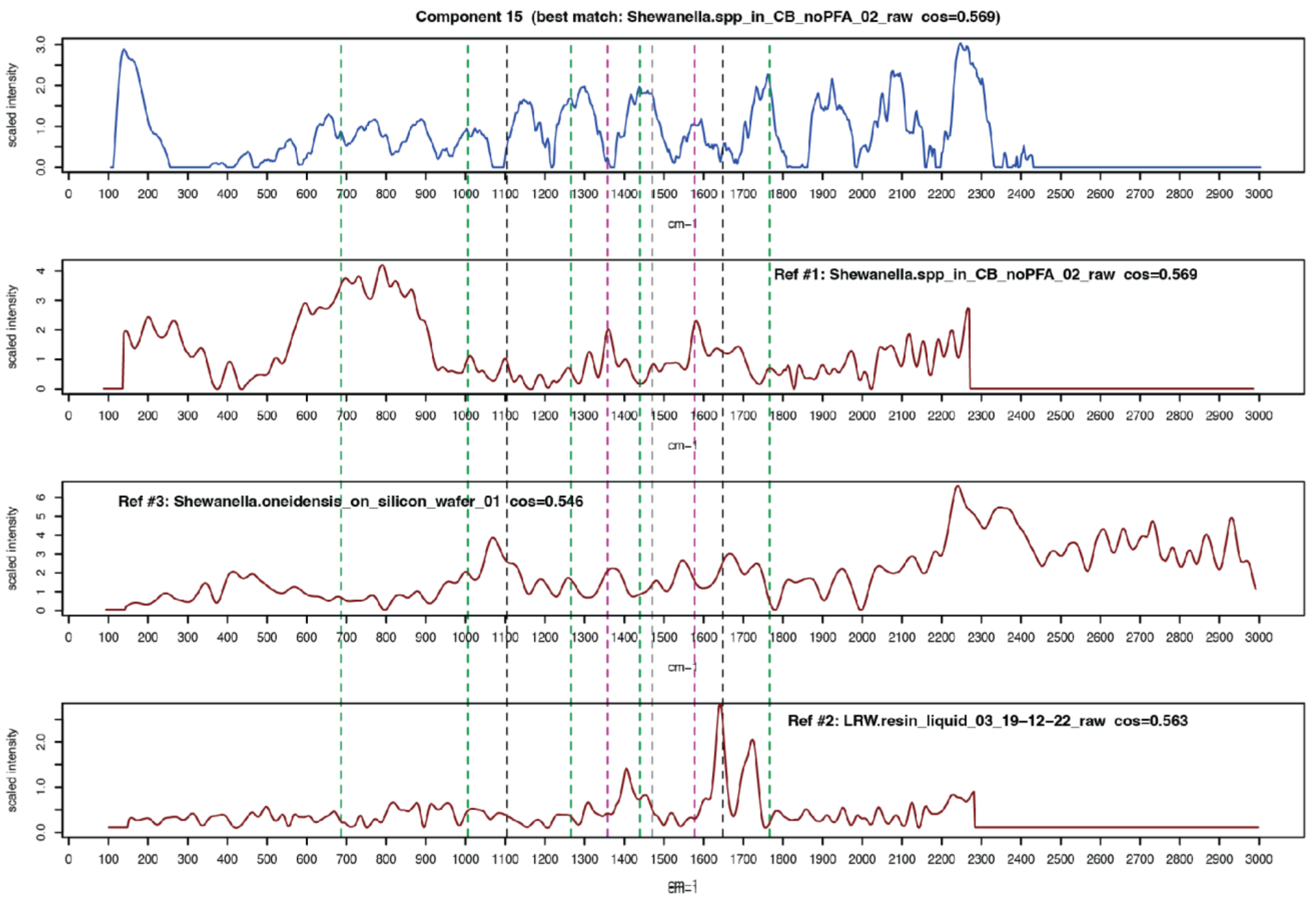
Component 15 (best match: Shewanella.spp_in_CB_noPFA_02_raw cos=0.569)
scaled intensity
cm-1
Ref #1: Shewanella.spp_in_CB_noPFA_02_raw cos=0.569
scaled intensity
cm-1
Ref #3: Shewanella.oneidensis_on_silicon_wafer_01 cos=0.546
scaled intensity
cm-1
Ref #2: LRW.resin_liquid_03_19–12–22_raw cos=0.563
scaled intensity
cm-1

Fig. S2. Manual peak matching for the PN components. The top row for each component is the component's spectrum, while rows 2-4 are the spectra of the top three matches from our expanded reference database.

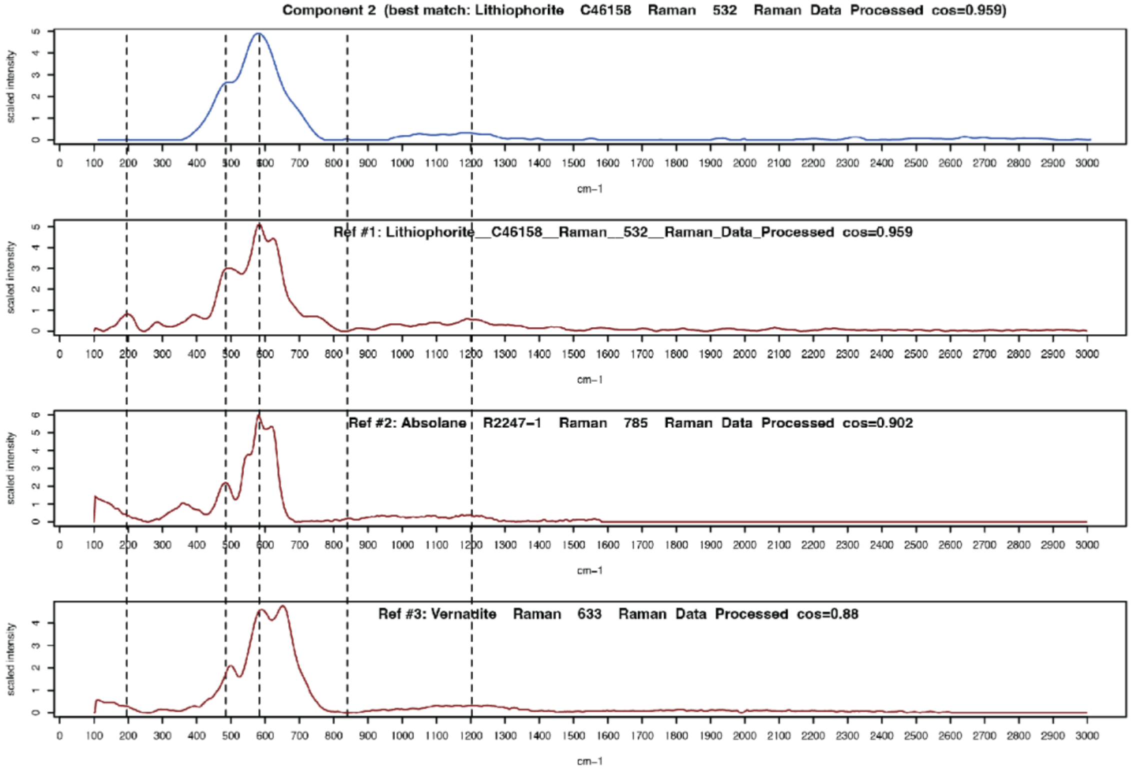

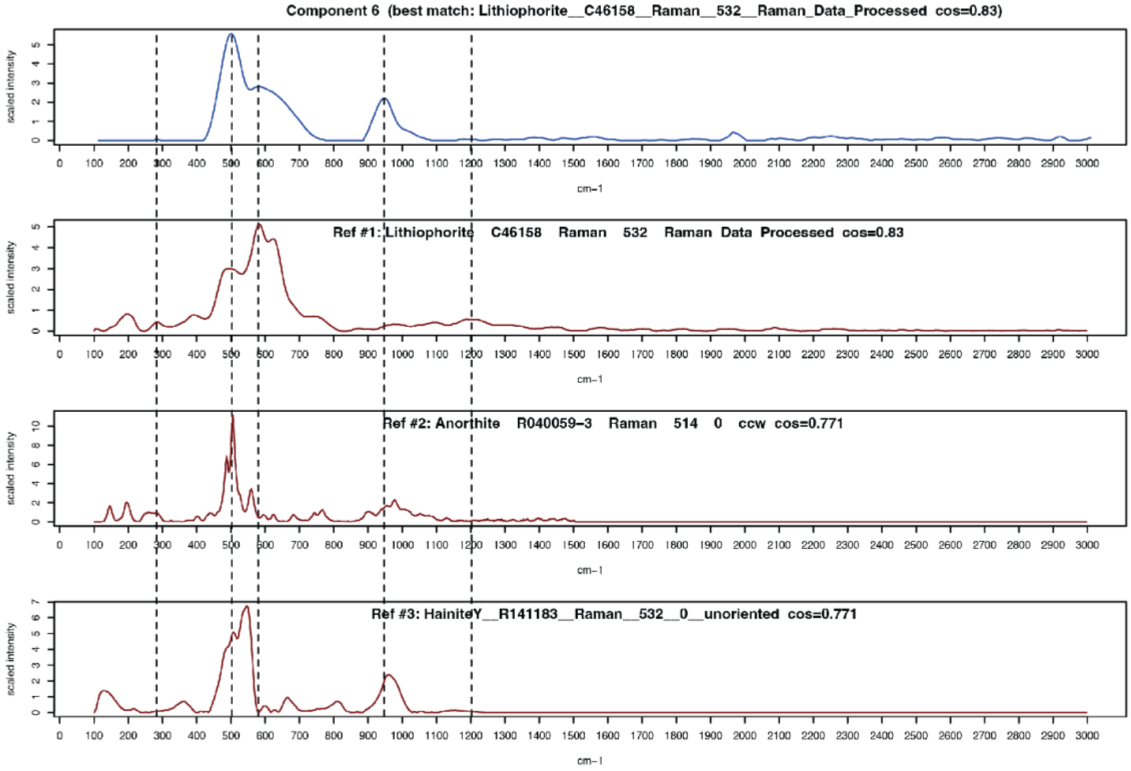

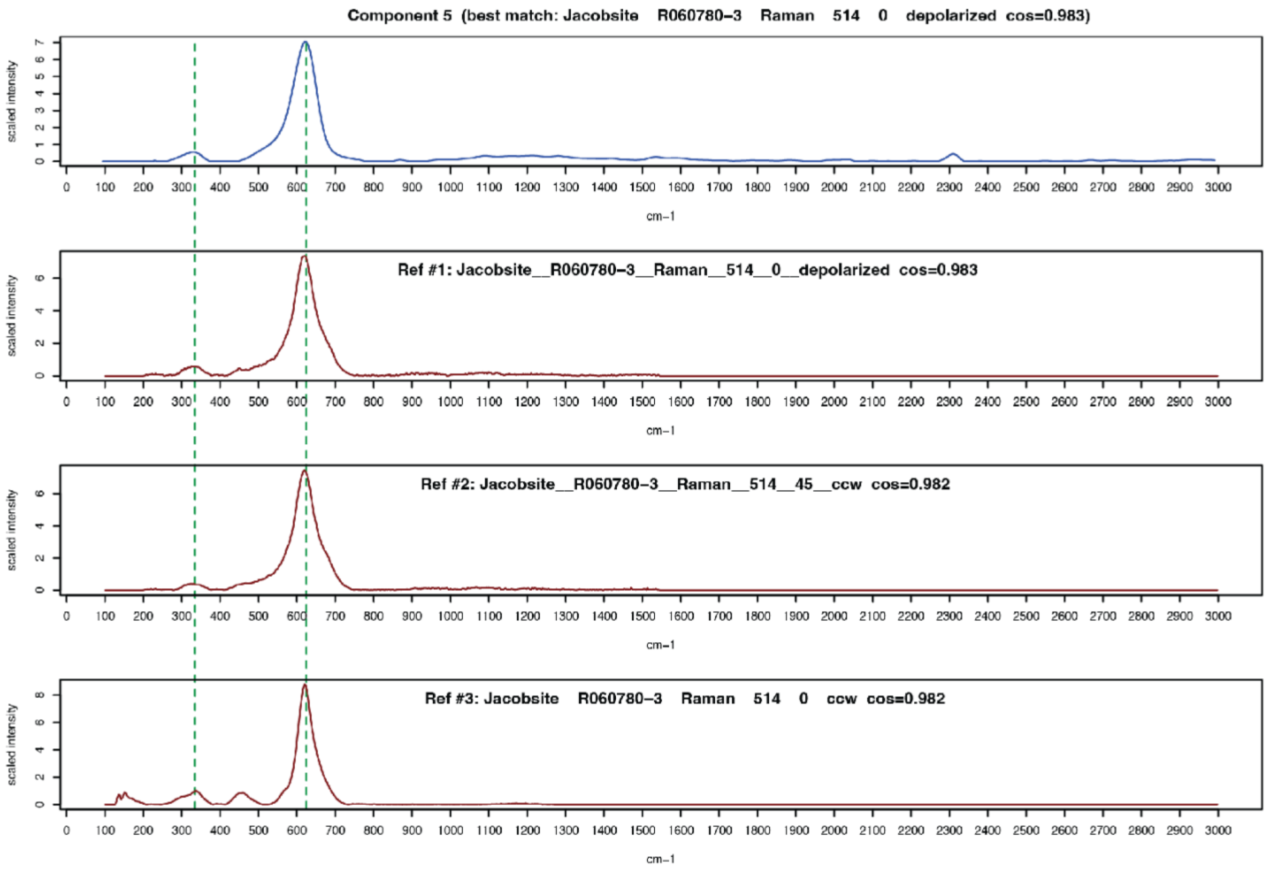

Component 5 (best match: Jacobsite R060780-3 Raman 514 0 depolarized cos=0.983)
Ref #1: Jacobsite__R060780-3__Raman__514__0__depolarized cos=0.983
Ref #2: Jacobsite__R060780-3__Raman__514__45__ccw cos=0.982
Ref #3: Jacobsite R060780-3 Raman 514 0 ccw cos=0.982
scaled intensity
cm−1

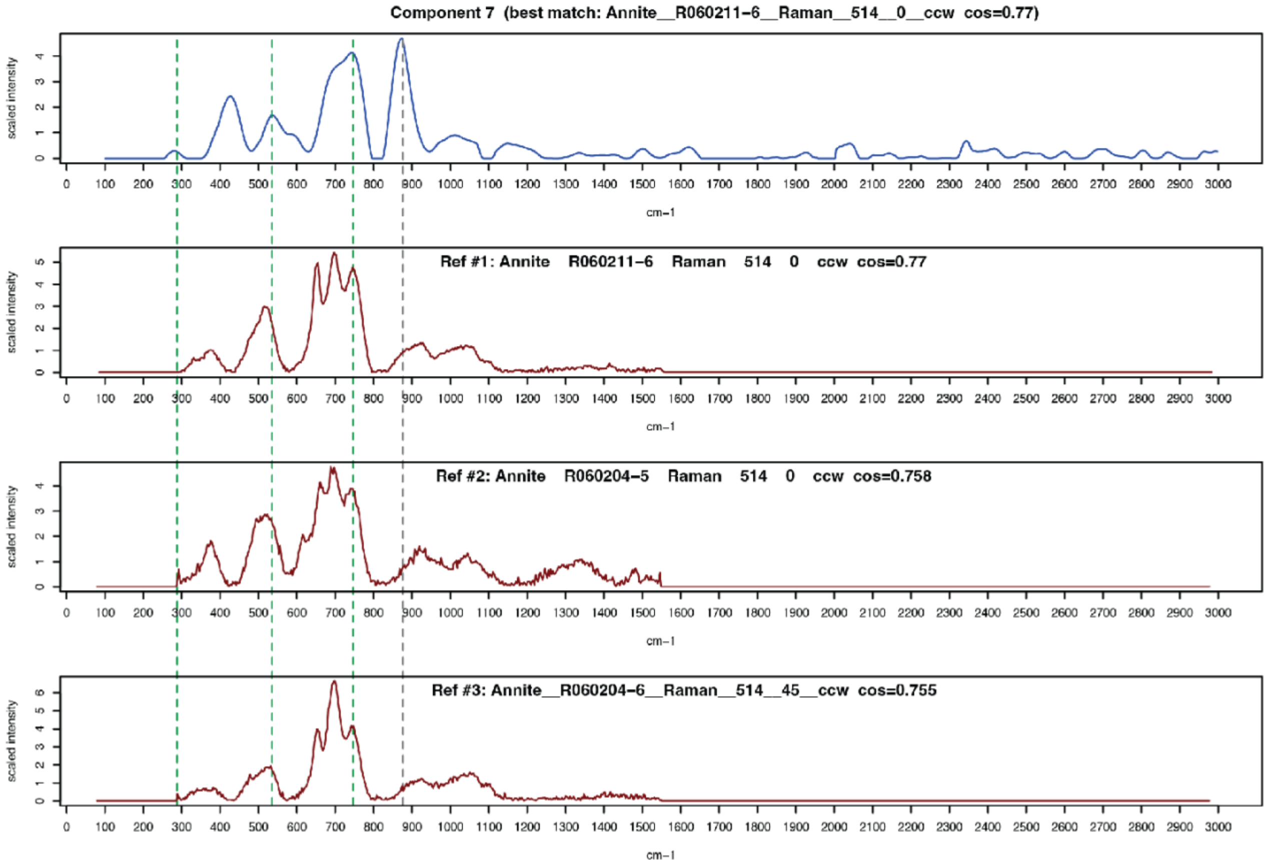

Component 7 (best match: Annite__R060211-6__Raman__514__0__ccw cos=0.77)
Ref #1: Annite R060211-6 Raman 514 0 ccw cos=0.77
Ref #2: Annite R060204-5 Raman 514 0 ccw cos=0.758
Ref #3: Annite__R060204-6__Raman__514__45__ccw cos=0.755
scaled intensity
cm−1

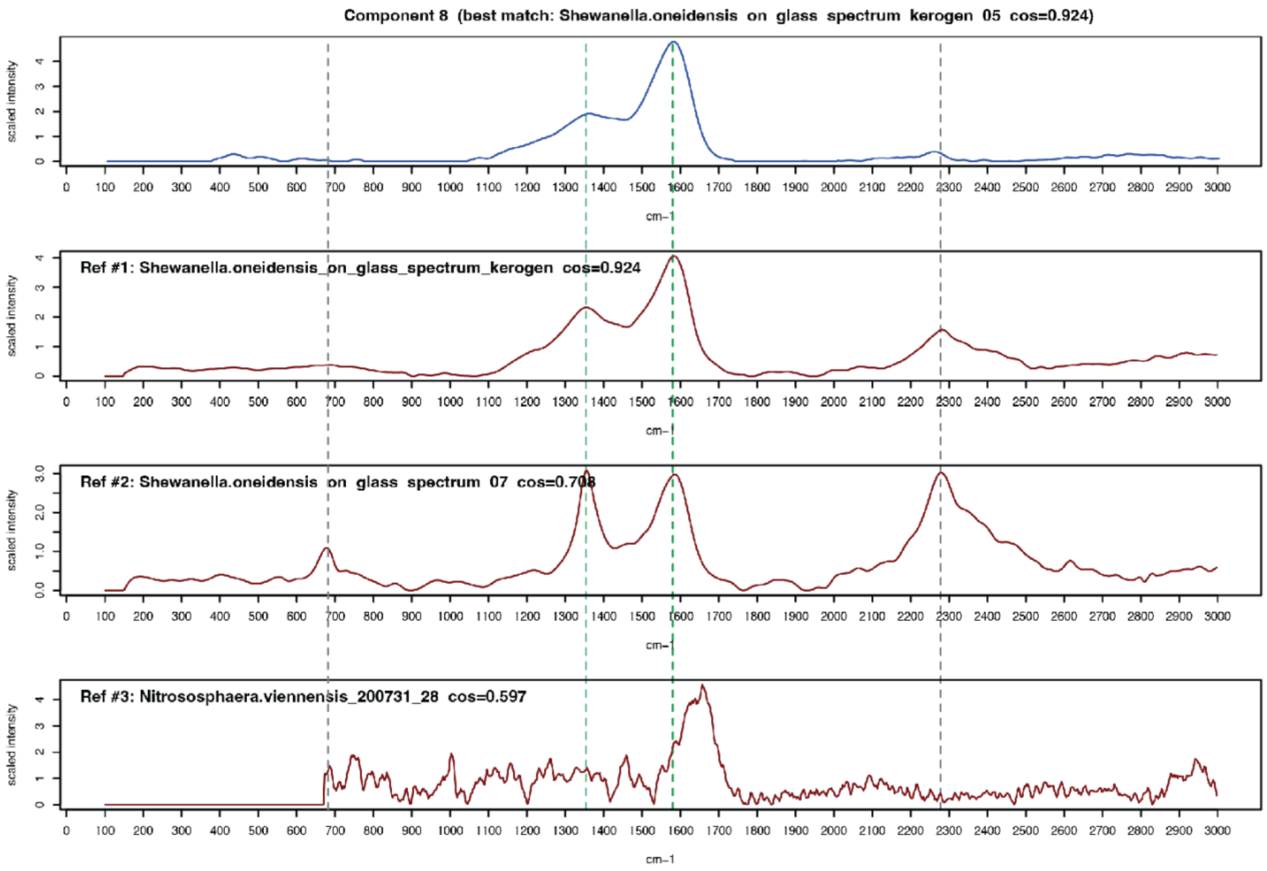
Component 8 (best match: Shewanella.oneidensis on glass spectrum kerogen 05 cos=0.924)
Ref #1: Shewanella.oneidensis_on_glass_spectrum_kerogen cos=0.924
Ref #2: Shewanella.oneidensis on glass spectrum 07 cos=0.708
Ref #3: Nitrososphaera.viennensis_200731_28 cos=0.597
scaled intensity
cm−1

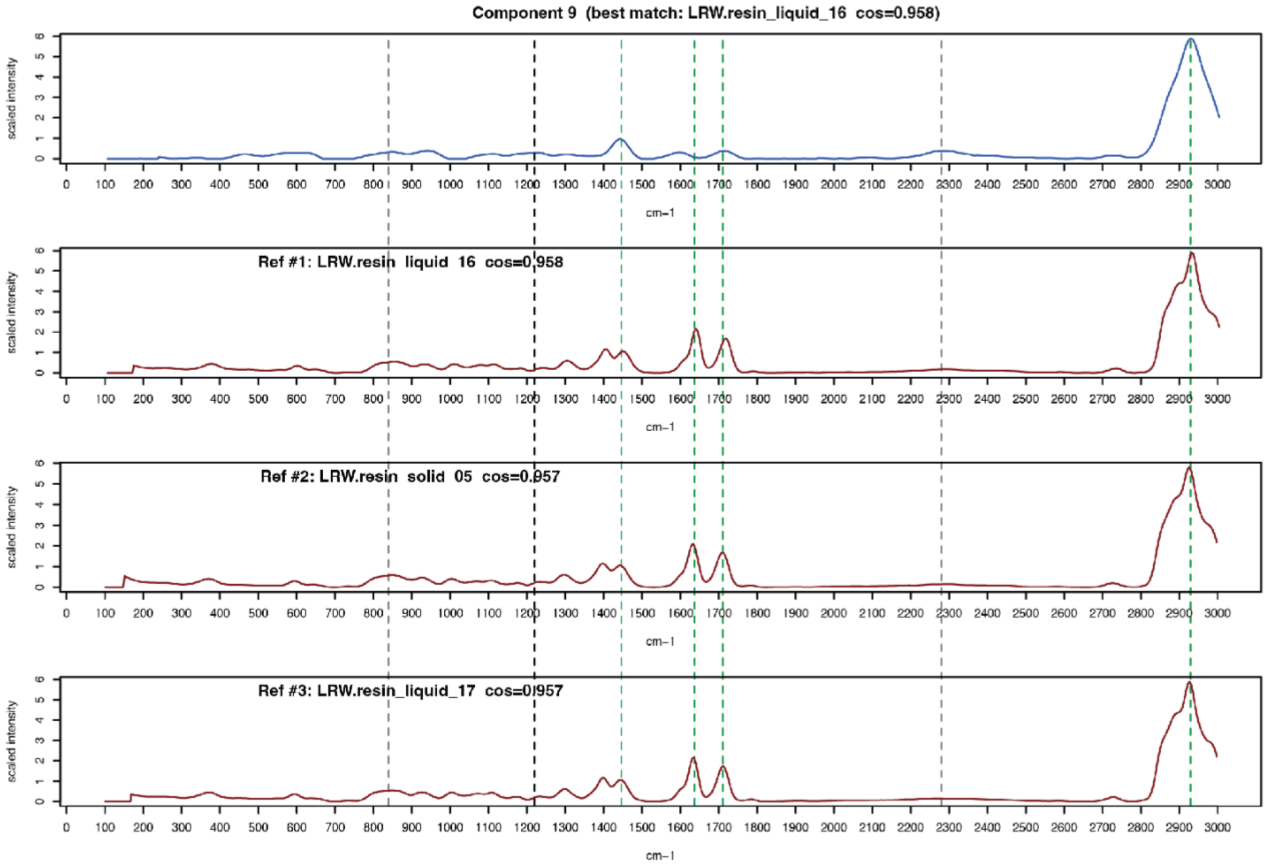
Component 9 (best match: LRW.resin_liquid_16 cos=0.958)
Ref #1: LRW.resin liquid 16 cos=0.958
Ref #2: LRW.resin solid 05 cos=0.957
Ref #3: LRW.resin_liquid_17 cos=0.957
scaled intensity
cm−1

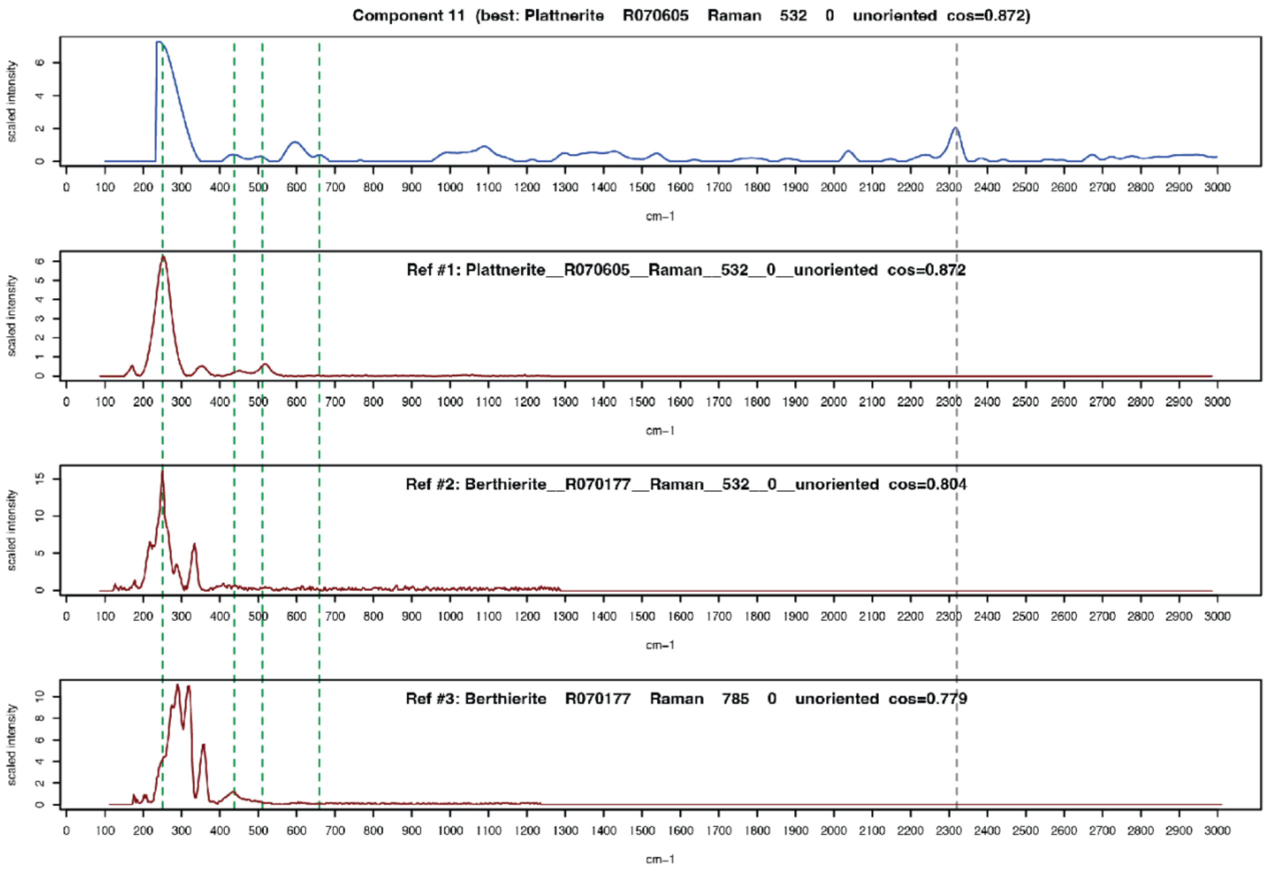
Component 11 (best: Plattnerite R070605 Raman 532 0 unoriented cos=0.872)
Ref #1: Plattnerite__R070605__Raman__532__0__unoriented cos=0.872
Ref #2: Berthierite__R070177__Raman__532__0__unoriented cos=0.804
Ref #3: Berthierite R070177 Raman 785 0 unoriented cos=0.779
scaled intensity
cm−1

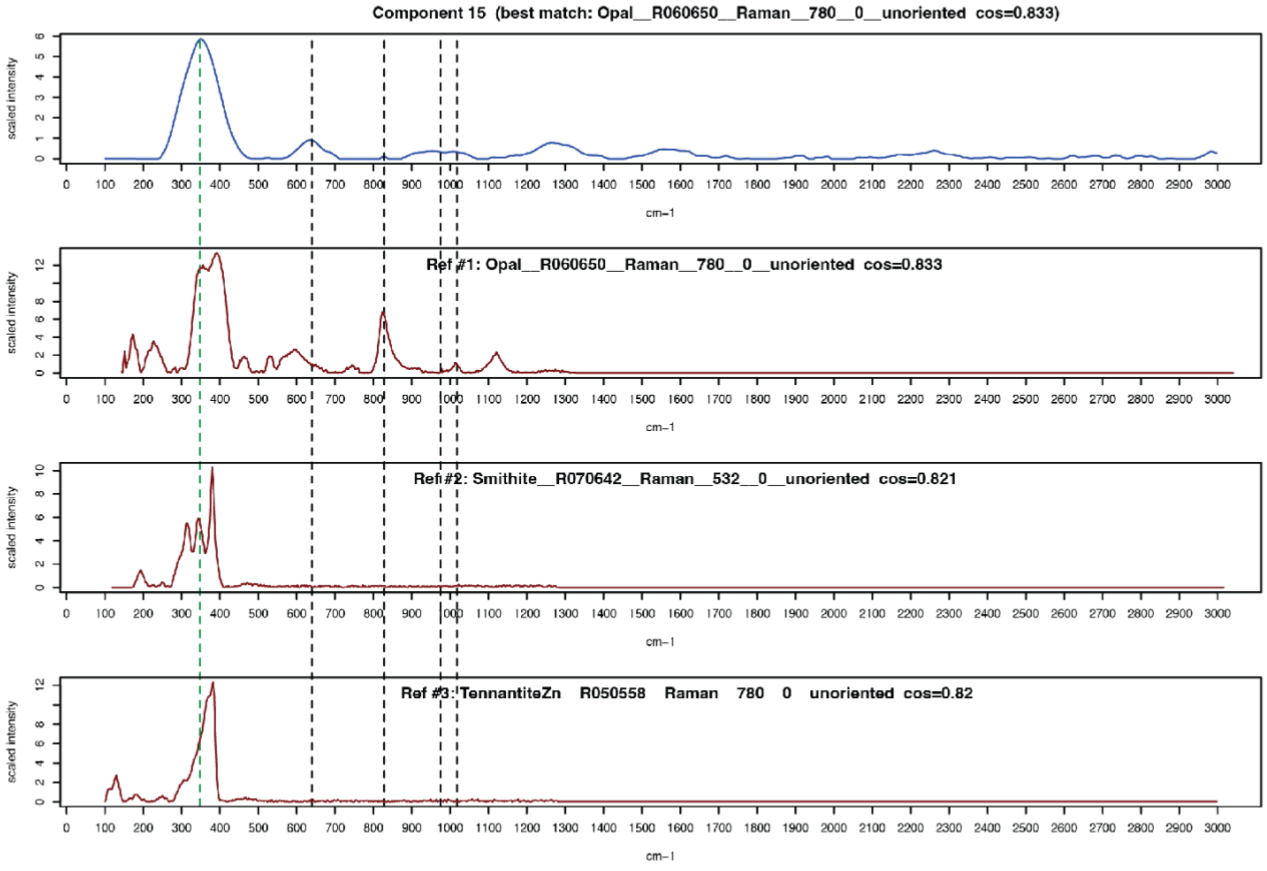
Component 15 (best match: Opal__R060650__Raman__780__0__unoriented cos=0.833)
Ref #1: Opal__R060650__Raman__780__0__unoriented cos=0.833
Ref #2: Smithite__R070642__Raman__532__0__unoriented cos=0.821
Ref #3: TennantiteZn R050558 Raman 780 0 unoriented cos=0.82
scaled intensity
cm−1

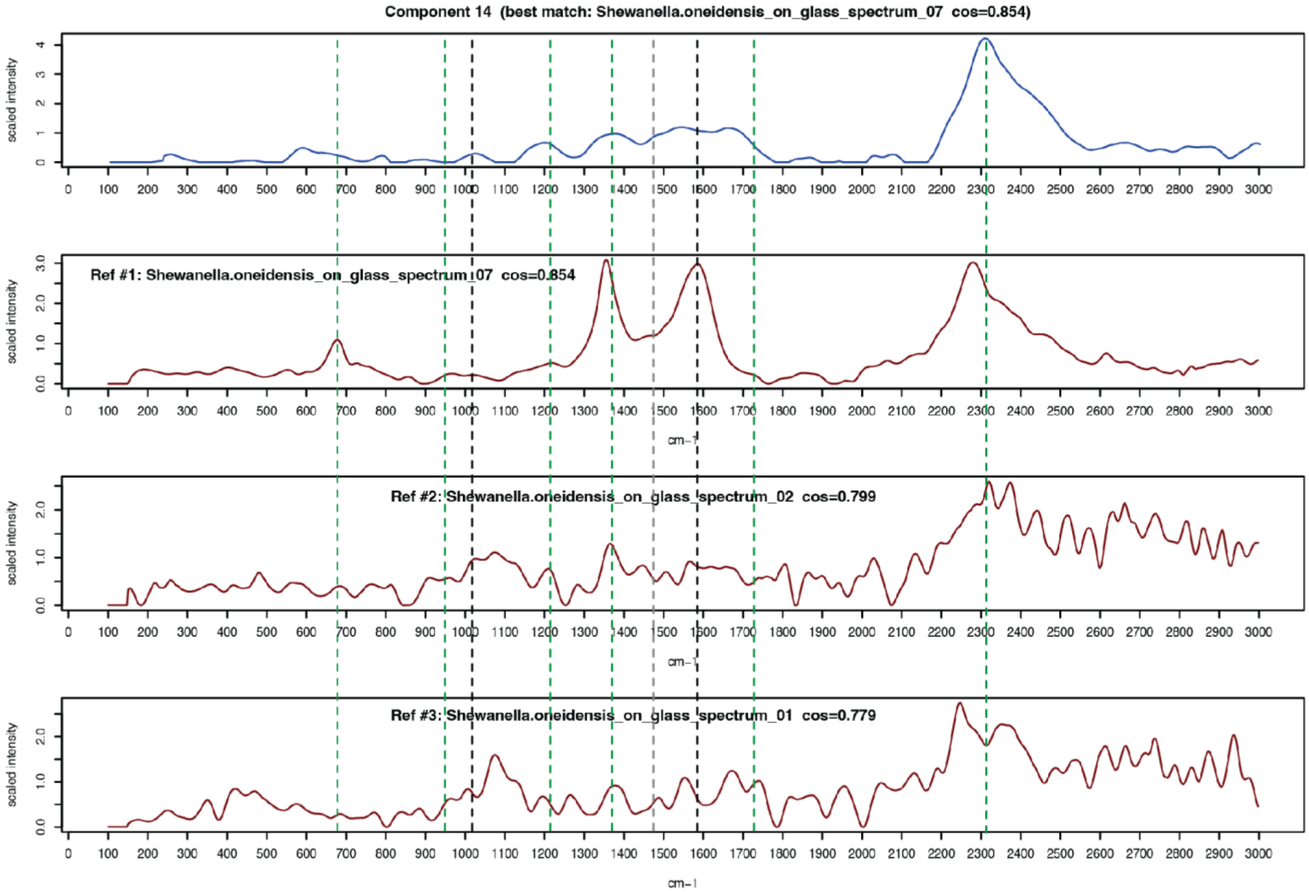

Component 14 (best match: Shewanella.oneidensis_on_glass_spectrum_07 cos=0.854)
Ref #1: Shewanella.oneidensis_on_glass_spectrum_07 cos=0.854
Ref #2: Shewanella.oneidensis_on_glass_spectrum_02 cos=0.799
Ref #3: Shewanella.oneidensis_on_glass_spectrum_01 cos=0.779
scaled intensity
cm-1

Fig. S3. Manual peak matching for the VB components. The top row for each component is the component's spectrum, while rows 2-4 are the spectra of the top three matches from our expanded reference database.

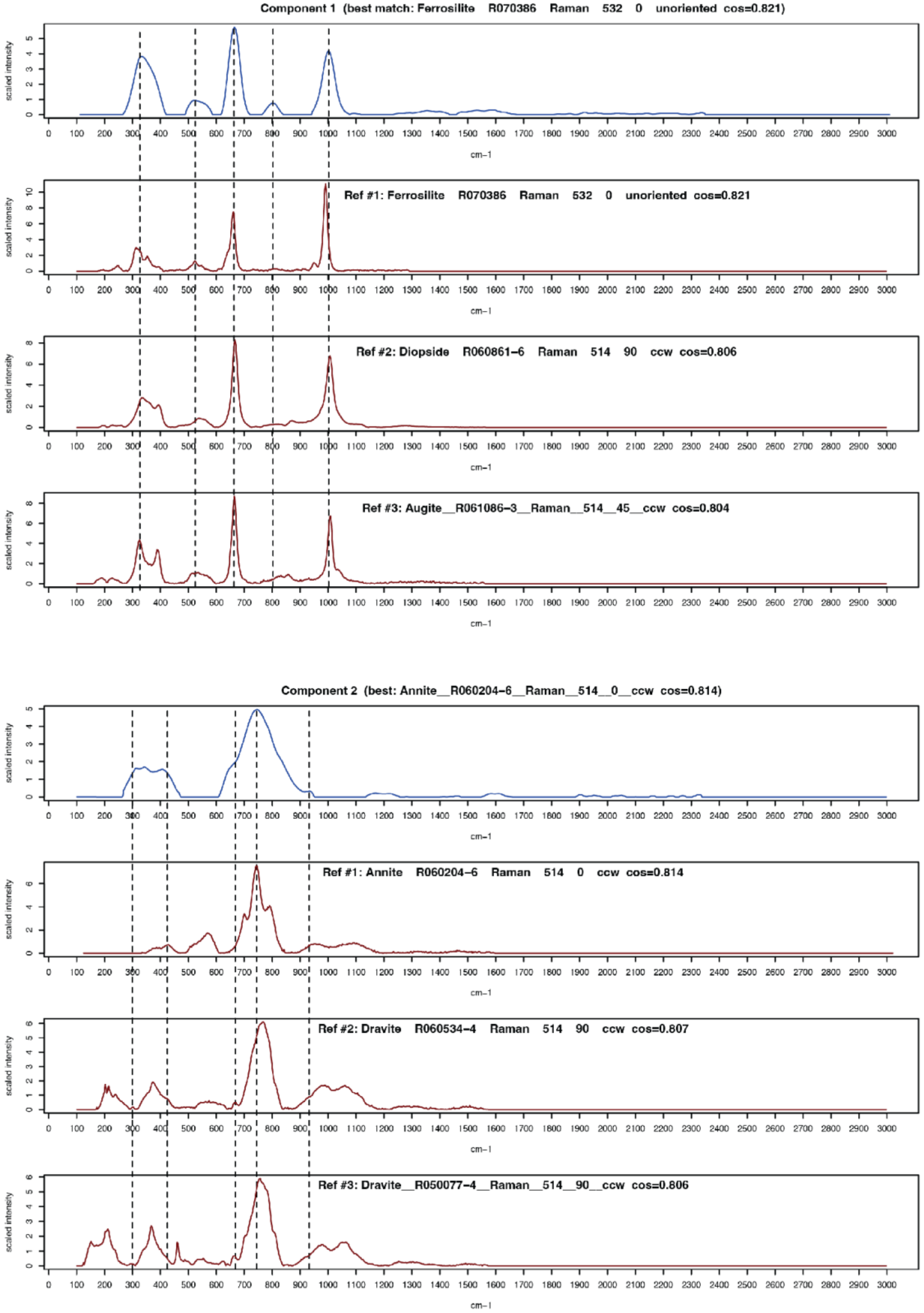

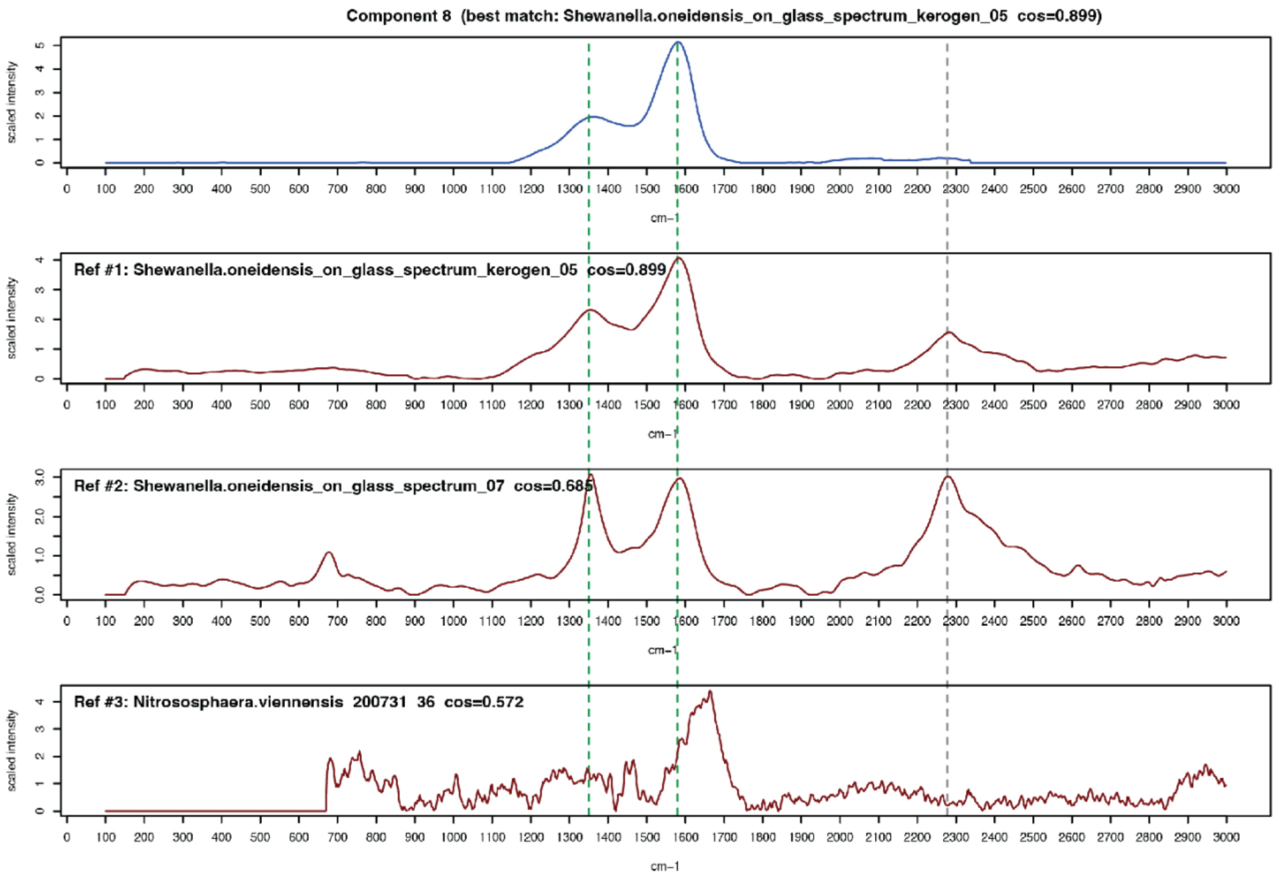
Component 8 (best match: Shewanella.oneidensis_on_glass_spectrum_kerogen_05 cos=0.899)
Ref #1: Shewanella.oneidensis_on_glass_spectrum_kerogen_05 cos=0.899
Ref #2: Shewanella.oneidensis_on_glass_spectrum_07 cos=0.685
Ref #3: Nitrososphaera.viennensis 200731 36 cos=0.572
scaled intensity
cm−1

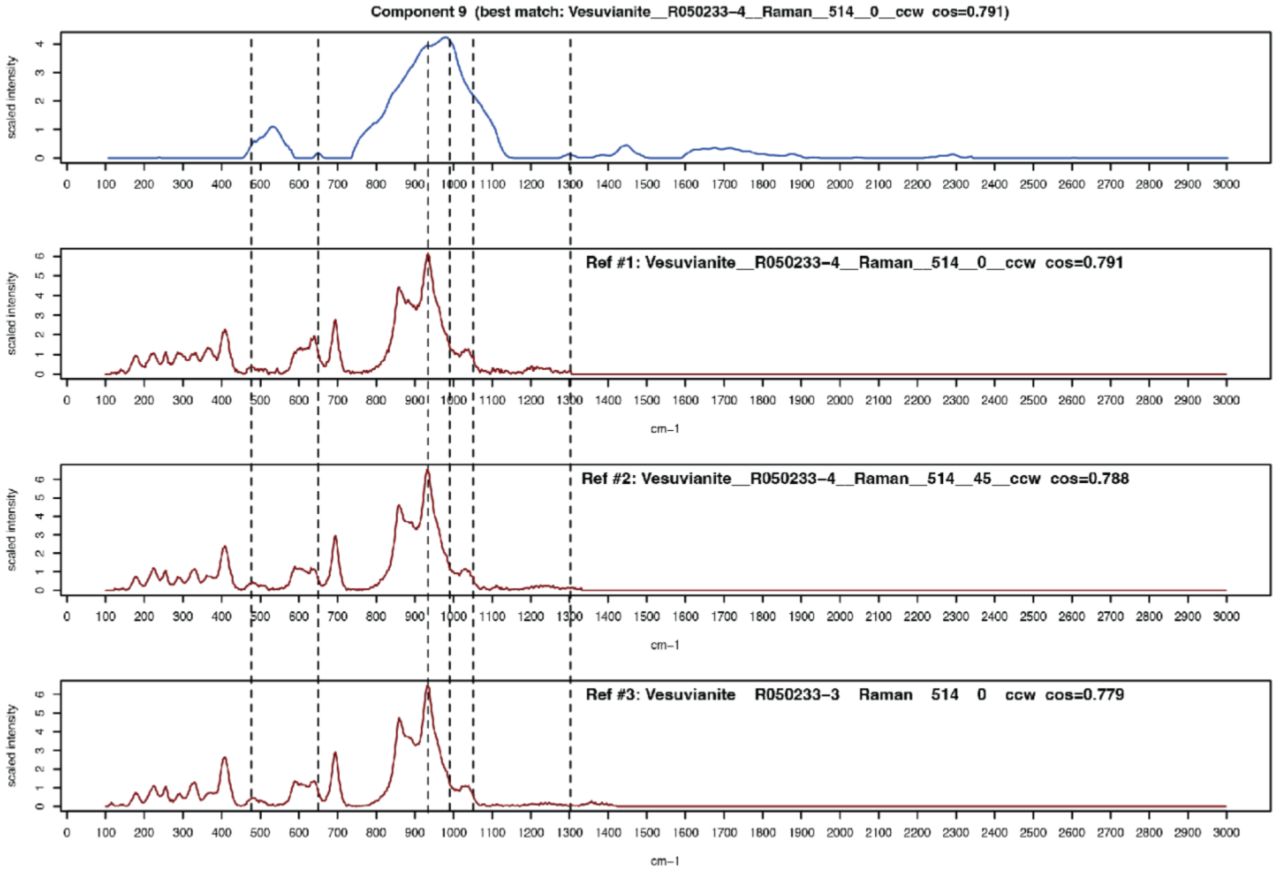
Component 9 (best match: Vesuvianite__R050233-4__Raman__514__0__ccw cos=0.791)
Ref #1: Vesuvianite__R050233-4__Raman__514__0__ccw cos=0.791
Ref #2: Vesuvianite__R050233-4__Raman__514__45__ccw cos=0.788
Ref #3: Vesuvianite R050233-3 Raman 514 0 ccw cos=0.779
scaled intensity
cm−1

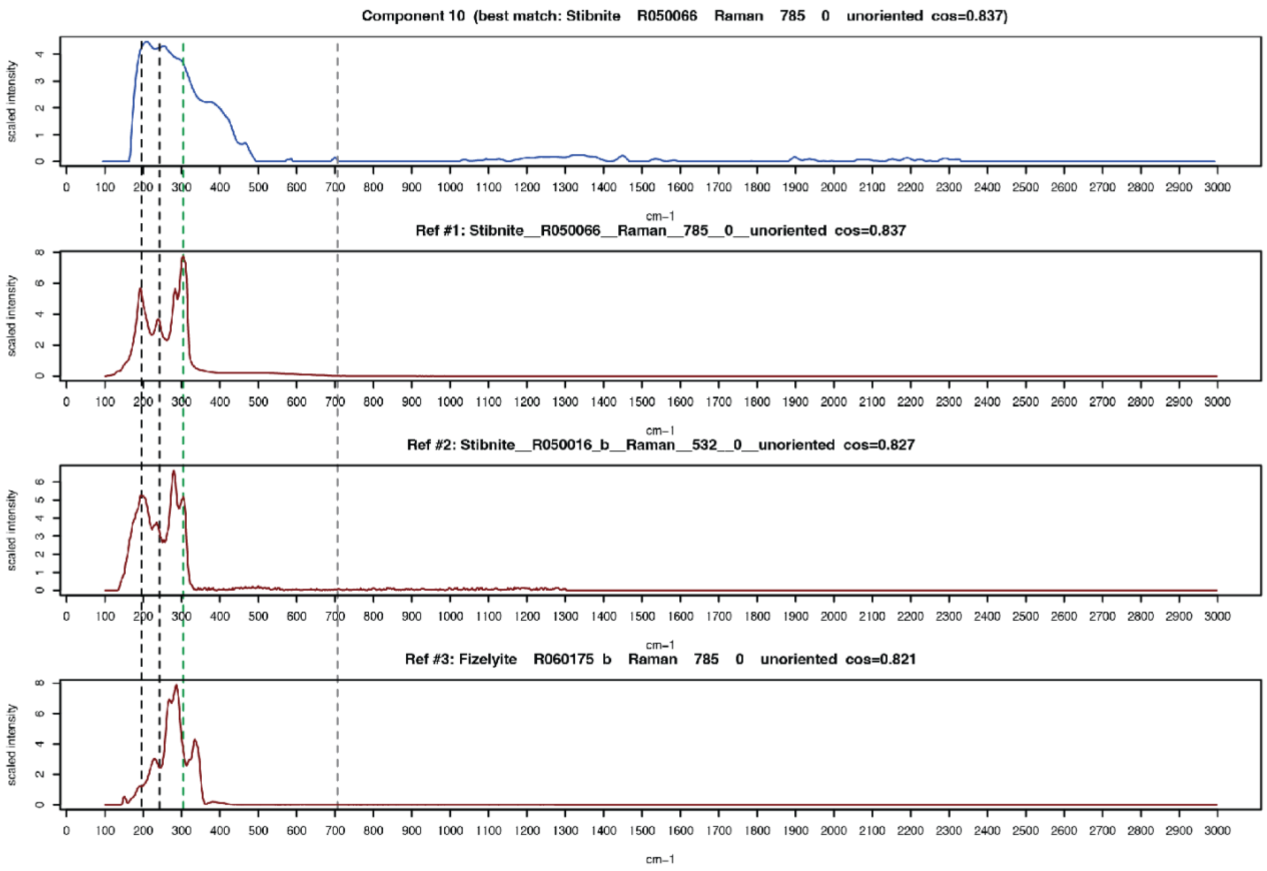
Component 10  (best match: Stibnite    R050066    Raman    785    0    unoriented  cos=0.837)
Ref #1: Stibnite__R050066__Raman__785__0__unoriented  cos=0.837
Ref #2: Stibnite__R050016_b__Raman__532__0__unoriented  cos=0.827
Ref #3: Fizelyite    R060175  b    Raman    785    0    unoriented  cos=0.821
scaled intensity
cm−1

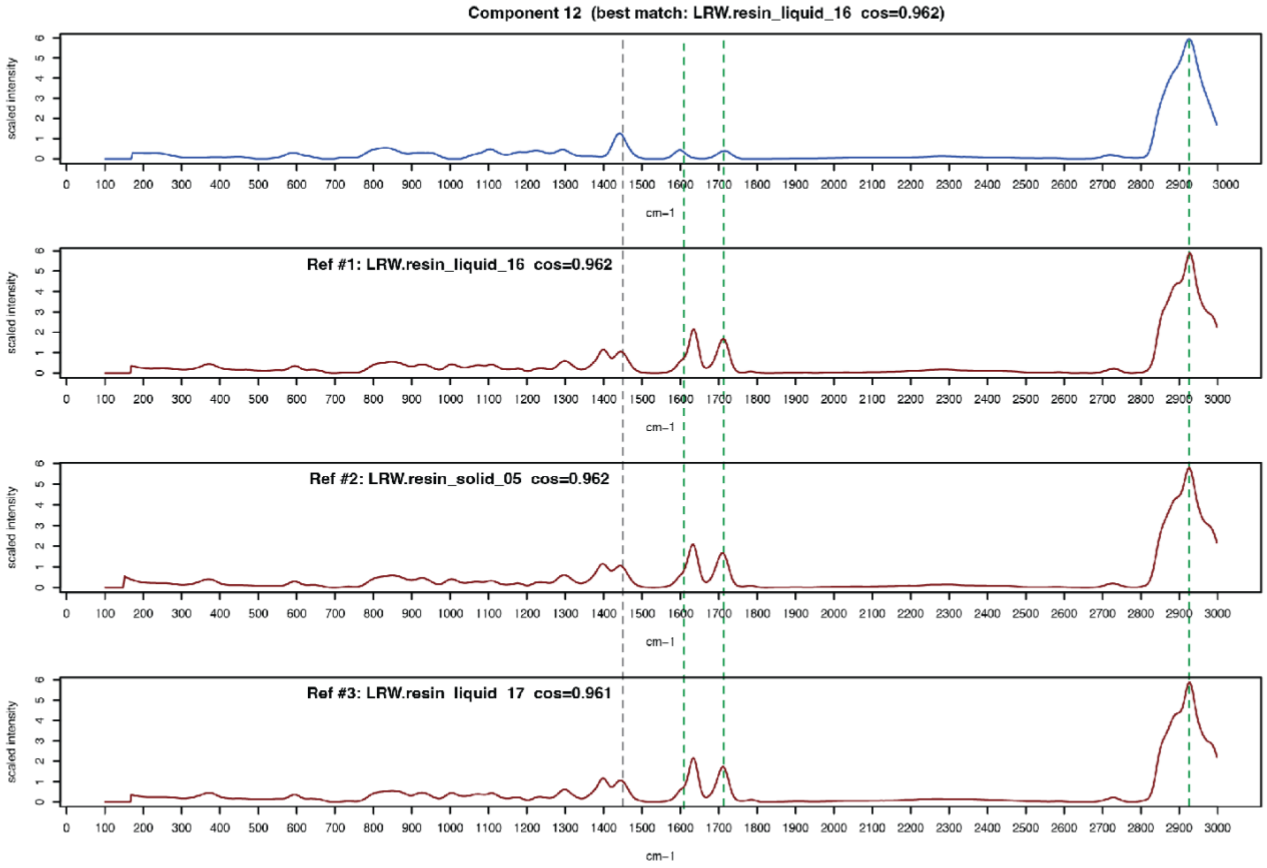
Component 12  (best match: LRW.resin_liquid_16  cos=0.962)
Ref #1: LRW.resin_liquid_16  cos=0.962
Ref #2: LRW.resin_solid_05  cos=0.962
Ref #3: LRW.resin  liquid  17  cos=0.961
scaled intensity
cm−1

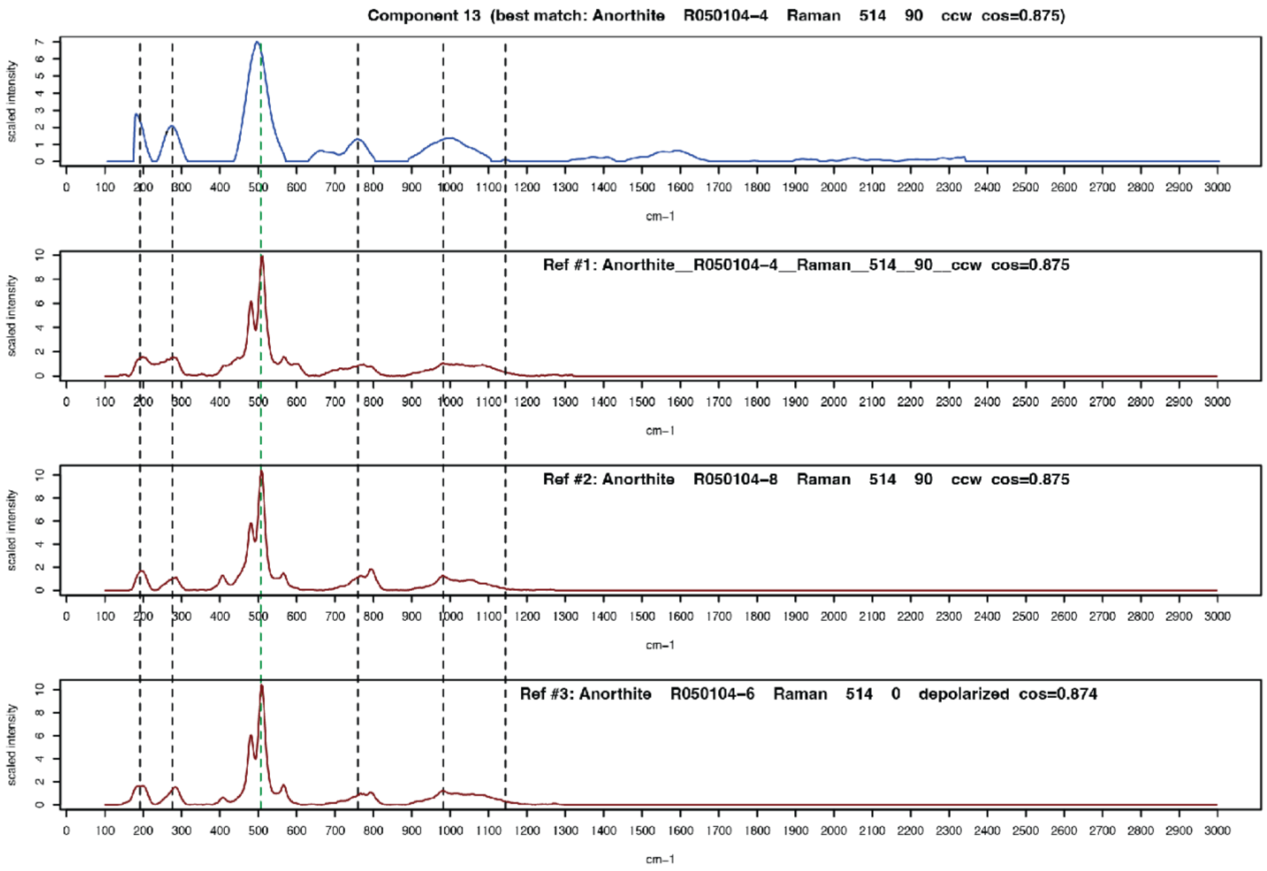
Component 13 (best match: Anorthite R050104-4 Raman 514 90 ccw cos=0.875)
Ref #1: Anorthite__R050104-4__Raman__514__90__ccw cos=0.875
Ref #2: Anorthite R050104-8 Raman 514 90 ccw cos=0.875
Ref #3: Anorthite R050104-6 Raman 514 0 depolarized cos=0.874
scaled intensity
cm-1

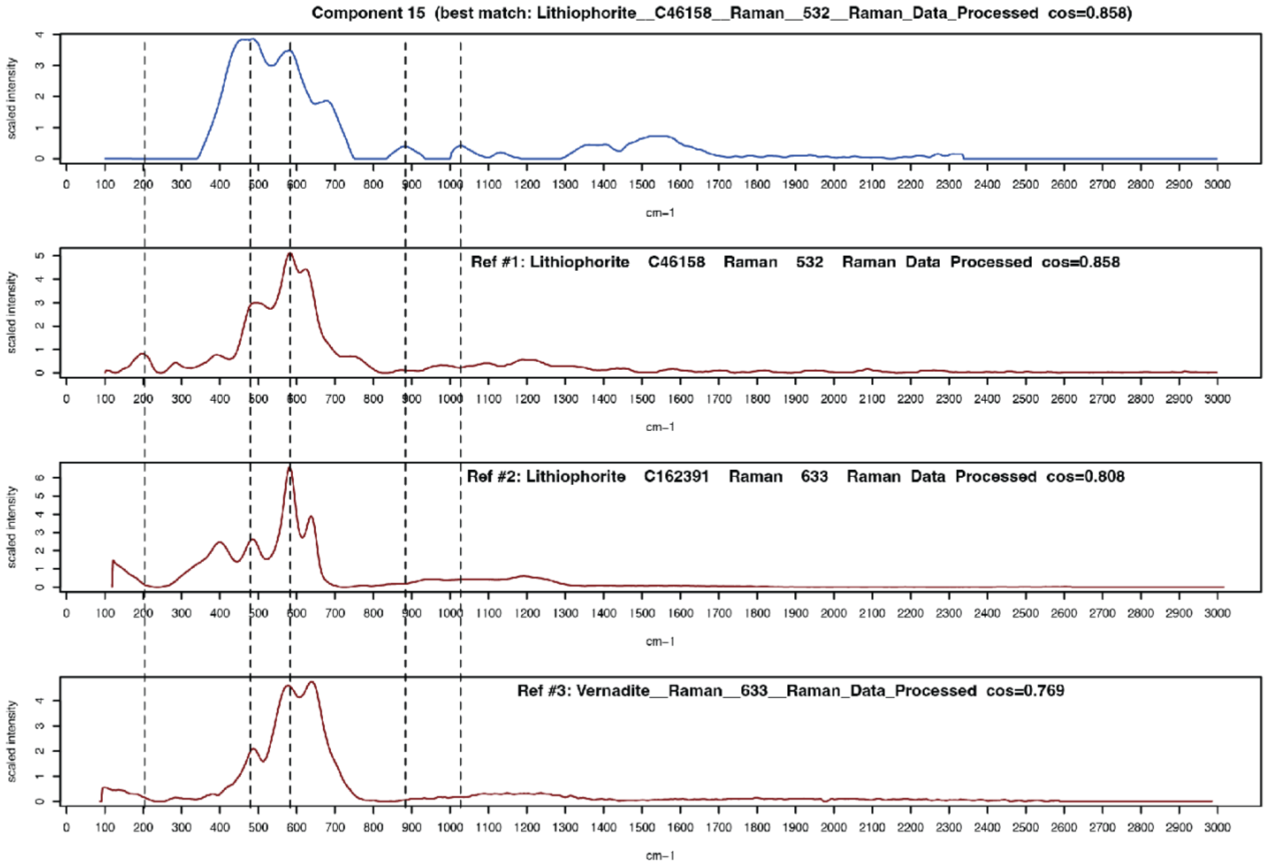
Component 15 (best match: Lithiophorite__C46158__Raman__532__Raman_Data_Processed cos=0.858)
Ref #1: Lithiophorite C46158 Raman 532 Raman Data Processed cos=0.858
Ref #2: Lithiophorite C162391 Raman 633 Raman Data Processed cos=0.808
Ref #3: Vernadite__Raman__633__Raman_Data_Processed cos=0.769
scaled intensity
cm-1

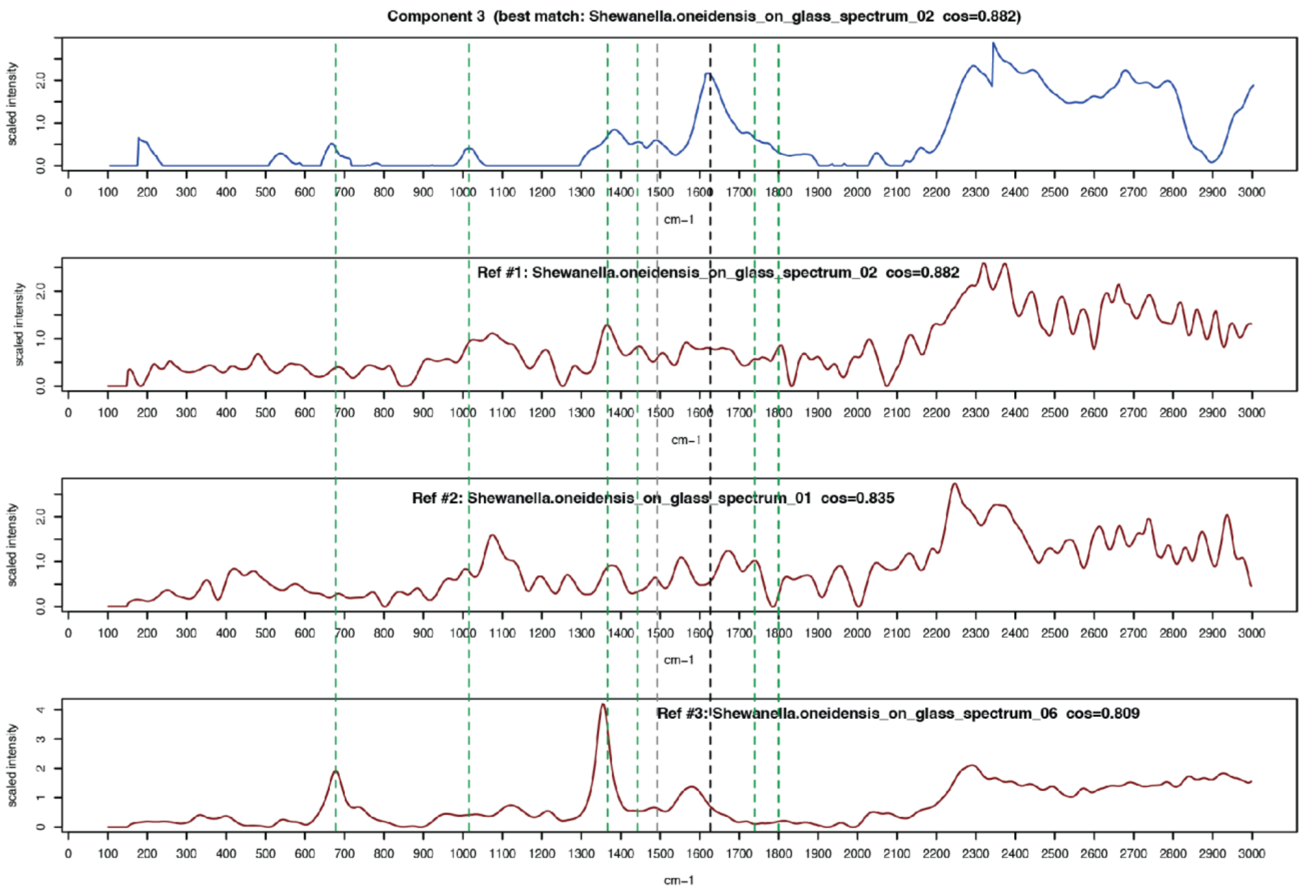

Component 3 (best match: Shewanella.oneidensis_on_glass_spectrum_02 cos=0.882)
Ref #1: Shewanella.oneidensis_on_glass_spectrum_02 cos=0.882
Ref #2: Shewanella.oneidensis_on_glass_spectrum_01 cos=0.835
Ref #3: Shewanella.oneidensis_on_glass_spectrum_06 cos=0.809
scaled intensity
cm−1

#### 2. Mineral and biomass components identified in the three mineralogically distinct samples.

Following spectral decomposition via NMF, nine components were identified for each sample (Table S1).

Table S1: The nine components for each sample, with the identity of the best spectral database match, that match's mineral class, and the percentage of pixels for which the component was the top match.

| Sample | Component # | Best Match | Mineral class (Dana Classification) | Percentage of pixels for which the component is the top match (n in parentheses) |
|---|---|---|---|---|
| SC | 15 | S. oneidensis | Biomass | 23 (1604) |
| SC | 14 | S. oneidensis (kerogen) | Biomass / kerogen | 16.5 (1150) |
| SC | 9 | Aragonite | Carbonate | 16.1 (1121) |
| SC | 4 | Vesuvianite | Sorosilicate | 14.5 (1009) |
| SC | 5 | LRW Resin | Resin | 10 (699) |
| SC | 11 | Kutnohorite | Carbonate | 9.3 (645) |
| SC | 1 | Stibnite | Sulfide | 4.8 (335) |
| SC | 10 | Feitknechtite | Oxide / Hydroxide | 3.7 (260) |
| SC | 2 | Phillipsite | Tektosilicate | 2.1 (147) |
| PN | 2 | Lithiophorite | Oxide / Hydroxide | 28.1 (2558) |
| PN | 5 | Jacobsite | Oxide | 23.7 (2151) |
| PN | 5 | Jacobsite | Oxide | 23.7 (2151) |
| PN | 14 | S. oneidensis | Biomass | 15.5 (1413) |
| PN | 9 | LRW Resin | Resin | 11 (1003) |
| PN | 8 | S. oneidensis | Biomass / kerogen | 9 (816) |
| PN | 11 | Plattnerite | Oxide | 7.6 (694) |
| PN | 6 | Lithiophorite | Oxide / Hydroxide | 2.8 (259) |
| PN | 15 | Opal | Tektosilicate | 1.3 (117) |
| PN | 7 | Annite | Phyllosilicate | 0.9 (83) |
| VB | 12 | LRW Resin | Resin | 39 (3612) |
| VB | 8 | S. oneidensis (kerogen) | Biomass / kerogen | 20 (1853) |
| VB | 3 | S. oneidensis | Biomass | 10.4 (960) |
| VB | 1 | Ferrosilite | Inosilicate | 8 (737) |
| VB | 15 | Lithiophorite | Oxide / Hydroxide | 6.7 (617) |
| VB | 10 | Stibnite | Sulfide | 4.8 (449) |
| VB | 9 | Vesuvianite | Sorosilicate | 4.5 (421) |
| VB | 2 | Annite | Phyllosilicate | 3.5 (325) |
| VB | 13 | Anorthite | Tektosilicate | 3.1 (291) |

Below, we discuss the environmental relevance of each component and its encompassing mineral class to further contextualize and verify the computational identifications.

##### 2.1. Seep Carbonate Components

In SC, the most abundant mineralogical component was aragonite (Table S1), which has been linked with periods of high seepage activity [1] and was previously detected in the Point Dume system [2] as well as many other cold seeps from different geological settings [3]. Mn-rich carbonate phases, such as kutnohorite, can form in methane seep systems involving deeply sourced methane and variable pore-water chemistry [4,5]. Kutnohorite has been also found within methane-derived authigenic carbonates in the Laptev Sea, indicating Mn-enriched pore waters and evolving fluid chemistry during geochemically distinct stages in methane seepage [6]. Sulfide minerals are commonly found in seep carbonates, given the elevated sulfide concentrations resulting from microbial sulfate reduction [7,8]. Pyrite ($FeS_2$) is the most commonly reported sulfide mineral at seeps, though sulfide can also play an important role in trace element sequestration [9,10], leading to additional types of minerals [11]. Stibnite ($Sb_2S_3$) has been detected at hydrothermally influenced hydrocarbon seeps [12], where its occurrence reflects precipitation from low-temperature, magmatically influenced fluids. Trace-metal pyritization, a process whereby heavy elements are incorporated into pyrite during early diagenesis, has been reported for antimony at cold seeps, as well as other metalloids such as As [11]. Nonetheless, it is unlikely that stibnite is actually the primary component of the SC sample; rather, the broad peaks of this component may represent a composite of sulfide minerals. Additional top hits for this component's spectrum were also sulfide minerals (Data Set 1), and we interpret this component more generally as a "sulfide-class" mineral throughout this study.

Feitknechtite is a manganese oxyhydroxide mineral that was identified as another component of our SC sample (Table S1). It forms from $Mn^{2+}$-rich fluids under oxidizing conditions, or through the alteration of other Mn minerals at low temperature [13]; for example, pyrochroite ($Mn(OH)_2$) passes through a feitknechtite transitional stage before stabilizing as minerals including manganite or hausmannite [14,15]. While feitknechtite had the top cosine similarity score for this spectral component, the top five matches were all members of the broader Mn oxyhydroxide mineral class (with a range of additional metal cations represented; Data Set 1). Mn oxyhydroxides have been implicated as a molybdenum shuttle at methane seeps [16], and Mn oxides more generally have been documented in methane-rich continental margin and cold seep environments and can be used in AOM as alternative electron acceptors to sulfate [17].

Two components of the SC spectral catalog were identified as silicate minerals (Table S1, Data Set 1). For component 2, the top match was Phillipsite-Ca and all of the other top ten matches belong to the tectosilicates, which have been documented in deep-sea sedimentary environments [18]. For component 4, the top five cosine similarity scores identified vesuvianite, a sorosilicate mineral. These identifications reflect a diversity of silicates in the SC sample. Carbonates at seeps often incorporate detrital sediment silicate grains [2,19], particularly when seep sites are close to shore [20]. Microbial consortia of Anaerobic Methanotrophic Archaea (ANME) and Sulfate-Reducing Bacteria (SRB) can also induce the precipitation of amorphous, silica-rich phases with clay-like morphology both in the natural environment [21] and long-term laboratory enrichments [22]. These detrital and authigenic constituents may be shaped by local diagenesis into silicates such as Phillipsite-Ca, which forms under alkaline pH condition in subsurface pore water [23].

##### 2.2. Polymetallic Nodule Components

In PN, iron oxyhydroxides and manganese oxides have positive and negative surface charges, respectively, that attract and enrich metal carbonate anions and Cu, Ni, and Co cations [24,25]. An early mineralogical survey included both lithiophorite and jacobsite, the top matches of the most abundant components in our

PN sample (Table S1) as putative phases in deep-sea nodules occurring in these samples [26]. Lithiophorite is a lithium-bearing manganese oxide, and it is compositionally related to asbolane, which also showed a strong match to this Raman component (Data Set 1) and has been reported in polymetallic nodules [27]. These two minerals can exhibit mixed compositions, referred to as asbolane–lithiophorite intermediates, which are challenging to distinguish due to their structural complexity and poor crystallinity [28]. Jacobsite is a $Mn^{2+}$-$Fe^{3+}$ oxide mineral that was initially discovered in terrestrial Mn ore deposits but also occurs in deep-sea nodules [27]. The top several matches for the most prevalent components in sample PN are consistent within mineral subclasses, indicating that the nodule likely contains substantial quantities of a Mn-Fe oxide (component 5) and a Mn hydroxide doped with an array of metal cations (components 2 and 6; Table S1, Data Set 1). In addition, the high cosine similarity score between component 11 and plattnerite is consistent with the detection of trace amounts of lead and other heavy metal oxides in ferromanganese nodules [24,27,29].

CCZ sediments contain silica in the form of biogenic opal, which is derived mostly from diatomaceous and other siliceous microfossils that are recrystallized in benthic sediments around and within nodules [30], and can reach up to 30-60% of the sediment composition in some cases [31]. Given previous observations that opal is associated with iron-bearing smectite clays in CCZ sediments [31] and nodules [32], and other nontronite-like iron-rich smectite, combined with the assignment of annite as the best match to one of our PN sample's components 7, it is possible that smectite clay minerals were misclassified as annite, as discussed below for the VB samples.

#### 2.3. Volcanic Basalt Components

In VB, the most abundant minerals were silicates (Table S1). The inosilicate ferrosilite, an iron-rich endmember of pyroxene minerals, was reported in a previous mineralogical study of Hvalfjordur basalts of the Reykjanes geothermal system [33]. Metal hydroxides, including Mn-containing minerals such as lithiophorite, can form from degassing lavas at Fagradalsfjall [34]. However, most MnO manifests as a minor component within ferromagnesian minerals such as olivine, and lower concentrations occur in basaltic glass, highlighting patterns of Mn partitioning during crystallization of tholeiitic magma typical of Fagradalsfjall [35]. Although Raman can typically distinguish pure phases of MnO-type oxides from Mn hydroxides, overlapping Mn–O vibrational bands in complex natural samples can limit spectral precision [36]. Nonetheless, the top six matches to component 15 were all metal hydroxide minerals (Data Set 1), providing confidence that this level of mineralogical specificity accurately captures this mineral constituent. Sulfide minerals such as stibnite can form through hydrothermal fluid interactions with basaltic fractures and seawater-derived fluids at depth in the Reykjanes geothermal system [37,38]. Trace element studies reveal that metalloids such as Sb (together with Pb, Ag, and As) are enriched during basalt alteration and partition into sulfide precipitates during boiling of hydrothermal fluids [37,38]. Stibnite was detected in a terrestrial mud volcano, associated with strong fluid-mediated transport of metals [39], and co-occurred with pyrite at Steamboat Springs in Nevada, a process that was linked to geothermal and volcanic activity [40].

Annite is the iron-rich endmember of biotite, a phyllosilicate mineral within the mica group, which was found in a subsurface hydrothermal system on the Reykjanes Peninsula as a result of reactions between hot brines and basaltic rock [41]. While Annite was the top match for component 2, we cannot rule out other phyllosilicates, such as smectite and chlorite, which are produced through low-temperature hydrothermal alteration at Fagradalsfjall [42]. It is often challenging to distinguish specific phyllosilicates with Raman spectroscopy because similar layered structures produce overlapping low-frequency absorption bands, and variable Fe–Mg substitutions can shift peak positions and increase spectral similarity [43]. Given the basaltic composition of the Fagradalsfjall lavas, Mg-rich smectites such as saponite are expected alteration products in Icelandic hydrothermal and weathering environments, which is not present in the RRUFF database, precluding its identification in our workflow. Anorthite-rich plagioclase (within the tectosilicate mineral class) is commonly observed in Fagradalsfjall basalts [44], where plagioclase phenocrysts range from ~77-80% anorthite in rims to 87-89% in mineral cores and constitute ~2–15% of the lava by volume [45].

**2.4. Additional Spectral Decomposition Considerations**

While the vast majority of identified components were consistent with mineralogical studies of their respective samples, not all of them were previously noted as substantial contributors to the mineral assemblage. Discrepancies in the component IDs between computationally determined mineral species and previously detected minerals in the three rock systems are likely due to well-established issues with Raman spectra databases, such as non-standardized equipment and acquisition parameters; variable spectral resolutions of contributed spectra; inconsistent spectral processing, leading, in some cases, to high noise; and spectral variation based on crystal orientation [46-48]. We stress, however, that even in cases where the specific mineral is not traditionally associated with the analyzed rock, the more generalized "class" of mineral is, and this is the level at which we interpret the results.

Our computational approach to spectral decomposition offers an opportunity to address some of the challenges that limit the throughput and widespread use of Raman microspectroscopy in geomicrobiological studies. We note that the ultimate applicability of the results is dependent upon the quality of the Raman spectra and the spectral library being used. While we did supplement the RRUFF database with biological and resin standards relevant to our workflow and microbial focus, generating a fully customized database with pure and mixed phases from all three sample types was beyond the scope of this study. Doing so may allow future studies to interpret mineral components with a greater degree of specificity rather than using mineral class-level assignments.

### 3. Correlative approach

The presence of biomass was verified using confocal microscopy after DNA staining with Sybr Green (see methods in the Main Text). Biomass was observed in each Field of View (FOV) of all three rock samples using fluorescence microscopy and then the same FOV as analyzed with Raman spectroscopy.

Fig. S4: Representative fields of view (FOVs) for each sample. The same region is shown as imaged by the Raman instrument's dark-field camera (left panels) and the confocal fluorescence microscope (right panels). Blue rectangles indicate the imaged area. The biomass component heat map is shown alongside the corresponding SYBR Green fluorescence image to illustrate the spatial correlation between the Raman-derived biomass signal and SYBR Green staining.

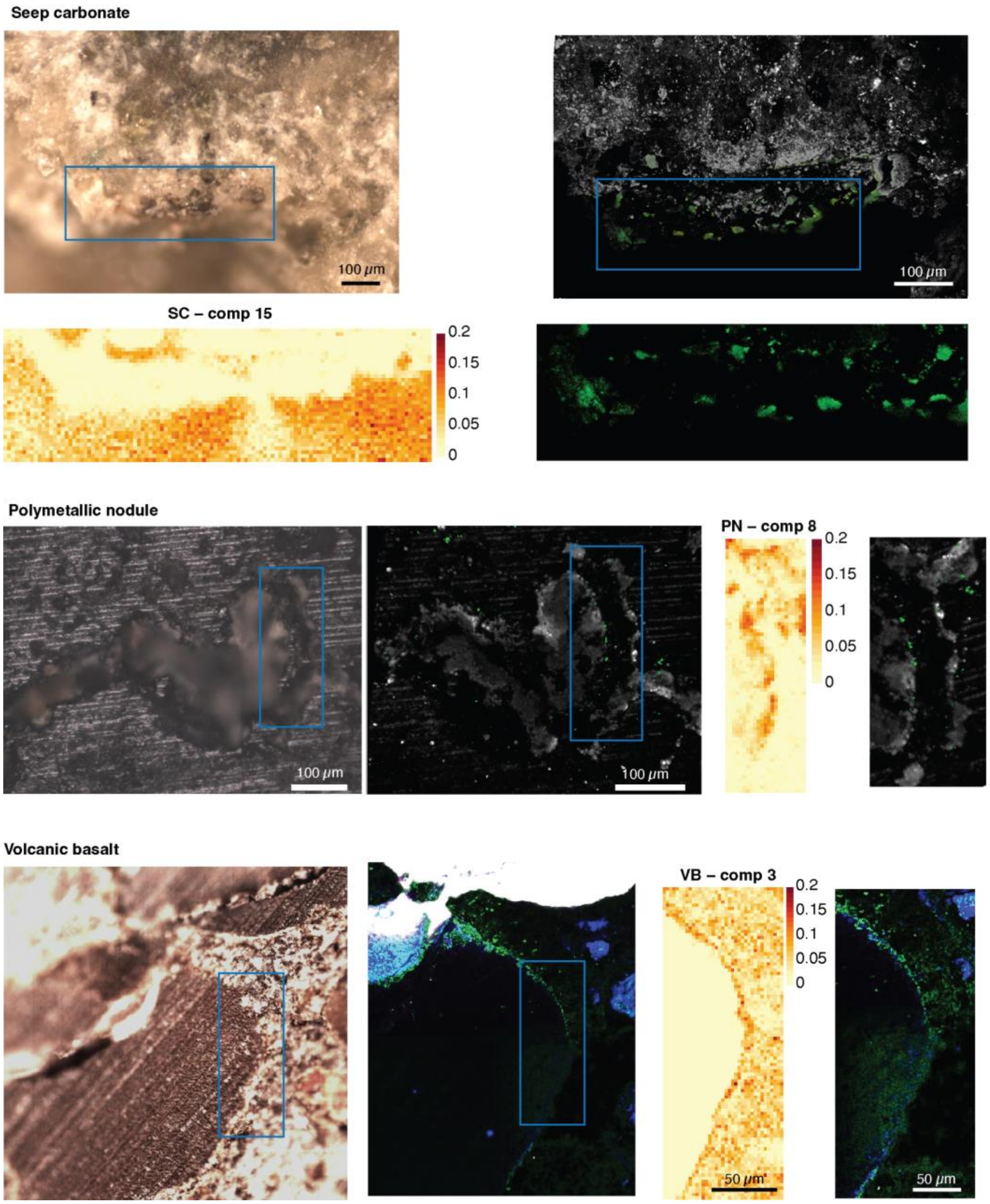

#### 4. Statistical assessment of effect size

Differences in compositional-heterogeneity indices between biomass and mineral pixels were assessed within each sample type (SC, VB, PN) separately for each index (intra-pixel: 1 – Gini and Shannon entropy; inter-pixel: the spectral heterogeneity score, and the modified Rao's quadratic entropy) and for each component condition (all components, dominant component omitted, biomass component omitted).

The Hodges–Lehmann (HL) method was used to estimate the median difference between biomass and mineral pixels across all comparison cases, providing an additional robust, nonparametric measure of effect size. This approach was used to quantify the direction and magnitude of the between-group difference and to assess the overall trend in the comparison without assuming a specific data distribution. Unlike a p-value, this approach quantifies the magnitude and direction of the difference. HL = 0 indicates no shift, and thus no difference between the groups. HL > 0 indicates that biomass pixels tend to be higher than mineral pixels (by approximately the HL estimate number), while HL < 0 indicates that biomass pixels tend to be lower than the mineral pixels (by approximately the absolute value of the HL estimate).

HL estimators calculate the median of all pairwise differences between observations (within-pair differences between biomass and mineral pixels), rather than the difference between the two sample medians, thus quantifying both the magnitude and direction of the difference. For our samples, the effect sizes measured as HL are small to moderate throughout the comparisons, though the effect direction across the three geologically distinct substrates and across metrics were consistent (Data Set 2). For the all-components condition, 11 of 12 biomass vs. mineral contrasts point in the hypothesized direction — biomass pixels are internally more homogeneous (1 − Gini and Shannon $\delta < 0$) and, by Rao's Q, sit in more heterogeneous surroundings ($\delta > 0$) — across all three samples. The single exception is the SC spectral heterogeneity score (1 − SHS).

Cliff's delta values and directions are provided in Fig. S5 below. By our sign convention $\delta > 0$ indicates that biomass pixels tend to exceed mineral pixels in terms of the heterogeneity metric.

Fig. S5. Cliff's delta directions across all tested conditions.

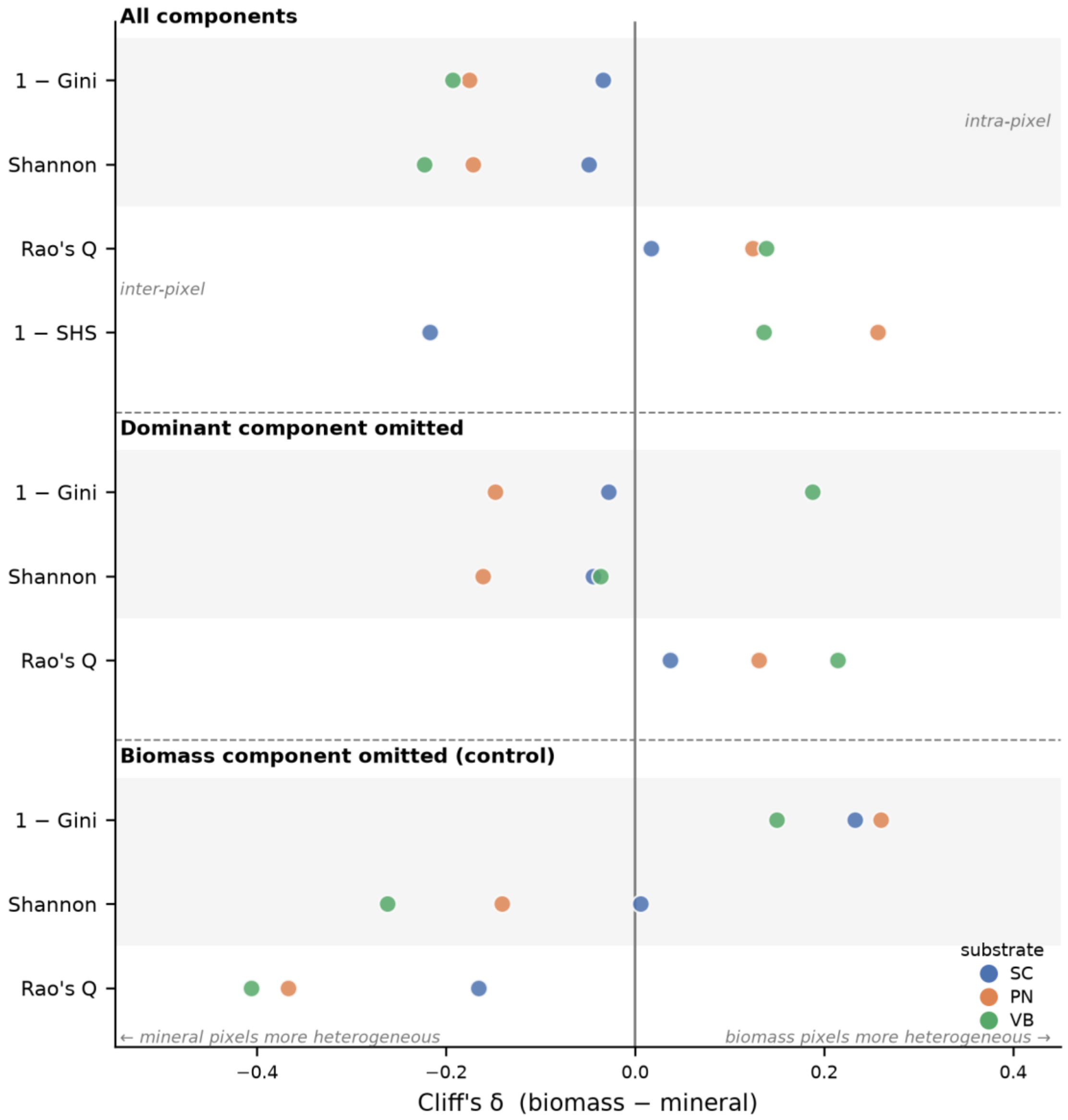

All components
1 − Gini
Shannon
Rao's Q
1 − SHS
intra-pixel
inter-pixel
Dominant component omitted
1 − Gini
Shannon
Rao's Q
Biomass component omitted (control)
1 − Gini
Shannon
Rao's Q
substrate
SC
PN
VB
← mineral pixels more heterogeneous
biomass pixels more heterogeneous →
−0.4
−0.2
0.0
0.2
0.4
Cliff's δ (biomass − mineral)

#### 5. Calculation of the area analyzed for each sample

FIJI (Image J2 - Version 2.16.0/1.54p) was used to estimate the area of each mineral section. A Region of Interest (RoI) was drawn around the rock part of the section using the “Polygon selection” and “Wand tracing” tools, and the “Measure” tool was used to quantify the area within the ROI. This calculation included the entire surface area of the mineral section, including void space embedded by resin and minor cracks within the area, and was intended to approximate the surface area that was analyzed with our microscopy workflow.

Fig. S6: Seep Carbonate rock section embedded in resin. The yellow line encompasses the rock section that was analyzed. The external perimeter of the disc is composed of resin.

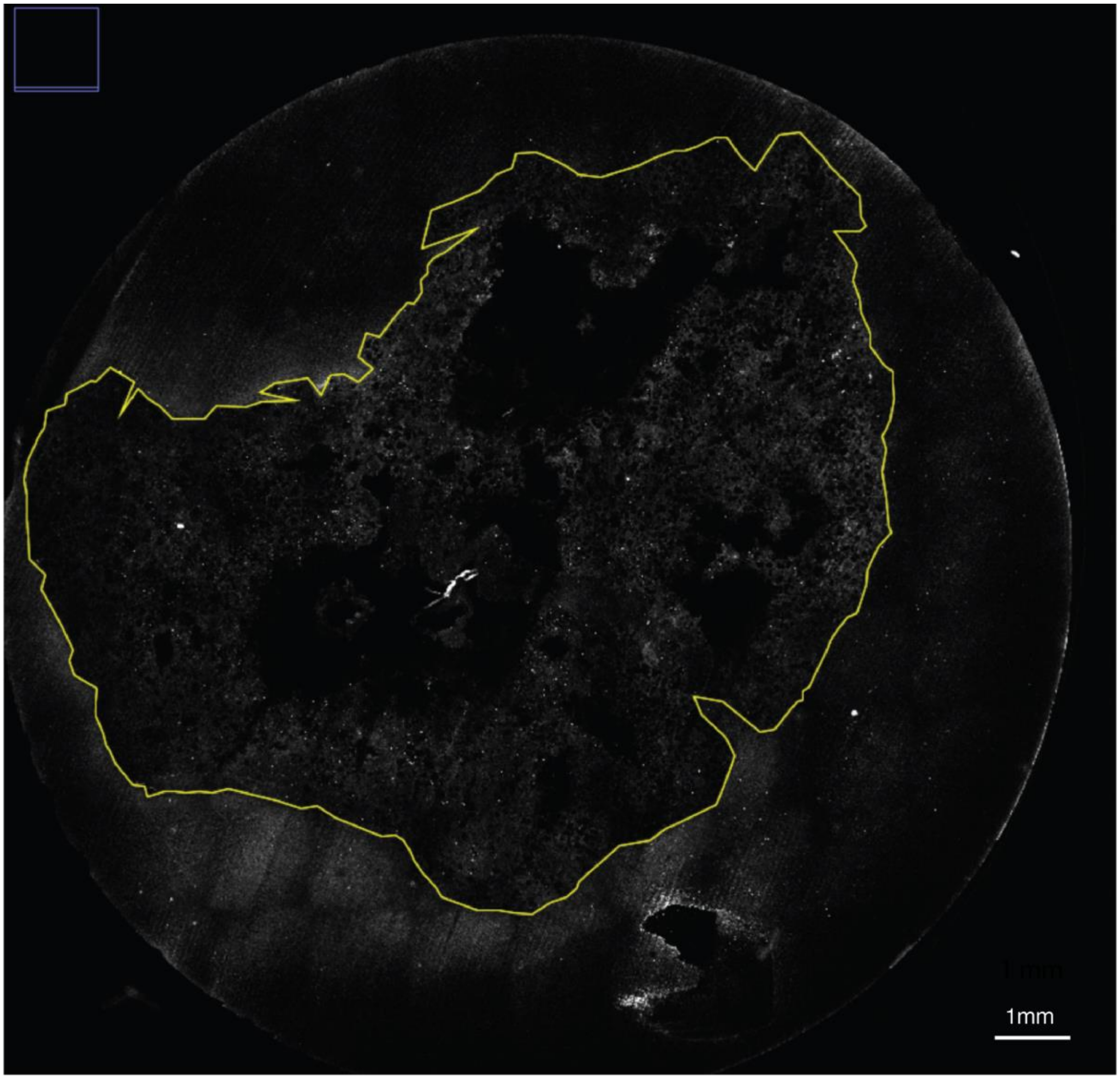

Fig. S7: Volcanic Basalt rock section embedded in resin. The black area is the rock section that was analyzed. The external perimeter of the disc is composed of resin.

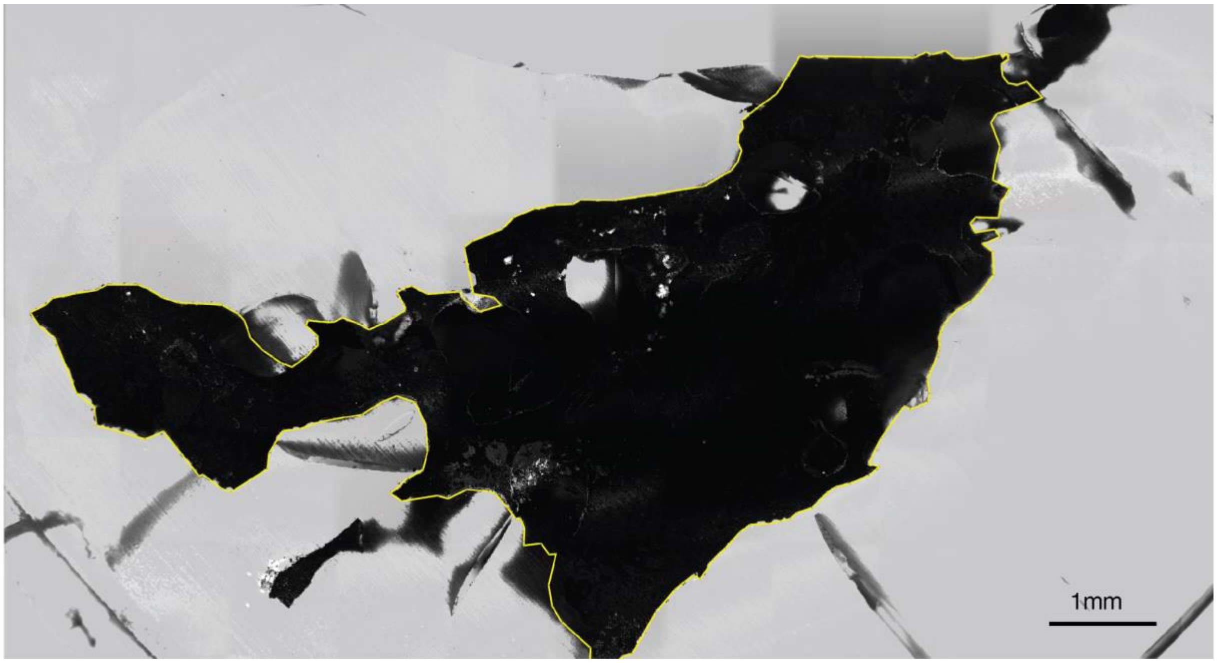

Fig. S8: Polymetallic Nodule rock section embedded in resin. The yellow line encompasses the rock section that was analyzed. The external perimeter of the disc is composed of resin.

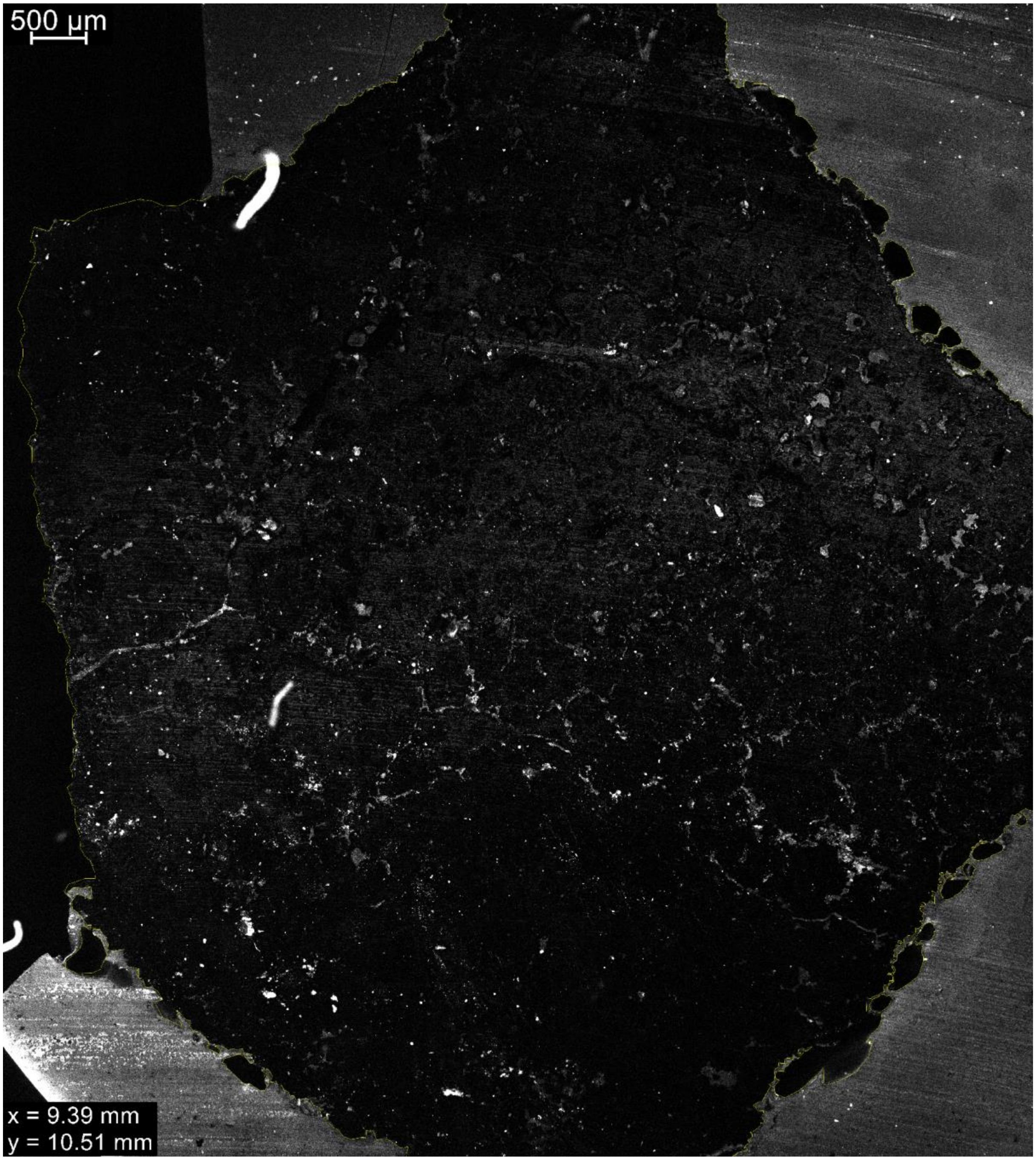

#### 6. Evaluating the effects of spatial interdependence

To determine if our results are independent of the influence of spatial proximity, we tested the effect of repeatedly and intensively thinning the data for each FOV, using approximately one pixel per spatial-decorrelation block (Table S2). (The block size was set from a per-FOV empirical variogram; 1,000 random thinnings). A variogram describes how similar observations are as a function of the distance between them. Under this thinning (1,000 random draws) the intra-pixel contrasts (Gini, Shannon) retain the full-data effect size direction in the large majority (≈88–99%) of draws across all three samples (i.e., in one sample, 88-99% of the 1,000 thinned datasets gave the same effect direction as the full dataset). For the inter-pixel metric Rao's Q, ≈73–94% of draws resulted in the same effect size direction. These results indicate that the observed effects are not simply an artifact of spatial autocorrelation or oversampling of neighbor pixels, but rather that the reported effects are qualitatively consistent after accounting for spatial dependence through thinning.

**Table S2. Summary of the spatial interdependence analysis following spatial thinning.**
Spatial thinning was performed to evaluate the robustness of the estimated differences in heterogeneity indices accounting for spatial interdependence between the biomass and mineral pixels. We computed the same analysis for each *sample, each metric* (heterogeneity indices*)* and *each condition* investigated in this study. *delta_raw* is the difference in the spatial interdependence metric between the bio and min pixel calculated from the original (unthinned) data. *thin_n_bio_med* and *thin_n_min_med* are the median numbers of observations retained in the bio and min pixels across all thinning iterations. *thin_delta*_med is the median difference in the metric across thinning iterations. *thin_delta_low* and *thin_delta_high* represent the lower and upper bounds of the empirical interval of the delta (Δ) values obtained after thinning. *frac_same_dir* is the proportion of thinning iterations in which the sign of Δ was the same as in the original analysis, indicating the robustness of the direction of the observed difference. The biomass pixels analyzed were 2754, 2229, and 2813 while the mineral pixels were 3207, 5862, and 2840 for SC, PN, and VB, respectively.

| sample | metric | condition | delta_raw | thin_n_bio_med | thin_n_min_med | thin_delta_med | thin_delta_low | thin_delta_high | frac_same_dir |
|---|---|---|---|---|---|---|---|---|---|
| SC | 1-Gini | all components | -0.034 | 14 | 39 | -0.208 | -0.533 | 0.146 | 0.881 |
| SC | 1-Gini | biomass omitted | 0.233 | 14 | 39 | -0.027 | -0.390 | 0.346 | 0.437 |
| SC | 1-Gini | dominant omitted | -0.028 | 14 | 39 | -0.206 | -0.524 | 0.134 | 0.869 |
| SC | Shannon | all components | -0.049 | 16 | 42 | -0.217 | -0.509 | 0.111 | 0.895 |
| SC | Shannon | biomass omitted | 0.006 | 16 | 42 | -0.219 | -0.534 | 0.080 | 0.082 |
| SC | Shannon | dominant omitted | -0.044 | 16 | 42 | -0.234 | -0.535 | 0.080 | 0.929 |
| SC | SHS(1-SHS | all components | -0.217 | 9 | 14 | -0.424 | -0.767 | -0.030 | 0.980 |
| SC | RaoQ | all components | 0.017 | 21 | 128 | 0.081 | -0.187 | 0.306 | 0.725 |
| SC | RaoQ | biomass omitted | -0.165 | 21 | 128 | -0.118 | -0.333 | 0.136 | 0.836 |
| SC | RaoQ | dominant omitted | 0.037 | 21 | 128 | -0.086 | -0.349 | 0.154 | 0.248 |
| PN | 1-Gini | all components | -0.175 | 14 | 20 | -0.417 | -0.729 | -0.068 | 0.991 |
| PN | 1-Gini | biomass omitted | 0.260 | 14 | 20 | 0.157 | -0.194 | 0.493 | 0.795 |
| PN | 1-Gini | dominant omitted | -0.148 | 14 | 20 | -0.319 | -0.653 | 0.122 | 0.925 |
| PN | Shannon | all components | -0.171 | 12 | 20 | -0.359 | -0.676 | 0.024 | 0.967 |
| PN | Shannon | biomass omitted | -0.141 | 12 | 20 | -0.317 | -0.655 | 0.091 | 0.941 |
| PN | Shannon | dominant omitted | -0.161 | 12 | 20 | -0.341 | -0.671 | 0.053 | 0.960 |
| PN | SHS(1-SHS | all components | 0.257 | 9 | 9 | 0.025 | -0.400 | 0.429 | 0.532 |
| PN | RaoQ | all components | 0.125 | 17 | 36 | 0.157 | -0.140 | 0.484 | 0.838 |
| PN | RaoQ | biomass omitted | -0.367 | 17 | 36 | -0.317 | -0.576 | -0.025 | 0.983 |
| PN | RaoQ | dominant omitted | 0.131 | 17 | 36 | 0.209 | -0.108 | 0.487 | 0.908 |
| VB | 1-Gini | all components | -0.193 | 40 | 42 | -0.179 | -0.403 | 0.056 | 0.932 |
| VB | 1-Gini | biomass omitted | 0.150 | 40 | 42 | 0.147 | -0.098 | 0.373 | 0.882 |
| VB | 1-Gini | dominant omitted | 0.188 | 40 | 42 | 0.287 | 0.067 | 0.510 | 0.991 |
| VB | Shannon | all components | -0.223 | 37 | 37 | -0.273 | -0.511 | -0.041 | 0.990 |
| VB | Shannon | biomass omitted | -0.262 | 37 | 37 | -0.375 | -0.595 | -0.156 | 1.000 |
| VB | Shannon | dominant omitted | -0.037 | 37 | 37 | -0.006 | -0.253 | 0.262 | 0.517 |
| VB | SHS(1-SHS | all components | 0.136 | 35 | 17 | 0.014 | -0.261 | 0.304 | 0.544 |
| VB | RaoQ | all components | 0.139 | 70 | 65 | 0.135 | -0.034 | 0.304 | 0.942 |
| VB | RaoQ | biomass omitted | -0.406 | 70 | 65 | -0.487 | -0.618 | -0.357 | 1.000 |
| VB | RaoQ | dominant omitted | 0.214 | 70 | 65 | 0.208 | 0.056 | 0.369 | 0.995 |

#### 7. Raman imaging parameters for each sample and field of view

Table S3: Pixel size and resolution for each field of view analyzed with Raman microspectroscopy in this study. The area of each FOV is provided in square millimeters ($mm^2$).

| **Sample** | **Resolution (µm)** | **Number of pixels** | **rows x columns** | **Areas in $mm^2$** |
|---|---|---|---|---|
| Seep Carbonate (SC) | | | | |
| FOV-1 | 5 | 3920 | 35 x 112 | 0.0980 |
| FOV-2 | 4 | 805 | 35 x 23 | 0.0129 |
| FOV-3 | 4 | 1846 | 26 x 71 | 0.0295 |
| FOV-4 | 2 | 399 | 19 x 21 | 0.0016 |
| *SC Sum* | | *6970* | | *0.1420* |
| Polymetallic nodule (PN) | | | | |
| FOV-1 | 3 | 1435 | 41 x 35 | 0.0129 |
| FOV-2 | 1 | 5772 | 78 x 74 | 0.0058 |
| FOV-3 | 2 | 208 | 26 x 8 | 0.0008 |
| FOV-4 | 3 | 1679 | 73 x 23 | 0.0151 |
| *PN Sum* | | *9094* | | *0.0346* |
| Volcanic basalt (VB) | | | | |
| FOV-1 | 7 | 1386 | 66 x 21 | 0.0679 |
| FOV-2 | 5 | 3552 | 96 x 37 | 0.0888 |
| FOV-3 | 5 | 2967 | 69 x 43 | 0.0742 |
| FOV-4 | 2 | 1360 | 34 x 40 | 0.0054 |
| *VB Sum* | | *9265* | | *0.2363* |

#### 8. Additional information on heterogeneity metrics

To quantify heterogeneity, we selected metrics that are appropriate for compositional data, namely the Shannon index, the Gini index, and Rao's quadratic entropy, as they capture complementary properties of composition, evenness, and pairwise dissimilarity within pixels.

The Gini coefficient was calculated to measure intra-pixel heterogeneity. The Lorenz curve (Fig. S9) was calculated for each pixel, plotting the cumulative percentage of component weights against the cumulative percentage of components. Gini values were computed according to equation A / (A+B), where A is the area between the line of perfect equality and the Lorenz curve, and B is the area between the Lorenz curve and the perpendicular lines bounding it.

Fig. S9: A graphical schematic of how the Gini coefficient is calculated.

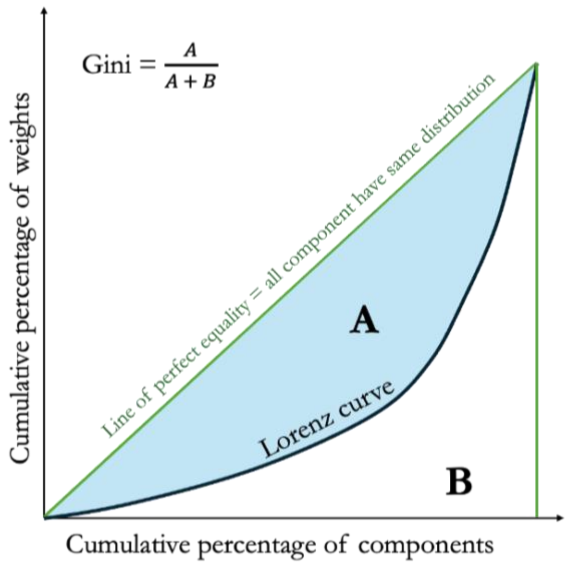
